# DNA Punch Cards: Storing Data on Native DNA Sequences via Nicking

**DOI:** 10.1101/672394

**Authors:** S Kasra Tabatabaei, Boya Wang, Nagendra Bala Murali Athreya, Behnam Enghiad, Alvaro Gonzalo Hernandez, Christopher J. Fields, Jean-Pierre Leburton, David Soloveichik, Huimin Zhao, Olgica Milenkovic

## Abstract

Synthetic DNA-based data storage systems have received significant attention due to the promise of ultrahigh storage density and long-term stability. However, all platforms proposed so far suffer from high cost, read-write latency and error-rates that render them noncompetitive with modern optical and magnetic storage devices. One means to avoid synthesizing DNA and to reduce the system error-rates is to use readily available native DNA. As the symbol/nucleotide content of native DNA is fixed, one may adopt an alternative recording strategy that modifies the DNA topology to encode desired information. Here, we report the first macromolecular storage paradigm in which data is written in the form of “nicks (punches)” at predetermined positions on the sugar-phosphate backbone of native dsDNA. The platform accommodates parallel nicking on multiple “orthogonal” genomic DNA fragments and paired nicking and disassociation for creating “toehold” regions that enable single-bit random access and strand displacement in-memory computations. As a proof of concept, we used the programmable restriction enzyme *Pyrococcus furiosus* Argonaute to punch two files into the PCR products of *Escherichia coli* genomic DNA. The encoded data is accurately reconstructed through high-throughput sequencing and read alignment.

## Introduction

All existing DNA-based data recording architectures store user content in synthetic DNA oligos (1-12) and retrieve desired information via next-generation (NGS; HiSeq and MiSeq) or third generation nanopore sequencing technologies (6). Although DNA sequencing can be performed routinely and at low cost, *de novo* synthesis of DNA strands with a predetermined nucleotide content is a major bottleneck due to multiple issues (13). First, DNA synthesis protocols add one nucleotide per cycle, with each cycle taking seconds, and therefore are inherently slow and prohibitively expensive compared to existing optical and magnetic writing mechanisms. Second, DNA synthesis is an error-prone procedure that often fails to account for a large volume of information-bearing oligos and which leads to oligo pools that contain close to 1% of substitution and indel errors (14). Third, the available mass per synthetic DNA oligo is usually small, enabling only a limited number of readouts (9).

To address the above limitations of DNA-based data storage systems and pave the way towards future low-cost molecular storage solutions we propose a new storage paradigm that represents information via *in vitro* topological modifications on native DNA (e.g., genomic DNA, cloned or PCR-amplified products). Unlike all previously proposed methods for DNA-based data storage, our system stores data in the sugar-phosphate backbone of DNA molecules rather than their sequence content. More precisely, binary information-bearing strings are converted into positional encodings that describe if a carefully preselected set of nicking sites is to be nicked or not. The information stored in nicks can be retrieved in an error-free manner using NGS technologies, similar to synthesis-based approaches. This is accomplished through alignment of DNA fragments obtained through the nicking process to the known reference genomic DNA strands. Due to the presence of the reference even very small fragment coverages lead to error-free readouts, which is an important feature of the system. Alternative readout approaches, such as non-destructive solid-state nanopore sequencing, can be used instead, provided that further advancement of the related technologies enable high readout precision.

Nick-based storage also allows for introducing a number of additional functionalities into the storage system, such as bitwise random access and pooling – both reported in this work – and in-memory computing solutions reported in two follow-up papers (15,16). These features come at the cost of reduced storage density which is roughly 50-fold smaller than that of synthetic DNA-based platforms.

It is important to point out that although there are many different means for topologically modifying DNA molecules, for data storage applications enzymatic nicking (i.e., creating a single-bond cut in the DNA sugar-phosphate backbone via an enzyme) appears to be the most specific, efficient and versatile recording method. DNA methylation, a well-known type of modification that may be imposed on cytosine bases, is insufficiently versatile and hard to impose in a selective manner. Topological alterations in DNA in the form of secondary structure may not be cost- or density-efficient as additional nucleotides are needed to create such structures (17).

The proposed native DNA-based storage architecture is depicted in Figure 1, encompassing a Write and Read unit. The Write unit involves extracting native DNA registers to be used as recording media, identifying proper nicking sites and encoding the data through positional codes. It also includes a parallel nicking procedure centered around highly specific controllable nicking endonucleases. The Read unit involves dsDNA denaturation, library preparation and NGS sequencing followed by read alignment and decoding.

**Figure 1.**
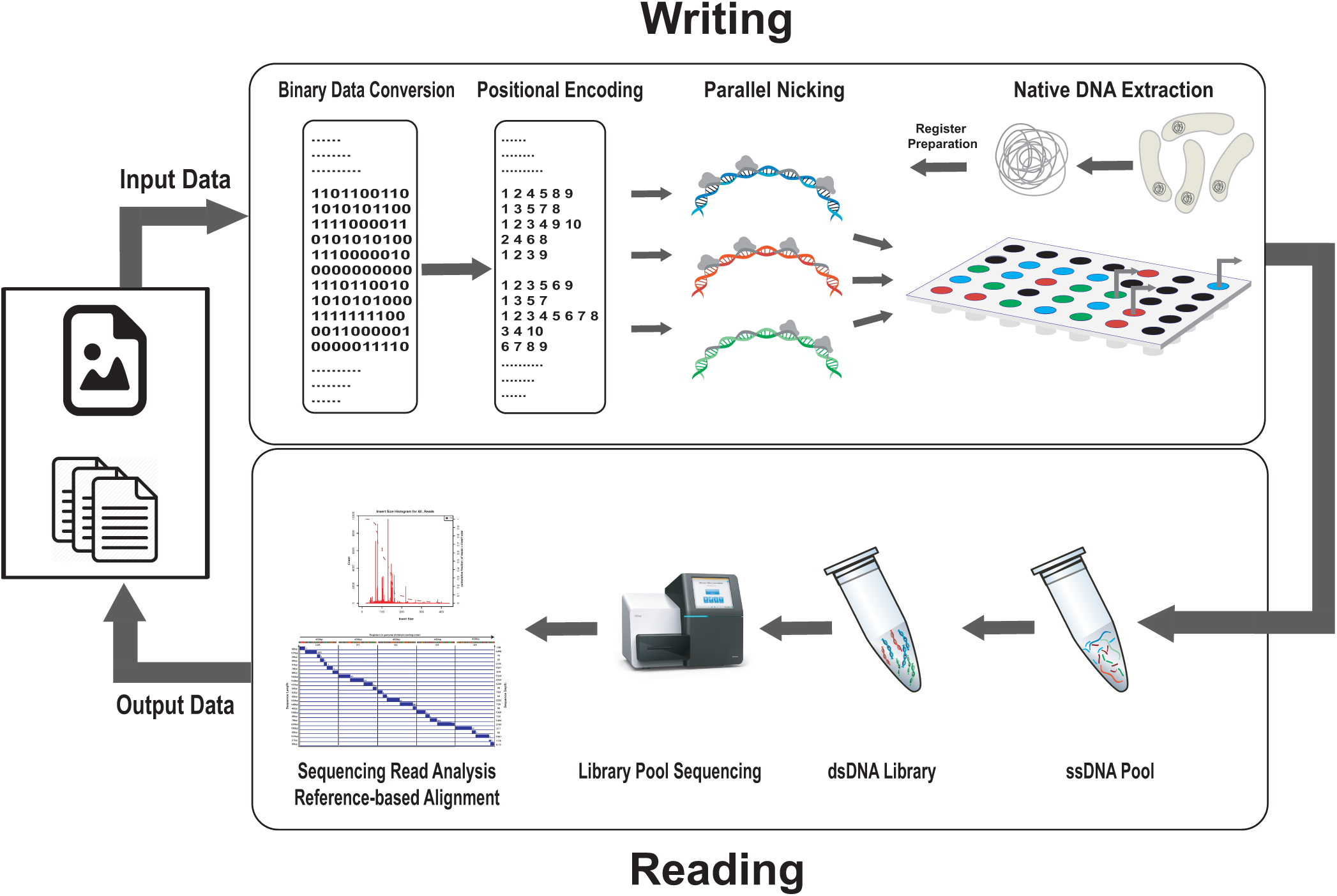
The native DNA-based data storage platform. In the **Write component**, arbitrary user content is converted into a binary message. The message is then parsed into blocks of *m* bits, where *m* corresponds to the number of nicking positions on the register (for the running example, *m* = 10). Subsequently, binary information is translated into positional information indicating where to nick. Nicking reactions are performed in parallel via combinations of *Pf*Ago and guides. In the **Read component**, nicked products are purified and denatured to obtain a pool of ssDNAs of different lengths. The pool of ssDNAs is sequenced via MiSeq. The output reads are processed by first performing reference-based alignment of the reads, and then using read coverages to determine the nicked positions.

The individual components of the Write and Read system are described in detail in the Results section.

## Results

### The Write architecture

#### The writing tools – nicking enzymes

To implement a nick-based data storage platform, one first needs to identify a nicking enzyme with optimized programmability. Until this work, this remained a challenge for reasons described in what follows.

Known nickases (natural/engineered) are only able to detect and bind specific sequences in DNA strands that tend to be highly restricted by their context. For example, nicking endonucleases only recognize specific sequences, usually 6 bps long, which makes them difficult to use for large-scale recording. Similarly, *Streptococcus pyogenes* Cas9 nickase (*Sp*Cas9n), a widely used tool for genetic engineering applications, requires the presence of a protospacer adjacent motif (PAM) sequence (NGG) at the 3’ site of the target DNA. The NGG motif constraint limits the nicking space to 1/16 of the available positions due to the GG dinucleotide in the PAM sequence. The *Sp*Cas9n complex uses RNA guides (gRNAs) to bind the target which makes it rather unstable and hard to handle. Furthermore, *Sp*Cas9n is a single turnover enzyme (18), i.e., one molecule of the enzyme can generate only one nick in a DNA molecule. These properties make *Sp*Cas9n exhibit low efficiency and versatility for storage applications.

To address these issues, we propose to use the DNA-guided programmable restriction enzyme *Pyrococcus furiosus* Argonaute (*Pf*Ago) (19) as our writing tool. *Pf*Ago is a highly accurate artificial restriction enzyme (ARE) for efficient double-stranded cleavage at arbitrary sites that generates defined sticky ends of various lengths (19). The enzyme has a significantly larger flexibility in double-stranded DNA cleaving compared to the Cas9 nickase, as it is not restricted by the presence of any special sequence at its recognition site. Most importantly, the enzyme has a high turnover rate as one enzyme molecule can be used to create a large number (∼100s) of nicks. *Pf*Ago also uses 16 nt DNA guides (gDNAs) that are stable, cheap to acquire and easy to handle *in vitro*. The *Pf*Ago enzyme has found various applications in genetic engineering and genomic studies, such as DNA assembly and DNA fingerprinting (19).

DNA cleavage requires the presence of two gDNAs, targeting both strands at positions close to each other, which is not needed nor desirable for storage applications.

We hence performed extensive experimental studies to demonstrate that under proper reaction conditions (e.g., buffer and temperature), *Pf*Ago with single gDNAs can also target only one of the DNA strands and successfully perform simultaneous nicking of multiple prescribed sites on that strand with high efficiency and precision within 40 min. An illustrative comparison of the nicking performance of *Sp*Cas9n, another potential DNA nicking enzyme, and *Pf*Ago is provided in Table S2 and Figure S3-4.

#### DNA register and nicking site selection

The genomic DNA was extracted from a culture of *E. coli* K12 MG1655 grown overnight. We found 450 bps to be a suitable length for our recording DNA fragments, henceforth referred to as registers, given that:

- This length can accommodate a sufficiently large number of nicking sites. For example, the working example register used for file storage includes 10 sites, which are separated by at least 25 bps. This distance constraint is imposed to ensure that inter-nick strands do not disassociate at room temperature and that the intra-nicking fragments are of sufficiently long length. Other length selections are possible as long as they do not lead to undesirable strand disassociations.
- As the nicking registers have to include nicking recognition sites that have large edit distance so as to avoid nonspecific nicking, one long content-predefined native DNA register may not be as versatile as a collection of shorter registers selected from different parts of the genomic sequence.
- It is a readable length for both NGS platforms and solid-state nanopores. Also, the selected length makes post-sequencing analysis and reference alignments easier and less computationally demanding.

Therefore, several 450 bp candidate registers were PCR amplified from the *E. coli* genomic DNA. Of these candidates, five registers were selected for system implementation and testing. The registers contain between five and ten nicking sites, all located on the antisense (bottom) strand (Figure S5a, S9b, c, d, e). The sequences of length 16 corresponding to the nicking sites are selected to be at a Hamming distance >8 in order to avoid nonspecific nicking. As the enzyme cuts a certain site with high accuracy (between positions 10 and 11 of the corresponding gDNA (19)), this selection eases the way towards predicting the lengths that are supposed be observed after sequencing. As a result, each register is associated with a particular set of intra-nicking fragment lengths that are expected to arise upon the completion of the nicking reactions.

#### Positional encoding

Each register contains *n* designated nicking positions. As previously described, even though long registers up to few kbs can be easily accommodated and amplified via PCR, multiple registers are preferred to long registers. The nicking positions at each strand are determined based on four straightforward to accommodate sequence composition constraints described in the Supplementary Information, Section B.1.

To perform the actual encoding, user files are parsed into n-bit strings which are converted into nicking positions of spatially arranged registers, according to the rule that a ‘1’ corresponds to a nick while a ‘0’ corresponds to the absence of a nick. The number of bits recorded is chosen based on the density of nicks and the length of the register. As an example, the string 0110000100 is converted into the positional code 238, indicating that nicking needs to be performed at the 2^nd^, 3^rd^ and 8^th^ positions (Figure 2a). Note that recording the bit ‘0’ does not require any reactions, as it corresponds to the “no nick” ground state. Therefore, nick-based recording effectively reduces the size of the file to be actually recorded by half. This property of native DNA storage resembles that of compact disks (CD) and other recorders. Note that the length, sequence composition and nicking sites of a register are all known beforehand, so that reading amounts to detecting the positions of the nicks.

**Figure 2.**
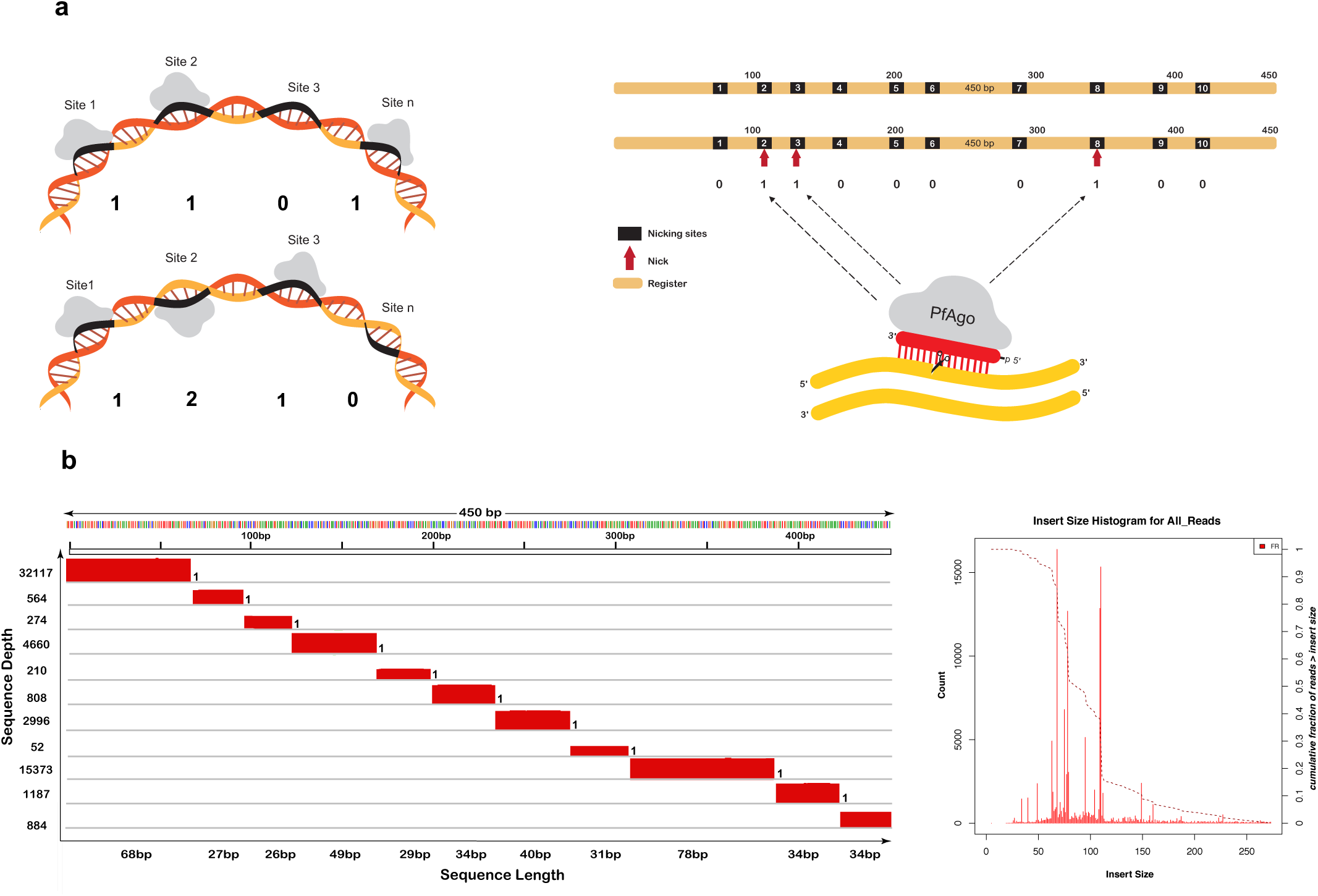
Writing and reading the encoded data. **a**) *Pf*Ago can nick several pre-designated locations on only one strand (left, **top**) or both strands (left, **bottom**), simultaneously. In the first register, the stored content is 110…1, while in the second register, the content is 121…0. The chosen register is a PCR product of a 450 bp *E. coli* genomic DNA fragment with 10 pre-designated non-uniformly spaced nicking positions. The positional code 238 corresponds to the binary vector 0110000100 (right). **b**) The MiSeq sequencing reads were aligned to the reference register to determine the positions of the nicks. The size distribution histogram (right) and coverage plots (left) are then generated based on the frequency and coverage depth of the reads. Coverage plots allow for straightforward detection of nicked and unnicked sites. In the example shown, all the ten positions were nicked, resulting in eleven aligned fragments.

#### Data recording

In the write component of the proposed system (Figure 1, top), binary user information is converted into a positional code that describes where native DNA sequence is to be topologically modified, i.e. nicked. Each nick encodes either log_2_ 2 = 1 bit (if only one strand is allowed to be nicked or left unchanged) or log_2_ 3 = 1.58 bits (if either of the two strands is allowed to be nicked or both left unchanged). Each register is nicked by mixing it with a combination of nicking enzymes that contain guides matched to the collection of sites to be nicked and the information content to be stored. Nicking is performed in parallel on all sites and the recording process takes ∼40 minutes.

To enable fast and efficient data recording, a library of registers with desired nicking site patterns may be created in a combinatorial fashion and in parallel using automated liquid handlers. More precisely, we designed *Pf*Ago guides for all predetermined nicking positions in the chosen register and created registers bearing all 2^10^ = 1024 nicking combinations (Table S3). The recording protocols for the registers are described in the Supplementary Information (Figure S8).

#### Data organization

The registers or mixtures of registers are placed into grids of microplates that enable random access to registers and spatially organize the data, similar to tracks and sectors on disks and tapes. The placement is dictated by the content to be encoded. For our proposed topological storage system prototype, the “plate-based” solution facilitates specific access to data both based on their well position and based on their content, through the use of toehold bitwise random access described in the Additional features section. The proposed positional organization also limits the scope/extent of data destruction during reading, as information is stored in a distributed manner and random-access methods based on PCR may only skew the content in a small number of wells. In its current form, the well-plate approach does not offer desired storage densities but there are several related efforts underway for designing scalable positional data storage systems using MALDI plates (20) or microfluidic devices (21). These solutions are currently not able to handle large data volumes but are expected to improve in the near future.

### The read architecture

The nicked registers are first denatured, resulting in ssDNA fragments of variable lengths dictated by the nicked positions. These length-modulated ssDNA fragments are subsequently converted into a dsDNA library, sequenced on Illumina MiSeq, and the resulting reads are aligned to the known reference register sequence. The positions of the nicks are determined based on read coverage analysis, the insert size distributions and through alignment with the reference sequence; potential nicking sites that are not covered are declared to be ‘0’s (Figure 2a-c).

#### The NGS sequencing process

Since a nick is a very small alteration in the DNA backbone, direct detection of this topological modification on a long DNA strand remains challenging. However, if one denatures the dsDNA and sequences the resulting intra-nick fragments, nick reading becomes straightforward. Hence, in the read component of our system, nicked DNA is first converted to single strands at high temperature; the resulting pool of ssDNA is converted into a library of dsDNA and this library is sequenced using a MiSeq device. The positions of the nicks are determined via read analysis and subsequent reference-based sequence alignment, as the native DNA registers are known a priori.

It is important to note that in our system, high coverages are not essential for accurate data readout, as even relatively small coverages (as low as 1) can be resolved without errors given that the reads can always be aligned to the reference. Also, as the fragment lengths are co-dependent and as the fragments have to jointly “tile/cover” the known reference DNA string, certain missing fragments can be accommodated as well. In simple terms, if one fragment has low or no coverage but leaves a “hole” in the aligned strings with respect to the reference, its presence is noted indirectly upon alignment.

#### NGS readout and alignment experiments

As a proof of concept, we report write-read results for two compressed files, containing a 272-word text file of size 0.4 KB containing Lincoln’s Gettysburg Address (LGA) and a JPEG image of the Lincoln Memorial of size 14 KB (Figure S6). Both files were compressed and converted into ASCII and retrieved with perfect accuracy. Of the whole stored data, a randomly selected set of sample 80 bits, including 5*10-bit registers from the image and 3*10-bit registers from the text file, was sequenced. Given the inherent redundancy of the sequencing process and the careful selection of the nicking sites and register sequences, no error-correction redundancy was needed, and data was perfectly retrieved (Figure 2b, c and Figure S5b-d). As may be observed from Figure 2 and 3, peak coverages are in perfect agreement with the fragment lengths predicted based on the nicking location.

**Figure 3.**
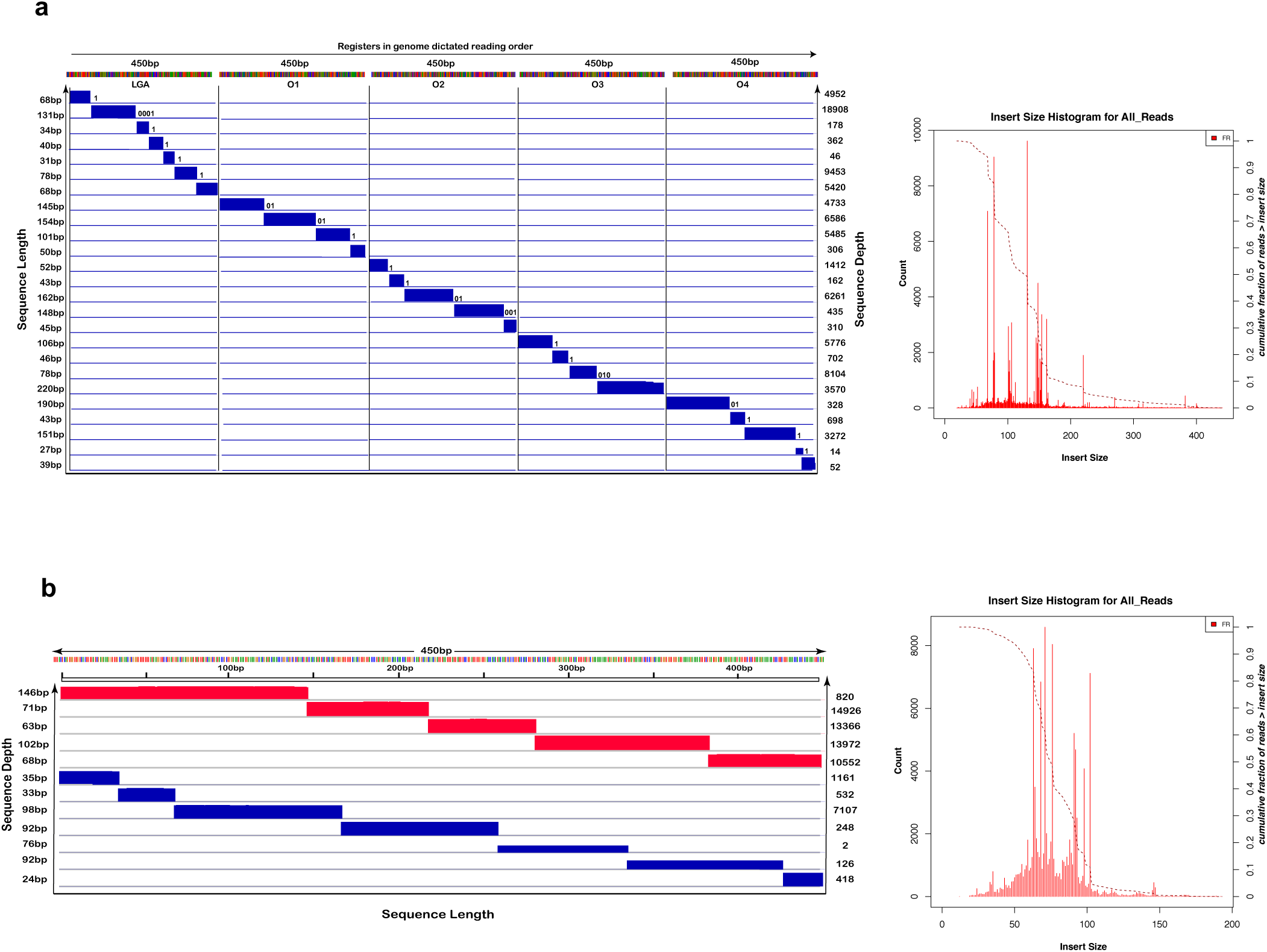
Multi-register and sense-antisense recording. a) Five orthogonal registers used instead of one single register. Each vertical section represents one register in genome dictated reading order, and each row shows the read lengths retrieved after sequencing analysis. Read lengths are recorded on the left and sequencing depths on the right axis. b) Nicking both the sense and antisense strands. Here, the binary format is switched to ternary format, i.e., for each nicking site, a nick on the sense strand denotes 1, a nick on the antisense denotes 2, and no nick denotes 0. The coverage (left) and size distribution (right) plots verify that the register can be nicked on 10 prescribed sites on both strands, six of which are located on the sense strand and four on the antisense strands, corresponding to the 10-bit string of data: 1212121211. The sense-antisense nicking approach may also be used for parallel recording on multiple registers.

#### Alternative readout platforms – solid-state nanopore simulations

A faster, portable and more cost-effective method for reading the nicked DNA registers is via two-dimensional (2D) solid-state nanopore membranes (22). Solid state nanopores also enable reading single information-bearing DNA molecules nondestructively and hence increase the lifecycle of molecular storage systems.

The main drawback of existing solid-state nanopore technologies is the lack of translocation controls that ensure more precise and less noisy nick-position readouts. One approach to mitigate this problem, described in (22), is to extend nicks to toehold, short single-stranded regions on dsDNA created through two closely placed nicks, instead of single nicks. As shown in the Supplementary Information, Section B.8., *Pf*Ago can be used to create toeholds of arbitrary lengths. Experimental evidence (22) reveals that toeholds can be easily and accurately detected using solid-state SiN_x_ and MoS_2_ nanopores.

However, the cost of creating toeholds is twice as high as that of nicks, since one needs two different nicking guides. Hence, one needs to identify alternative mechanisms for direct nick detection. To illustrate the feasibility of the direct readout approach, we performed Molecular Dynamics (MD) simulations along with quantum transport calculations to obtain the ionic and transverse current signals. In order to reduce the computational time of the MD simulations, DNA strands of 30 nucleotides with only one nicked site in the middle were considered. The simulations were performed at 1 V, with a total translocation time of a few nanoseconds. The results indicate a strong inverse correlation between the ionic and electronic sheet current signals along the membrane induced by nicks in MoS_2_ nanopores (Figures S12-S14 & Video S1). We also provide an additional analysis in support of our findings by calculating the Pearson’s correlation between the normalized signals to indicate the time frame when the nick is residing within the pore (Figure S12c).

### Additional Functionalities

In order to increase the per register and per-well storage density of the system we also introduced and experimentally tested three additional nick-based storage methods. The first method involves nicking DNA both at the sense and antisense strands in which case one achieves a 58% increase in density. The second method includes mixing what we refer to as orthogonal registers: Orthogonal registers are DNA fragments isolated from different parts of the genomic DNA that have small sequence similarity scores. The third method involves combinatorial mixing of the same register as dictated by a mathematically-grounded group testing scheme.

#### Sense-antisense recording

As a proof of concept, we used the running example DNA register (LGA) to test recording on both sense and antisense strands. In this setting, the recording alphabet is ternary, i.e., at any predesignated nicking position, a nick on the sense strand denotes 1, a nick on the antisense denotes 2, and no nick denotes 0. This increases the potential storable data for every nicking site ∼1.58 fold. We also experimentally showed that our register can be nicked on 10 prescribed sites on both strands, six of which are located on the sense strand and four on the antisense strands, corresponding to the 10-bit string of data: 1212121211 (Figure 3b).

#### Multi-register recording

To prove the principle of orthogonal register recording, we use the running example LGA register as well as four other orthogonal registers each of 450 bp length. All registers are extracted from the *E. coli* genome (O1-O4), they do not share substrings used as nicking sites and have <50% pairwise similarity matches. Furthermore, the bits stored in the registers are naturally ordered as the registers have an inherent order within their native genomic string. Using orthogonal registers, we designed a total number of 32 nicking positions on all the registers and encoded the title “Gettysburg Address” of size 126 bits via simultaneous nicking of all registers mixed together. This piece of data was also successfully recalled without errors as shown in (Figure 3a). Other technical details regarding implementations with orthogonal registers and with nicks on both DNA strands are provided in the Supplementary Information.

#### Combinatorial Mixing

As each register sequence requires different nicking guides to be used with the nicking endonuclease, the question arises if one can store information on multiple copies of the same register bearing different nicking patterns. Since there is no inherent ordering for the information stored on the same register string within a mixed pool, NGS sequencers cannot resolve the origin of the different fragments through alignment. Multiple solutions for this problem are available. One solution is to store information in the number of nicks, rather than their locations. Another, more effective approach is to use a combinatorial mixing scheme such as group testing or adder channel coding. Group testing effectively allows for mixing of k-out-of-N registers, where *k* is significantly smaller than N and the number of bits scales as *k* log_2_ *N*. The optimal mixing scheme for adder channels is the Lindström scheme (23), which does not impose any restrictions on the value of *k*.

In a nutshell, not all combinations of nicking patterns on the same register sequence can be mixed as one has to prevent multiple distinct alignment solutions for the union of the fragments generated from the mixture. This is accomplished through careful implementation of the Lindström scheme. This coding scheme was originally designed to allow for identifying individual components in a real-valued sum of binary weighted vectors. The scheme is asymptotically optimal in so far that it can discriminate between any of the 2^*N*^ distinct sums of N binary vectors of length as small as 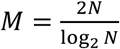. The mathematical explanation behind the scheme is described in detail in Section B.11 of the Supplementary Information, along with a number of experimental test results that indicate that the underlying combinatorial mixtures can be sequenced together and reassembled without any errors.

#### Bitwise Random Access

In addition to allowing for nanopore-based readouts, toeholds ay be used for bitwise random access and toehold-mediated DNA strand displacement, used for engineering dynamic molecular systems and performing molecular computations (24-26). Information is processed through releasing strands in a controlled fashion, with toeholds serving as initiation sites to which these input strands bind to displace a previously bound output strand.

Toeholds are usually generated by binding two regions of synthetic ssDNA and leaving a short fragment unbound. However, we experimentally demonstrated that *Pf*Ago can easily create toeholds in native DNA. To form a toehold, we introduce two nicks at a small distance from each other (in our experiment, 14 bps). Under appropriate buffer and temperature conditions, in a single reaction the strand between the two nicks disassociates, leaving a toehold on the double-stranded DNA (Figure S15).

Fluorescence-based toehold-reporting methods can be used to detect toeholds in a nondestructive manner and hence enable symbol-wise random access. In addition, these methods may be used to estimate the concentration of registers bearing a toehold without modifying the DNA registers. We illustrate this process on a register encoding 0010000000, with a toehold of length 14 nts at the nicking position 3. As shown in Figure 4a, a fluorophore and quencher labelled reporter strand with a sequence complementary to the toehold can hybridize to the toehold segment, producing a fluorescence signal resulting from an increase of the distance between the fluorophore and the quencher. We were also able to reliably measure different ratios of native DNA fragments with and without toeholds within 20 mins (Figure 4b). Since the reporter has a short single stranded overhang, it can be pulled off from the register upon hybridization, making the readout process non-destructive (Polyacrylamide gel electrophoresis analysis, Figure 4c). This feature equips our proposed storage system with unique nondestructive bitwise random access, since one is able to design specific reporters to detect any desired toehold sequence which accompanies a nick. It also enables in-memory computations on data encoded in nicks (15,16).

**Figure 4.**
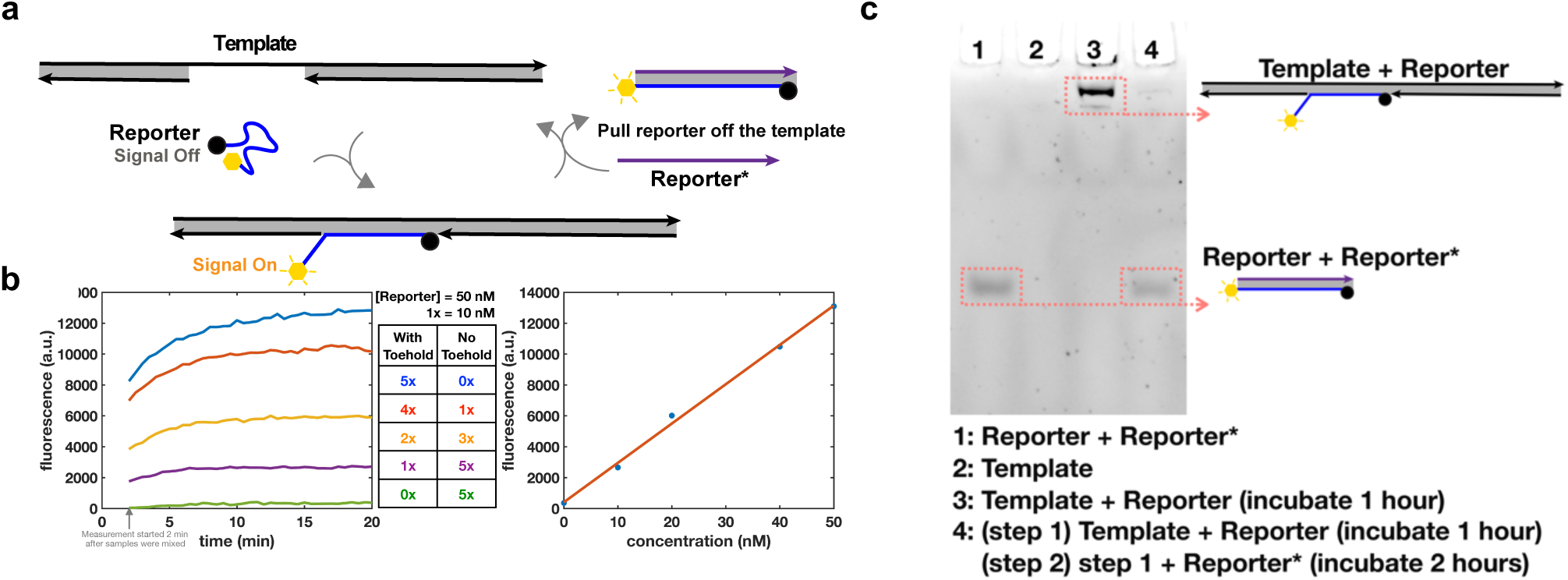
Non-destructive bitwise random access. **a)** Non-destructive detection of toeholds through a fluorophore and quencher labelled Reporter strand. Once the Reporter hybridizes with the toehold on the register strand, a fluorescence signal is observed due to the increase of the distance between the fluorophore and quencher. The Reporter strand can be pulled off from the register once the Reporter* strand hybridizes with the Reporter. **b)** Kinetics of detecting the concentrations of registers with and without toeholds in a mixtures (left). The fluorescence signals saturate within 20 minutes. The samples were mixed no more than 2 min before measurement. The concentration of toehold-ed DNA can be accurately quantified through fluorescence intensity (right), as it increases linearly with the concentration of the registers with toehold. **c)** PAGE gel results for non-destructive detection of a toehold. The gel was not stained with other fluorescence dyes, thus only the species with self-fluorescence is observed. After adding the Reporter, a large size complex appears in lane 3, indicating hybridization of the Reporter and the register. After the Reporter* is added, as seen in lane 4, the large size complex in lane 3 no longer exhibits self-fluorescence, indicating that the Reporter strand is pulled off from the register.

Note that since the positions of the nicks are predetermined and since orthogonal registers are mixed, one only needs to have one reporter sequence available for each bit location. For LGA register, one need only ten different fluorophores for bitwise random access to all ten positions simultaneously. If access is sequential, only one fluorophore is needed. This is to be contrasted with the first random access scheme we proposed and reported on in (4). There, one can only access individual oligos through a time-consuming process of PCR and sequencing.

## Discussion

By reprogramming *Pf*Ago to serve as a universal nickase and using native *E. coli* DNA strings we implemented the first DNA-based storage system that mitigates the use of long synthetic DNA strands for storing user information, and records data in DNA backbone rather than the sequence content. In contrast to synthesis-based methods, our platform has significantly reduced writing latency due to the ability of the enzyme to encode information in parallel. Also, our approach has exceptionally high reliability, in contrast to synthetic DNA-based data storage that suffer from high rates of synthesis and missing oligo errors. Note that the inherently high error-rate of synthetic platforms is further amplified when storing compressed data since errors in compressed formats tend to exhibit catastrophic error propagation. Hence, nick-based storage schemes are exceptionally well-suited for storing compressed data representation. Furthermore, as nicks may be easily superimposed on synthetic DNA molecules they may also be used to encode metadata that can be erased (through simple ligation) and recreated or altered in a time efficient manner.

Our storage system design also enables enzyme driven toehold creation for bitwise random access, a unique feature not paralleled by any other implementation. Furthermore, nick-based storage for the first time allows for in-memory computing on the data stored on molecular media. As reported in a companion paper (15), DNA-based strand displacement data processing schemes capable of parallel, in-memory computation, can perform on such system and thereby eliminate the need for sequencing and synthesizing new DNA on each data update.

With further improvements and optimization, our system may also allow for cost-efficient scaling since **a**. long registers and mixtures of orthogonal registers may be nicked simultaneously; **b**. most compressed data files do not contain all possible 10-mers or compositions of orthogonal *k*-mers so that not all guide combinations are needed. **c**. genomic DNA and *Pf*Ago, as the writing tool, are easily extracted and used in standard laboratory settings, and the mass of the created DNA products by far exceeds that of synthetic DNA. This may significantly increase the number of readout cycles with NGS devices.

**Table 1.** Comparison of synthetic and native DNA-based data storage platforms. Native DNA-based platforms outperform synthetic DNA-based approaches in all performance categories, except for storage density.

| DNA-based Storage Method | Writing Latency | Reading Latency | Enables Computation? | Bit-wise Random Access | Maximum achievable physical Density | Information Density | (Optimal)Coding Loss (10) |
| --- | --- | --- | --- | --- | --- | --- | --- |
| Synthesis - based | Sequential de novo synthesis/ Hours | NGS/hours | × | × | 200 Ebytes/g (9) | < 2 bits/bp (to account for coding loss, usually ~1.5 bits/bp) | 21% (10,11) |
| This work | Parallel Nicking/ < 40 min | NGS followed by reference alignment/ hours | ✓ | ✓ | 4 Ebytes/g | 0.036 bits/bp | 0% |

## Materials and Methods

### Guide DNA selection and positional coding

To minimize off-site nicking rates, increase the efficiency and accuracy of gDNA-binding, and eliminate readout errors, the nicking regions were selected by imposing the following constraints: each region is of length 16 bps, which allows each gDNA to bind to a unique position in the chosen registers; each region has a GC content in the range 20-60%; there are no G repeats longer than three (this feature is only needed in conjunction with nanopore sequencing); the gDNAs are at Hamming distance at least eight from each other; and the nicking sites are placed at least 25 bps apart. Positional coding is performed based on the number of orthogonal registers used, and the number of nicking positions selected on each register. For the single-register implementation with one-sided nicking, ten positions were selected on a 450 bp genomic fragment, bearing 10 bits.

Although choosing longer registers with a larger number of nicking positions is possible and indeed easily doable, we selected the given length for our proofs of concept in order to accommodate different sequencing technologies. The five orthogonal register implementation may encode 32 bits with one sided nicking, and roughly 50 bits with two-sided nicking. Hence, each binary encoded message is parsed into blocks of length either 10, or 32 or 50 bits, which are recorded via nicking.

### Genomic DNA isolation and PCR amplification

Genomic DNA was extracted from an overnight culture of *E. coli* K12 MG1655, using the Wizard^®^ Genomic DNA Purification Kit (Promega). The kit can be used for at least 100 isolations. One extraction yields up to 100 µg of genomic DNA (from 5 ml overnight culture) which can be used for several hundreds of amplification reactions. Isolated genomic DNA was subsequently stored at 4 **°**C. Two means of reducing the cost of DNA extraction are to either manually extract the DNA or fully automate the process. The former approach can lead to a very high yield of DNA at a small cost, and all used buffers and reagents can be made in-house. For more information, see [https://bio-protocol.org/bio101/e97#biaoti1286].

DNA amplification was performed via PCR using the Q5 DNA polymerase and 5X Q5 buffer (New England Biolabs) in 50 µl. All primers purchased from Integrated DNA Technologies (IDT). In all PCR reactions, 10-50 ng of *E. coli* genomic DNA and 25 pmol of forward and reverse primers were used. The PCR protocol consists of: 1) 3 min at 98 **°**C, 2) 20 s at 98 **°**C, 3) 20 s at 62 **°**C, 4) 15 s at 72 **°**C, 5) go to step 2 and repeat the cycle 36 times, 6) 8 min at 72 **°**C. Each PCR reaction produced ∼2-2.5 µg of the register string, sufficient for >100 reactions. PCR products were run on 1% agarose gel and purified using the Zymoclean gel DNA recovery kit (Zymo Research).

### Enzyme expression and purification

Enzyme expression and purification was performed as previously described (13). More than 200 nmols of *Pf*Ago were purified from 1 L of *E. coli* culture, enabling >50,000 reactions.

### *Pf*Ago nicking experiments

For ease of access and spatial organization of data, pools of registers are kept in 384-well plates. The distribution of the registers and enzymatic reagents was performed manually (when transferring volumes of reagents with volumes of the order of microliters) or using the Echo^®^ 550 liquid handler (LABCYTE) (when transferring minute volumes of reagents of the order of nanoliters). The latter allows for testing the reaction efficiency of *nanoliters* of reagents and it also represents a faster and more efficient manner of liquid handling at larger scales. *Pf*Ago reactions were performed in buffer conditions including 2 mM MnCl_2_ 150 mM NaCl, and 20 mM HEPES, pH 7.5, and a total volume of 10-50 μL. After adding the buffer, dsDNA registers, ssDNA phosphorylated gDNAs and the enzyme, the sample was thoroughly mixed by pipetting 6-8 times. Nicking was performed based on the following protocol: 1) 15 min at 70 **°**C, 2) 10 min at 95 **°**C, 3) gradual decrease of temperature (0.1 **°**C/s) to 4 **°**C. In all reactions, we used 3.75-5 pmol of *Pf*Ago and 20-50 ng of the register. gDNAs were either phosphorylated using T4 Polynucleotide Kinase (NEB) in lab or phosphorylated guides were purchased from IDT. For each nicking reaction, a (2-10):1 ratio of guides to enzymes was formed. All guides were used in equimolar mixtures.

### Cas9 nickase experiments

The Cas9 D10A nickase was purchased from IDT (Alt-R^®^ S.p. Cas9 D10A Nickase); crRNAs were designed via IDT’s Custom Alt-R^®^ CRISPR-Cas9 guide RNA design tool. Both crRNAs and tracrRNAs were purchased from IDT and hybridized based on the manufacturer’s protocol. The 10x Cas9 reaction buffer included: 200 mM HEPES, 1M NaCl, 50 mM MgCl_2_, 1mM EDTA, pH 6.5. All Cas9n nicking reactions were set-up based on the manufacturer’s protocol and performed at 37 °C for 60 min.

### Protocol verification via gel electrophoresis

ssDNA gel analysis was performed using a 2% agarose gel. Nicked dsDNA samples were first denatured at high temperature (99 °C) for 10 min, and immediately cooled to 4 °C. The ssDNA products were then run on a pre-made 2% agarose Ex-Gel (Thermo Fisher Scientific).

### Sample preparation for MiSeq sequencing

All nicked PCR products (obtained either via *Pf*Ago or Cas9n reactions) were purified using the Qiaquick PCR purification kit (QIAGEN) and eluted in ddH_2_O. The dsDNA registers were denatured at 99 **°**C for 10 min, and immediately cooled down to 4 **°**C. The ssDNA samples were first quantified via the Qubit 3.0 fluorometer. Next, the Accel-NGS^®^ 1S plus DNA library kit (Swift Biosciences) was used for library preparation following the manufacturer’s recommended protocol. Prepared libraries were quantitated with Qubit, and then run on a DNA Fragment Analyzer (Agilent, CA) to determine fragment sizes, pooled in equimolar concentration. The pool was further quantitated by qPCR. All steps were performed for each sample separately and no nicked DNA samples were mixed.

### MiSeq sequencing

The pooled libraries were loaded on a MiSeq device and sequenced for 250 cycles from each end of the library fragments with a Nano V2 500 cycles kit (Illumina). The raw fastq files were generated and demultiplexed with the bcl2fastq v2.20 Conversion Software (Illumina).

### Reference alignment

Data was processed using a Nextflow-based workflow (27), implemented as follows. Sequence data was trimmed using Trimmomatic v.0.36 (28) in paired-end mode using the options “ILLUMINACLIP: adapters/TruSeq3-PE-2.fa:2:15:10 LEADING:20 TRAILING:20 SLIDINGWINDOW:4:15 MINLEN:20”. Reads were aligned to the reference sequence using bwa v 0.7.10 (29) with the command “bwa mem -t 12 <REFERENCE> <R1> <R2>“. Alignments were sorted and processed using samtools v1.6 (30). Insert size statistics were collected using Picard v.2.10.1 (31). Aligned files (BAMs) were then split based on expected fragment size using sambamba (32) with the option “sambamba view -t 4 -f bam -h -F “(template_length >= [LOWER] and template_length <= [UPPER]) or (template_length >= -[UPPER] and template_length <= – [LOWER])”, with the upper and lower bound settings in brackets originally set to allow for one additional base greater and lesser than the expected size. Read coverage files were then generated using bedtools (33) and bedGraphToBigWig (34). Alignment and coverage information was visualized in IGV v2.3.10 (35). All the scripts used for data analysis are available from the corresponding authors upon request.

### Nanopore simulations

To obtain the trajectories of the nicked molecule translocating through the nanopore, all-atom Molecular Dynamics simulations were performed using NAMD (36). For these simulations, the DNA structure (30 nucleotides around the 5^th^ nicking site in the register) was obtained from the 3D-DART webserver (37) and described using the CHARMM27 force field (38). Appropriate backbone molecules were manually removed to create the nicks in the desired locations of the strand. Note that in order to obtain a stronger nanopore current signal from the DNA backbone, the PO_3_ groups located at the nicked position may be removed by treatment of the nicked dsDNA with a phosphatase enzyme such as BAP (bone alkaline phosphatase).

The DNA molecule was placed just above the nanopore of a Molybdenum disulfide (MoS_2_) membrane to ensure a successful translocation process. The nanopore membrane and the biomolecule were then solvated in a water box with ions (K^+^ and Cl^-^) placed randomly to reach a neutrally charged system of concentration 1 M. Van der Waals energies were calculated using a 12 Å cutoff. Each system was minimized for 5000 steps and further equilibrated for 2 ps in an NPT ensemble, where the system was maintained at 1 atm pressure by a Langevin Piston (39) and at constant 300 K temperature using a Langevin thermostat. After equilibration, an external electric field was applied to the system in vertical direction to drive the nicked DNA through the nanopores.

A trajectory file of molecules driven through the nanopore by the applied electric field obtained from the MD simulations was used to calculate the ionic current via Equation (1) (40), where *q*_*i*_ and *z*_*i*_ denote the charge and z-coordinate of ion *i*, respectively; *V* denotes the voltage bias (1 V) and *L* the length of the water box along the z-direction, while *N* represents the number of ions and *Δt* the interval between the trajectory frames:

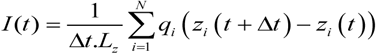

For each frame of the trajectory, the electrostatic potential is calculated using the following non-linear Poisson Boltzmann formula

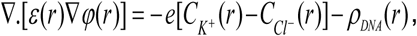

where *ρ*_*DNA*_ denotes the charge density of DNA, *ε*(*r*) the local permittivity, and where 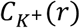 and 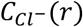 equal the local electrolyte concentrations of K^+^ and Cl^-^ and obey the Poisson-Boltzmann statistics. The detailed description of the method used is outlined elsewhere (41). The calculated electrostatic potential is used to obtain the transverse sheet conductance in MoS_2_ quantum point contact nanopore membranes. The electronic transport is formulated as a self-consistent model based on the semi-classical thermionic Poisson-Boltzmann technique using a two-valley model within the effective mass approximation. The calculated conductance at a given energy mode is described according to

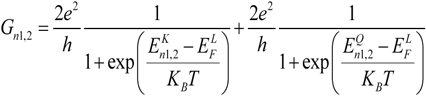

where 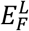 denotes the quasi-Fermi level and is set depending on the carrier concentration (chosen to be 10^12^ cm^-2^); in addition, *n*_*1,2*_ represents the energy modes of the two conductance channels while 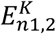 and 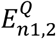 stand for the energy modes at these two channels caused by the effective masses K and Q, respectively. A detailed discussion of the thermionic current model is described elsewhere (42).

The MoS_2_ membrane was modeled using VMD (43) from a basic unit cell. The Lennard–Jones parameters for MoS_2_ are σ_Mo-Mo_ = 4.4 Å, ε_Mo-Mo_ = 0.0135 kcal/mol and σ_S-S_ = 3.13 Å, ε_S-S_ = 0.3196 kcal/mol, taken from Stewart et Al (44). All MoS_2_ atoms were fixed to their initial positions. Pore of diameter 2.6 nm is drilled by removing the atoms whose coordinates satisfy the condition x^2^ + y^2^ ≤ r^2^, where r is the radius of the pore. Nick in each dsDNA strand were created by manually removing the phosphodiester bond and phosphate atoms in the backbone (45).

As a final remark, we observe that the simulations in Figure S12-S14 indicating strong negative correlations of the global minimum and maximum of the sheet and ion current may be interpreted as follow: When nicked DNA translocates through the pore, the oscillations of the nicked backbone allow more ions to pass through the pore, leading to a steep increase (maximum) in the ion current. At the same time, the absence of the PO_3_ group charges leads to a decrease in the sheet current to its global minimum.

## Acknowledgements

This work was funded by the DARPA W911NF-18-2-0032 Molecular Informatics program. We thank SMH Tabatabaei Yazdi, and Jianhao Peng for many discussions and help. The authors also gratefully acknowledge supercomputing resources offered by Extreme Science and Engineering Discovery Environment (XSEDE) grant: TG-MCB170052 and Blue Waters grant: baxi.

## Author contributions

O.M. and H.Z. developed the nicking-based data recording platform. SK.T. performed the nicking and toehold creation experiments and all other writing experiments. D.S., B.W. and O.M. designed the bitwise random-access system. D.S. and B.W. performed the bitwise access experiments. JP.L. and N.A. implemented the nanopore simulations. A.G.H., D.S., O.M. and SK.T. designed the readout system, while A.G.H. performed all the MiSeq sequencing experiments. C.F. performed the MiSeq read analysis and reference alignment, while B.E. performed initial *Pf*Ago nicking activity verifications and helped with nicking experiment designs and protein purifications.

## Correspondence and requests for materials

should be addressed to O.M. or H.Z.

## References and Notes

1. Skinner. G.M, Visscher. K, Mansuripur. M, Biocompatible writing of data into DNA, Journal of Bionanoscience, 1, 1–5 (2007).

2. Church. G. M., Gao. Y, Kosuri. S. Next-generation digital information storage in DNA. Science 337, 1628–1628 (2012).

3. Goldman. N, Bertone. P, Chen. S, Dessimoz. C, Leproust. E, Sipos. B, Birney. E. Towards practical, high-capacity, low-maintenance information storage in synthesized DNA. Nature 494, 77–80 (2013).

4. Yazdi. S. H. T., Yuan. Y, Ma. J, Zhao. H, Milenkovic. O. A rewritable, random-access DNA-based storage system. Sci. Rep. 5, 14138 (2015).

5. Grass. R. N., Heckel. R., Puddu. M., Paunescu. D. & Stark. W. J. Robust chemical preservation of digital information on DNA in silica with error-correcting codes. Angew. Chem. Int. Ed. 54, 2552–2555 (2015).

6. Yazdi. S. H. T., Gabrys. R., and Milenkovic. O. “Portable and error-free DNA-based data storage.” Scientific reports 7.1 (2017).

7. Shipman. S. L., Nivala. J., Macklis. J. D. & Church. G. M. CRISPR–Cas encoding of a digital movie into the genomes of a population of living bacteria. Nature 547, 345–349 (2017).

8. Zhirnov. V., Zadegan. R. M., Sandhu. G. S., Church. G. M. & Hughes. W. L. Nucleic acid memory. Nat. Mater. 15, 366–370 (2016).

9. Erlich. Y. & Zielinski. D. DNA fountain enables a robust and efficient storage architecture, Science, 355, 950–954 (2017).

10. Yazdi. S. H. T., Kiah. HM, Ruiz-Garcia. E, Ma. J, Zhao. H., Milenkovic. O. DNA-based storage: Trends and Methods. IEEE Transactions on Molecular, Biological and Multi-Scale Communications, 1, 3, 230–248, (2015).

11. Laure. C., Karamessini. D., Milenkovic, O., Charles. L., Lutz. J.F., Coding in 2D: using intentional dispersity to enhance the information capacity of sequence-coded polymer barcodes. Angewandte Chemie International Edition, 55(36), pp.10722–10725 (2016).

12. Milenkovic. O., Gabrys. R., Kiah. H. M, Yazdi. S. H. T., Exabytes in a Test Tube. IEEE Spectrum, 55 (5), 40–45 (2018).

13. Palluk. S., Arlow. D. H, de Rond. T, Barthel. S, Kang. J. S, Bector. R, Baghdassarian. H. M, Truong. A. N, Kim. P.W, Singh. A. K, Hillson. N. J, Keasling. J. D., De novo DNA synthesis using polymerase-nucleotide conjugates, Nat Biotechnol., 36, 645–650 (2018).

14. Pan. C., Yazdi. S. H. T., Tabatabaei. SK., Hernandez. A. G., Schroeder. C., Milenkovic. O., Image processing in DNA, 1910.10095 (2019).

15. Wang. B., Chalk. C, Soloveichik. D, SIMDNA: Single Instruction, Multiple Data Computation with DNA Strand Displacement Cascades DNA 25 Conference, Seattle, WA, U.S.A. (2019).

16. Chen. T, Riedel. M, Parallel Binary Sorting and Shifting with DNA, 11th International Workshop on Bio-Design Automation (IWBDA), Cambridge, England, U.K. (2019).

17. Chen. K., Kong. J., Zhu. J., Ermann. N., Predki. P., Keyser. UF., Digital data storage using DNA nanotructures and solid-state nanopores, Nano Letters, 19 (2), 1210–1215 (2019).

18. Andres. C, Jinek. M. In vitro enzymology of Cas9, Methods Enzymol. 546, 1–20 (2016).

19. Enghiad. B., Zhao. H. Programmable DNA-guided artificial restriction enzymes. ACS Synth. Biol., 6, 752–757 (2017).

20. Kennedy. E., Arcadia. C. E., Geiser. J., Weber. P. M., Rose. C., Rubenstein. B. M., Rosenstein. J. K. Encoding information in synthetic metabolomes, PLoS ONE, 14 (7); e02173064 (2019).

21. Newman. S., Stephenson. A.P., Willsey. M. et al. High density DNA data storage library via dehydration with digital microfluidic retrieval. Nat Commun 10, 1706 (2019).

22. Liu. K., Pan. C, Kuhn. A, Nievergelt. A.P, Fantner. G, Milenkovic. O, Radenovic. A, Detecting topological variations of DNA at single-molecule level, Nature Communications, 10, 3 (2019).

23. B. Lindström et al., J. N. Srivastava, ed., A survey of Statistical Design and Linear Models, North-Holland Publishing Company, (1975).

24. Yurke. B, Turberfield. A.J, Mills. A.P, Simmel. F.C, Neumann. J.L, A DNA-fueled molecular machine made of DNA. Nature, 406:605–608 (2000).

25. Zhang. DY., Seelig. G, Dynamic DNA nanotechnology using strand-displacement reactions. Nat Chem., 3:103–113 (2011).

26. Wang. B., Thachuk. C, Ellington. A., Winfree. E., Soloveichik. D., Effective design principles for leakless strand displacement systems, PNAS, 115 (52), E12182–E12191 (2018).

27. Di Tommaso, P., et al., Nextflow enables reproducible computational workflows. Nat Biotechnol, 35(4) 316–319 (2017).

28. Bolger, A.M., M. Lohse, and B. Usadel, Trimmomatic: a flexible trimmer for Illumina sequence data. Bioinformatics, 30(15): p. 2114–20 (2014).

29. Li, H. Aligning sequence reads, clone sequences and assembly contigs with BWA-MEM. ArXiv e-prints, 1303 (2013).

30. Li, H., et al., The Sequence Alignment/Map format and SAMtools. Bioinformatics, 25(16): p. 2078–9 (2009).

31. Institute, B. Picard Tools. [2018 2017]; Available from: http://broadinstitute.github.io/picard/.

32. Tarasov, A., et al., Sambamba: fast processing of NGS alignment formats. Bioinformatics, 31(12): p. 2032–4 (2015).

33. Quinlan, A.R., BEDTools: The Swiss-Army Tool for Genome Feature Analysis. Curr Protoc Bioinformatics, 47: p. 11 12 1–34, (2014)

34. Kent, W.J., et al., BigWig and BigBed: enabling browsing of large distributed datasets. Bioinformatics, 26(17): p. 2204–7 (2010).

35. Thorvaldsdottir, H., J.T. Robinson, and J.P. Mesirov, Integrative Genomics Viewer (IGV): high-performance genomics data visualization and exploration. Brief Bioinform, 14(2): p. 178–92 (2013).

36. Phillips, J. C. et al. Scalable molecular dynamics with NAMD. J. Comput. Chem. 26, 1781–1802 (2005).

37. Van Dijk, M. & Bonvin, A. M. J. J. 3D-DART: a DNA structure modelling server. Nucleic Acids Res. 37, W235–W239 (2009).

38. Foloppe, N. & MacKerell, Jr., A. D. All-atom empirical force field for nucleic acids: I. Parameter optimization based on small molecule and condensed phase macromolecular target data. J. Comput. Chem. 21, 86–104 (2000).

39. Feller, S. E., Zhang, Y., Pastor, R. W. & Brooks, B. R. Constant pressure molecular dynamics simulation: The Langevin piston method. J. Chem. Phys. 103, 4613–4621 (1995).

40. Aksimentiev, A., Heng, J. B., Timp, G. & Schulten, K. Microscopic Kinetics of DNA Translocation through Synthetic Nanopores. Biophys. J. 87, 2086–2097 (2004).

41. Girdhar, A., Sathe, C., Schulten, K. & Leburton, J.-P. Graphene quantum point contact transistor for DNA sensing. Proc. Natl. Acad. Sci. 110, 16748–16753 (2013).

42. Sarathy, A. & Leburton, J. P. Electronic conductance model in constricted MoS_2_ with nanopores. Appl. Phys. Lett. 108 (2016).

43. Humphrey, W.; Dalke, A.; Schulten, K. VMD – Visual Molecular Dynamics. J. Mol. Graphics, 14, 33–38 (1996).

44. Stewart, James A., and D. E. Spearot. “Atomistic simulations of nanoindentation on the basal plane of crystalline molybdenum disulfide (MoS2).” Modelling and Simulation in Materials Science and Engineering 21, no. 4: 045003 (2013).

45. Aymami, J. et al. Molecular structure of nicked DNA: a substrate for DNA repair enzymes. Proc Natl Acad Sci USA, 87, 2526 (1990).

