## Supplementary Information for "DNA Punch Cards: Storing Data on Native DNA Sequences via Nicking" (from another version of this preprint: version 5, 2019-08-04)

**This file contains:**

Supplementary Notes and Data
Supplementary Figures S1-S17
Tables S1-S5
References

**Other Supplementary Materials for this manuscript include the following:**

Video S1

### Table of Contents

**A**. **List of the Used Abbreviations (3)**

**B**. **Supplementary Notes and Data (4-54)**
1. Design of guide DNAs for *Pf*Ago (4-5)
2. Nicking activity verification (6-8)
3. Comparison of the nicking activity of *Pf*Ago and Cas9 nickase (9-15)
4. Lincoln's Gettysburg Address (LGA) and Lincoln's Memorial Image (LMI), writing and readout (16-21)
5. Data organization (22-23)
6. Approaches to increase the information density (24-33)
7. Simulations of solid state nanopore nick detection (34-38)
8. Toehold creation (39-40)
9. Non-destructive toehold detection (41-43)
10. Extended discussion on Table 1 (44-47)
11. Combinatorial register mixing (48-54)

**C. Additional Supplementary Tables (55-59)**
Table S4; The used sequences (55-56)
Table S5; The used oligos (57-59)

**D**. **References (60)**

#### A. List of the Used Abbreviations

**NEN:** Nicking Endonuclease
**LGA:** Lincoln's Gettysburg Address
**LMI:** Lincoln Memorial Image
**O1-4:** Orthogonal Sequences 1-4

#### B. Supplementary notes and data

##### 1. Design of guide DNAs (gDNAs) for *Pf*Ago

The guide DNAs for *Pf*Ago are designed using the following criteria:

1) All are 16 nt long, 5'-phosphorylated ssDNAs,
2) All have a GC content of more than 20% and less than 80%,
3) Long runs of G/Cs or A/Ts (>7-mers) are avoided,
4) All nicking sites (between positions 10 & 11 of the guide) are kept at least 25 nt apart, in order to prevent the inter-nicked strands from disassociation in room temperature.

The first two constraints do not impose strong limitations on the number of potential nicking sites. Still, there exists a fundamental limit of how many sites may be nicked in the *E. coli* genome. The reason for this limit is that the guides are of length 16 nts, and there are $4^{16}$=4,294,967,296 possibilities for the guides. In comparison, the length of the genome is 4,639,694 bps, resulting in 4,639,678 potential nicking sites. Clearly, due to fewer options for the guides, not all these positions may be nicked. In addition, sequences of length 16 within the genome may not be unique. Genome wide search (Table S1) reveals that length 2816 is the smallest length for which each substring of that length appears at most once in the genomic sequence of the chosen *E. coli* strain. Hence, multiple nicking locations may have to be eliminated from consideration to avoid undesired false bit recording. Furthermore, this finding suggests using shorter registers as these tend to have smaller unique length. Table S1 also reveals that starting from length 7, some k-mers do not appear in the genome. Another constraint arises from the fact that the best distance-4 codes of length 16 have size 2048 (1). This suggests that in each round of nicking, only 2048 position should be targeted simultaneously. This is not a major issue since genomic DNA is first fragmented so that nicking may be performed separately on subsets of fragments. The GC-constraint introduces negligible coding loss. The run length <7 constraint simultaneously imposed on all bases has capacity zero, which follows from the fact that the largest root of the capacity-defining equation in a complex variable z, which reads as $z^8(z-4)+3=0$, is 1 (2). But since our run length has to be imposed only on the symbol G and only when nanopore sequencers are used, the reductions in the sequence space is limited to 10-15%.

**Table S1| Count of distinct k-mers in the *E. coli* genome, for k=1,…,13.** Results are shown for $\mathbf{k} \leq \mathbf{13}$. As 13 is the upper bound on the length for which the number of distinct k-mers increases. For k>13, the number of distinct k-mers will decrease. The k-mer counting code is available upon request.

| **K (length)** | **Number of distinct K-mers** | **Log Base 4 of the number of distinct K-mers** |
|---|---|---|
| 1 | 4 | 1 |
| 2 | 16 | 2 |
| 3 | 64 | 3 |
| 4 | 256 | 4 |
| 5 | 1024 | 5 |
| 6 | 4096 | 6 |
| 7 | 16,383 | 6.9999 |
| 8 | 65,360 | 7.9981 |
| 9 | 256,527 | 8.9844 |
| 10 | 898,115 | 9.883 |
| 11 | 2,196,861 | 10.5335 |
| 12 | 3,478,960 | 10.8651 |
| 13 | 4,170,362 | 10.9959 |

All gDNAs were either ordered with 5'-phosphorylation or phosphorylated via T4 polynucleotide kinase (New England Biolabs) based on the manufacturer's protocol. All sequences and maps shown in the main text and this file are visualized via SnapGene Viewer 4.2.9 (SnapGene®). All reference-based alignments and coverage plots are visualized in IGV v2.3.10 (3).

#### 2. Nicking activity verification

Previously, *Pf*Ago was shown as a programmable Artificial Restriction Enzyme (ARE), able to perform double stranded digestion using two 5'-phosphorylated guide DNAs (gDNA), each binding to one strand of a target dsDNA, and cutting both strands at the target sequence between positions 10 and 11 of the gDNA. However, its DNA nicking activity was not demonstrated. To this end, we used two commercially available nicking endonucleases (NEN), Nb.BtsI and Nt.BstNBI (New England Biolabs), to nick one strand of a randomly amplified linear dsDNA from the *E. coli* K12 MG1655 genome. We used *Pf*Ago with one gDNA targeting the complementary sequence of the NEN's target region (Figure S1). These two reactions need to be done sequentially, since none of the enzymes is active in the other's buffer conditions. We performed the nicking reactions in both orders (*Pf*Ago → NEN or NEN → *Pf*Ago) and observed the same results. The final digestion product was run on 1% agarose gel to determine the nicking efficiency of *Pf*Ago.

Based on the buffer conditions used for *Pf*Ago-based restriction digestion activity previously reported[1], we hypothesized that increasing the $Mn^{2+}$ concentration in the reaction buffer may help the nicking efficiency since: 1) $Mn^{2+}$ is shown to be essential for the enzyme to get activated[1] and, 2) Our previous experiments had indicated (Data not shown) that with two positive charges, $Mn^{2+}$ may help to keep the strands separated at high temperatures in which the enzyme works. Other buffer components (NaCl and HEPES) were kept constant at their concentrations as noted elsewhere[1]. As shown in Figure S1, *Pf*Ago shows its highest nicking activity at [$MnCl_2$] = 2mM ([$MnCl_2$] =4 mM in a 2X buffer). Therefore, we determined our 2X buffer components concentrations as: [NaCl] = 300 mM, [HEPES]= 40 mM, [$MnCl_2$] = 4 mM, pH = 7.5. The best reaction temperature was also determined to be 95 °C (Figure S1).

Further verification was performed on ssDNA using 2% agarose gel. Figure S2a shows the process of sample preparation for gel electrophoresis analysis. After the nicking reaction via *Pf*Ago, dsDNA samples are denatured at high temperature (99 °C, for 5-10 min), followed by immediately cooling down to 4 °C. The ssDNA products are then run on 2% agarose gel. In this step, we performed all experiments on a selected 450 bp PCR-amplified linear DNA from *E. coli* K12 MG1655 genome. To determine the best reaction temperature and prove that the previously mentioned buffer conditions (buffer KT: 4 mM $MnCl_2$, 300 mM NaCl and 40 mM HEPES, pH 7.5) offer excellent reaction conditions for nicking, we also screened the results through different temperatures and $MnCl_2$ concentrations (Figure S2b, c).

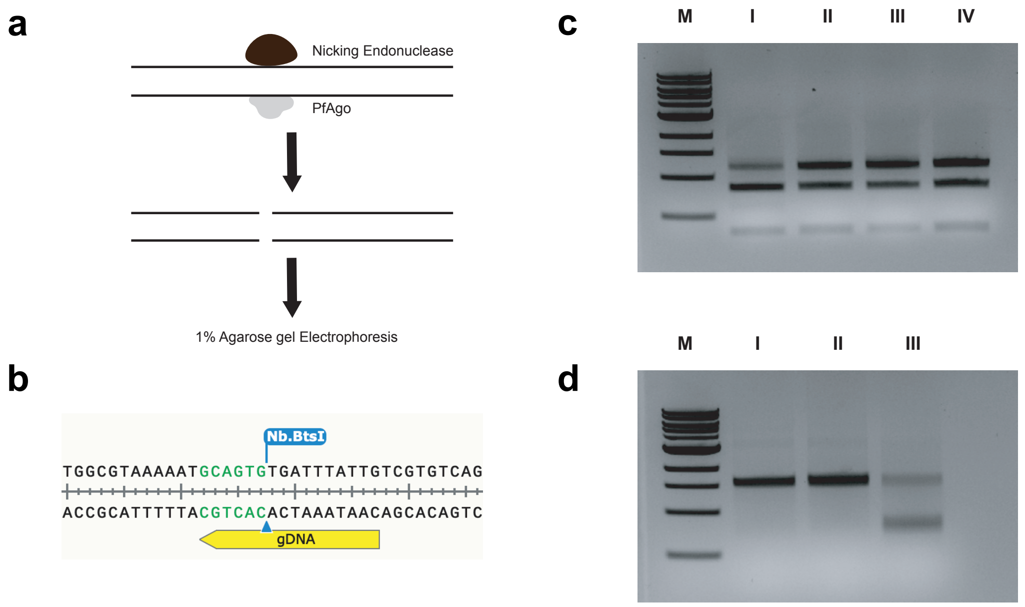


**Figure S1 | Verification of *Pf*Ago's nicking activity using nicking endonucleases. a)** A commercial nicking endonuclease and *Pf*Ago are used to double digest a randomly selected piece of dsDNA from the *E. coli* K12 MG1655 genome. The final digestion product was run on 1% agarose gel to visualize the result. In case of full digestion, two pieces of DNA were expected and observed. **b)** A flanking region of the target position in a chosen 1 kb PCR product of *E. coli* genome. The 16 nt 5'-phosphorylated gDNA (Yellow) targets the upper strand and cuts the phosphodiester bond between nucleotides at positions 10 and 11 (T and G, respectively). The nicking endonuclease cuts a phosphodiester bond, in this example, between A and C in the strand appearing at the bottom. **c)** A randomly chosen 1.5 kb PCR product of the *E. coli* K12 MG1655 genome is nicked at the same site on both strands with Nt.BstNBI, and *Pf*Ago in different temperatures (Lane I: 95 °C, Lane II: 90 °C, Lane III: 87 °C, Lane IV: 85 °C). **d)** *Pf*Ago shows perfect nicking activity in 2 mM $MnCl_2$ (Lane III), while there is almost no activity observed in 0.5 and 1 mM $MnCl_2$ (Lanes I and II). All *Pf*Ago reactions were performed at 95 °C. In this experiment, the companion NEN is Nb.BtsI. All Nb.BtsI and Nt.BstNBI reactions were performed based on the manufacturer's instructions. All DNA sequences are available in **Table S4**. M: 1 kb dsDNA ladder.

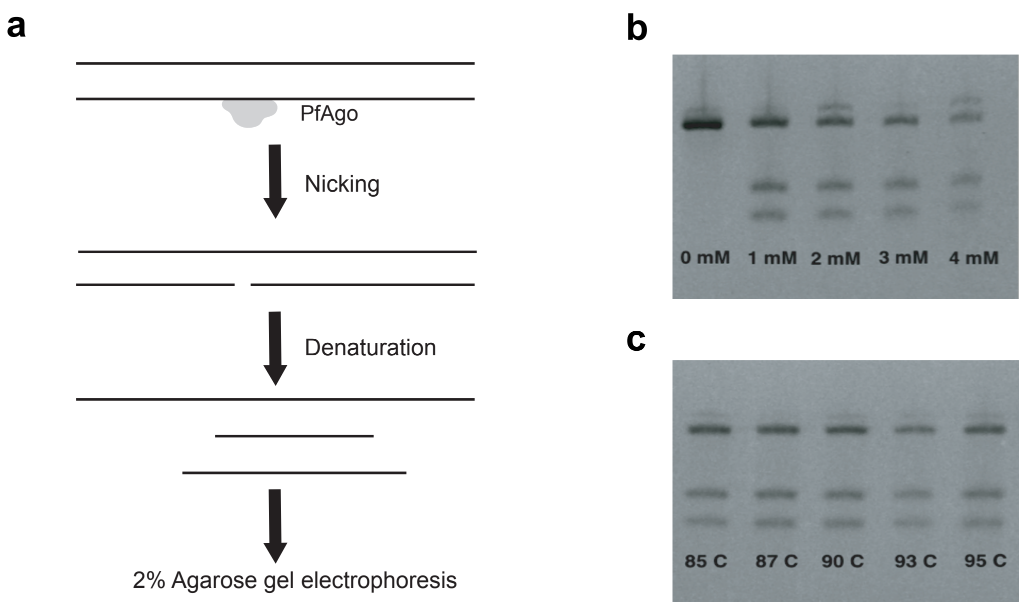


**Figure S2 | ssDNA Agarose gel analysis. a)** The process of sample preparation for ssDNA analysis on 2% agarose gel. **b,c)** The same 450 bp piece of linear DNA is nicked at one site via *Pf*Ago at different temperatures (85 °C to 95 °C) and in different buffers ([$MnCl_2$] = 0,1,2,3,4 mM). The upper bands are the 450 nt ssDNA and the others are 273 nt and 177 nt nicking products.

**3. Comparison of *Pf*Ago and Cas9n nicking activity**

To further explore the potential ability of *Pf*Ago as a writing tool for native DNA-based data storage, we focused on demonstrating its ability to create multiple nicks on the same piece of dsDNA. In all nicking experiments with Cas9n, we used the SpCas9 D10A nickase purchased from Integrated DNA Technologies (IDT) (Alt-R® S.p. Cas9 D10A Nickase). The crRNAs were designed via IDT's Custom Alt-R® CRISPR-Cas9 guide RNA design tool. Both crRNAs and tracrRNAs were purchased from IDT and hybridized based on the manufacturer's protocol. We used a 10X Cas9 reaction buffer: 200 mM HEPES, 1M NaCl, 50 mM $MgCl_2$, 1 mM EDTA, pH 6.5. All Cas9n nicking reactions were set up based on the manufacturer's protocol and were performed at 37 °C for 60 min. The ssDNA gel analysis was performed on an 2% agarose gel. Nicked dsDNA samples were first denatured at high temperature (99 °C) for 5-10 min, followed by immediate cooling to 4 °C. The ssDNA products were then run on pre-made 2% agarose gel (Invitrogen). All nicked PCR products (whether via *Pf*Ago or Cas9n) were then purified using the Qiaquick PCR purification kit (QIAGEN).

On a 450 bp linear DNA (our register for LGA and LMI recording), amplified from the *E. coli* genomic DNA, we chose three nicking sites for *Sp*Cas9n with highest on-target scores (based on IDT's Custom Alt-R® CRISPR-Cas9 guide RNA design tool). Along with these sites, we designed three nicking sites nearby for *Pf*Ago. We also selected 3 sites for nicking with *Pf*Ago, on another ~450 bp PCR product of *E. coli* genomic DNA (which is also the DNA used for toehold creation as discussed below) and designed three *Sp*Cas9n near the mentioned sites (Figure S3a, b). All gDNAs, crRNAs, and tracrRNA were ordered from IDT (the sequences are available in Table S5). Both crRNAs and tracrRNA were hybridized based on the manufacturer's instructions, while the Cas9n reactions were performed based on the manufacturer's protocol described elsewhere (4).

Both versions of the nicked products were sequenced via MiSeq. MiSeq reads were aligned to the reference sequence through the steps described in the Methods section. The final insert size distribution and coverage plots can be found in Figure S3c, d. While in both cases *Pf*Ago showed consistently higher precision of DNA cleavage, *Sp*Cas9n performed as good as *Pf*Ago in the first case, but significantly underperformed in the second case.

Table S2 provides a comparison between the two enzymes based on their specificity, guide type and length, and enzymatic activity. Other than having higher levels of activity on the same substrate and a 16-fold higher versatility (since it does not need a PAM sequence for target binding), *Pf*Ago is also considerably more efficient in terms of cost because the enzyme has a high yield (~200 nmol/L) and its gDNAs can be obtained for a much lower cost (~50 times) than Cas9n's gRNAs. Enzymatic activity and specific activity are calculated by considering the complete nicking of 3.6 pmols (1 µg) of the 450 bp PCR product in the proper reaction time (10 min for *Pf*Ago and 60 min for Cas9n) and sufficient amount of enzyme (3.75 pmols vs. 20 pmols) (Figure S4).

**a**

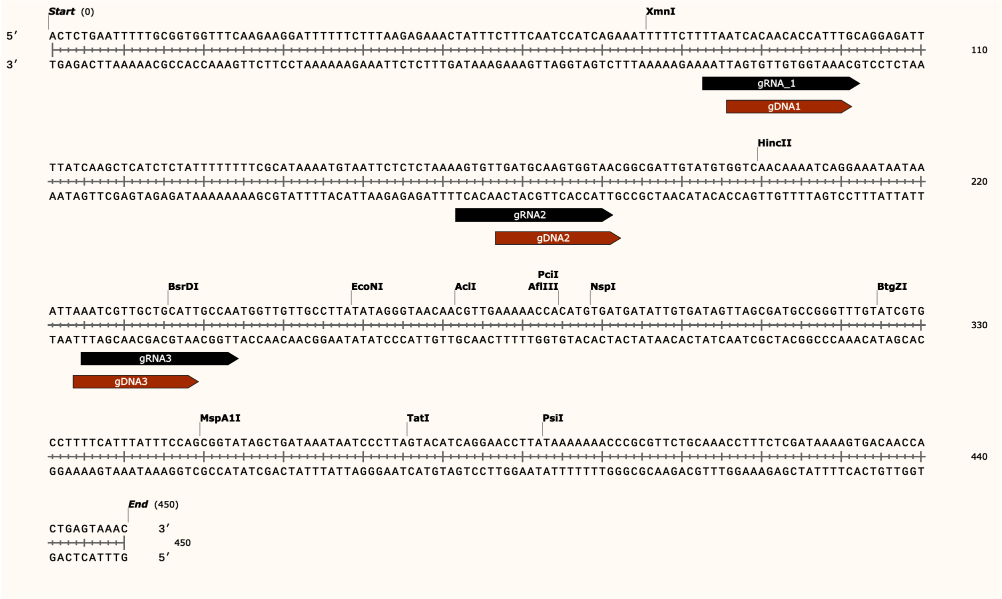
Start (0)
XmnI
HincII
BsrDI
EcoNI
AclI
PciI
AflIII
NspI
BtgZI
MspA1I
TatI
PsiI
End (450)
gRNA_1
gDNA1
gRNA2
gDNA2
gRNA3
gDNA3
110
220
330
440
450

**b**

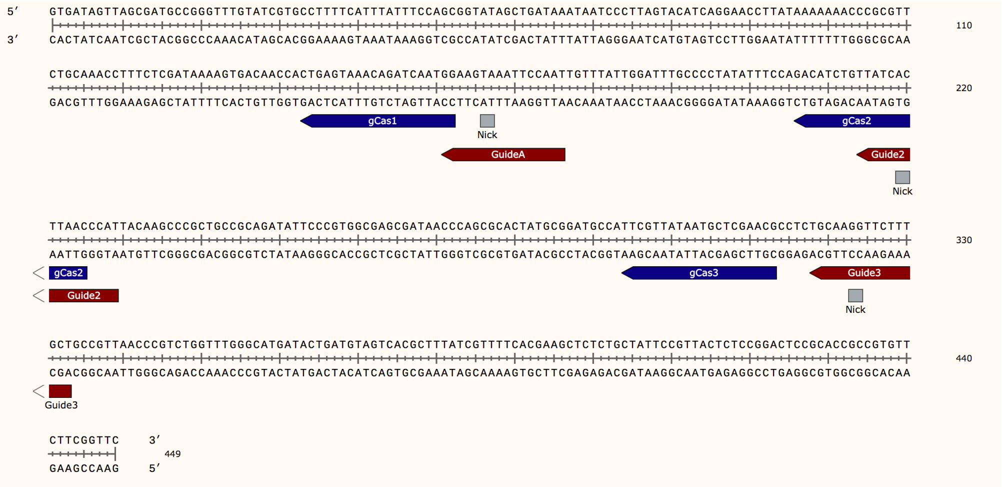
gCas1
gCas2
gCas3
GuideA
Guide2
Guide3
Nick
110
220
330
440
449

c

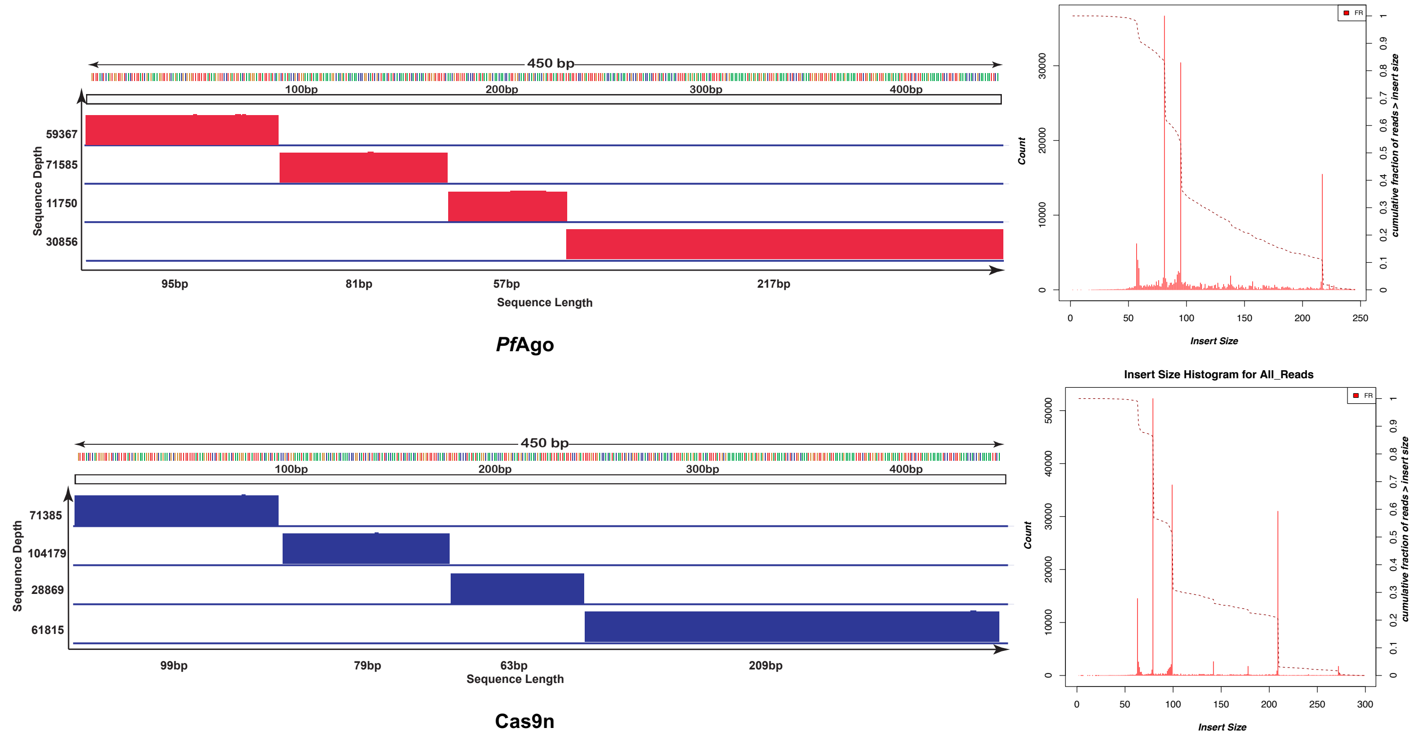
450 bp
100bp
200bp
300bp
400bp
Sequence Depth
59367
71585
11750
30856
95bp
81bp
57bp
217bp
Sequence Length
*Pf*Ago
Insert Size Histogram for All_Reads
Count
Insert Size
cumulative fraction of reads > insert size
FR
450 bp
100bp
200bp
300bp
400bp
Sequence Depth
71385
104179
28869
61815
99bp
79bp
63bp
209bp
Sequence Length
Cas9n
Insert Size Histogram for All_Reads
Count
Insert Size
cumulative fraction of reads > insert size
FR

d

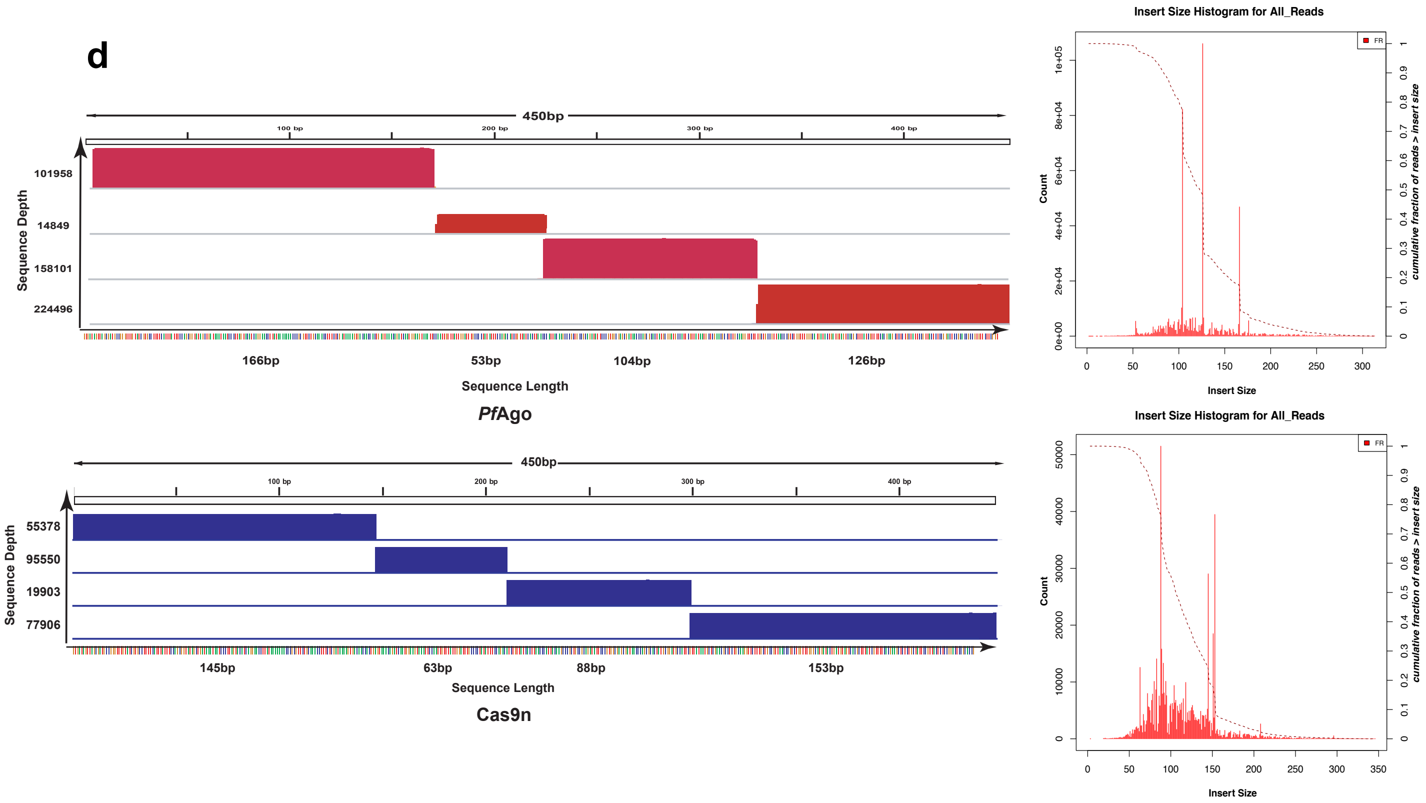
450bp
100 bp
200 bp
300 bp
400 bp
Sequence Depth
101958
14849
158101
224496
166bp
53bp
104bp
126bp
Sequence Length
*Pf*Ago
Insert Size Histogram for All_Reads
Count
Insert Size
cumulative fraction of reads > insert size
FR
450bp
100 bp
200 bp
300 bp
400 bp
Sequence Depth
55378
95550
19903
77906
145bp
63bp
88bp
153bp
Sequence Length
Cas9n
Insert Size Histogram for All_Reads
Count
Insert Size
cumulative fraction of reads > insert size
FR

**Figure S3 | *Pf*Ago vs. Cas9n.**

**a, b**) Sequence and map of the selected substring of the *E. coli* genome (LGA and SEQ1). Note that the gDNAs, crRNAs, nicks and their corresponding positions are marked.

**c, d)** MiSeq output coverage plots (left) and insert size histograms (right) of the nicked products (LGA and SEQ1, respectively), via *Pf*Ago and *Sp*Cas9n D10A.

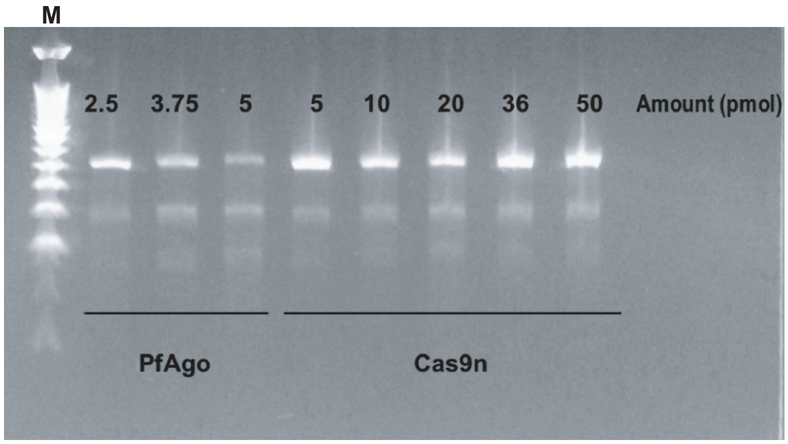


**Figure S4 |** To further evaluate the enzymatic activity of *Pf*Ago vs Cas9n D10A, we nicked 1 μg of the 450 bp substrate DNA at one site, with different ratios of enzyme to substrate. The final products were analyzed on 2% agarose gel. *Pf*Ago showed efficient nicking with only 3.75 pmols of the enzyme used. Based on most protocols and publications on *in vitro* nicking via SpCas9n, a ratio of 10:10:1 (gRNA: enzyme: DNA substrate) is needed for efficient nicking, which agrees with the gel results shown above.
M: 50bp DNA ladder.

**Table S2 | Comparing *Pf*Ago vs *Sp*Cas9n.** Unlike conventional restriction/nicking endonucleases, neither of the two enzymes (Cas9n and *Pf*Ago) requires a specific recognition site. However, Cas9n requires the presence of a PAM sequence on the 3' site of the target sequence, while *Pf*Ago does not need such a special recognition sequence. Furthermore, *Pf*Ago takes short DNA guides which are cheaper, more stable, and easier to handle and store; Cas9n uses gRNAs, which can be transcribed *in vitro* or prepared by hybridizing crRNA (36 nt) and tracrRNA (~67 nt). Both are more time-consuming and expensive procedures than their *Pf*Ago counterparts. To efficiently nick at least one site for 1 μg of our 450 bp register (~3.6 pmol), it suffices to use 3.75 pmols of *Pf*Ago (~1:1 enzyme to substrate ratio), while based on most protocols for Cas9n *in vitro* nicking, nicking the same amount of DNA with Cas9n on at least one site requires as much as 10-folds more enzyme, i.e., around 36 pmol for sufficiently efficient nicking. Our results show that roughly 20 pmol of Cas9n suffices for equally efficient nicking compared to *Pf*Ago, which is still >5 times the amount needed for *Pf*Ago**.**

| **Enzyme type** | **Context specificity** | **PAM requirement** | **Guide type/Length** | **Cost** | **Enzyme: substrate ratio*** | **Specific activity (μmol min$^{-1}$ mg$^{-1}$)** | **Turnover** | **Enzymatic activity (μmol min$^{-1}$)** |
|---|---|---|---|---|---|---|---|---|
| *Pf*Ago | **-** | - | DNA/16 nt | low | ~1 | $1\times10^{-3}$ | Multiple | $3.6\times10^{-7}$ |
| *Sp*Cas9n | **-** | + | RNA/~100 nt | high | >5 | $1.8\times10^{-5}$ | Single | $6\times10^{-8}$ |

***** For nicking one site on 1 μg (3.6 pmols) of the register**.**

#### 4. Lincoln's Gettysburg Address (LGA) and Lincoln's Memorial Image (LMI) recording

To test our DNA storage method, we selected a 450 bp DNA sequence from the *E. coli* K12 MG1655 genome as the register for both LGA and LMI recording. For simplicity, and with a slight abuse of naming convention, we henceforth use LGA to describe the chosen register. All (ten) gDNAs (red) are designed based on the rules described in the Methods section and all nicks (black) occur on the sense strand, between positions 10 and 11 of the corresponding guide (Figure S5-6). The full binary encoding of the LGA and LMI data is available in the Supplementary files.

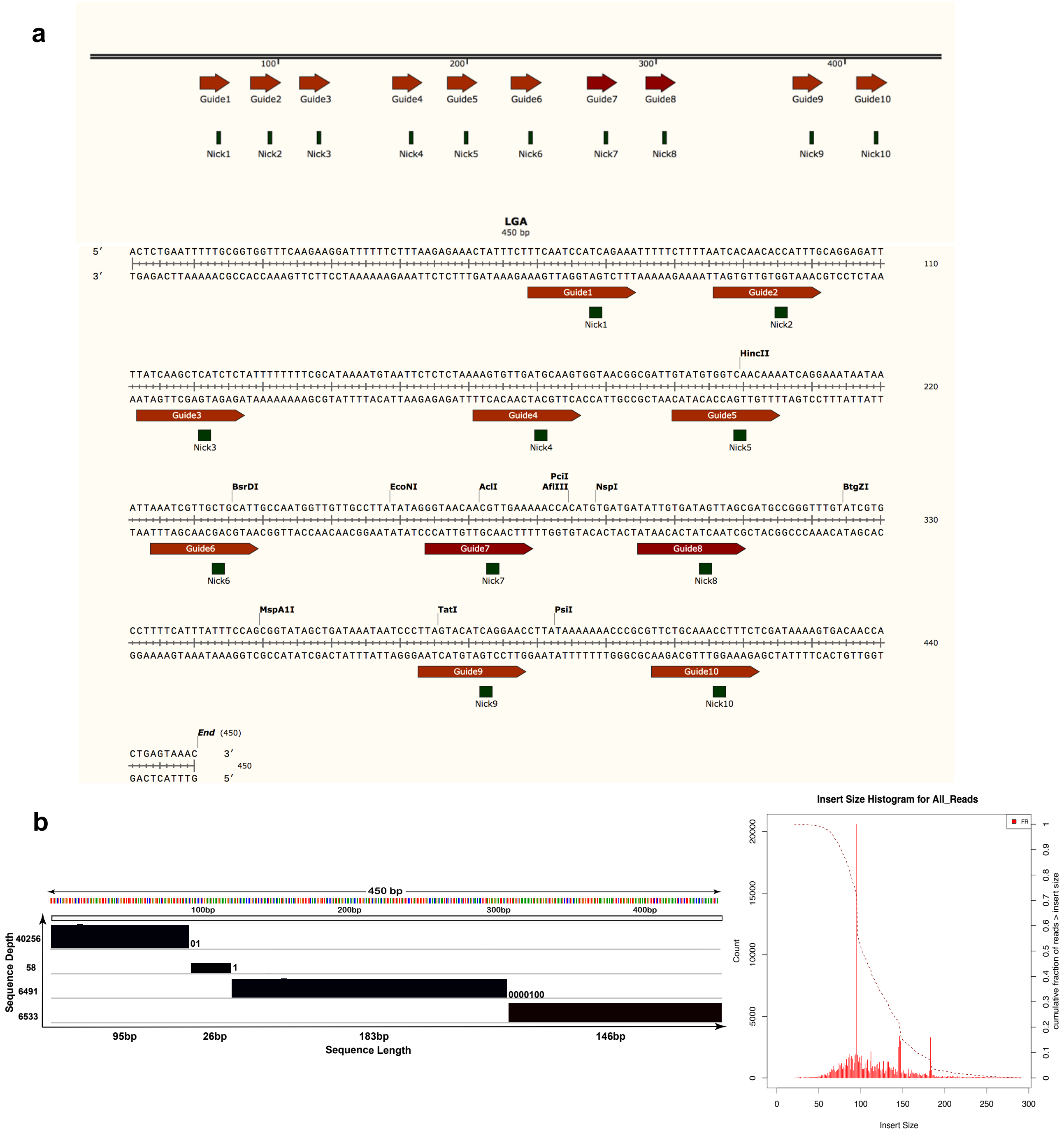

**c**

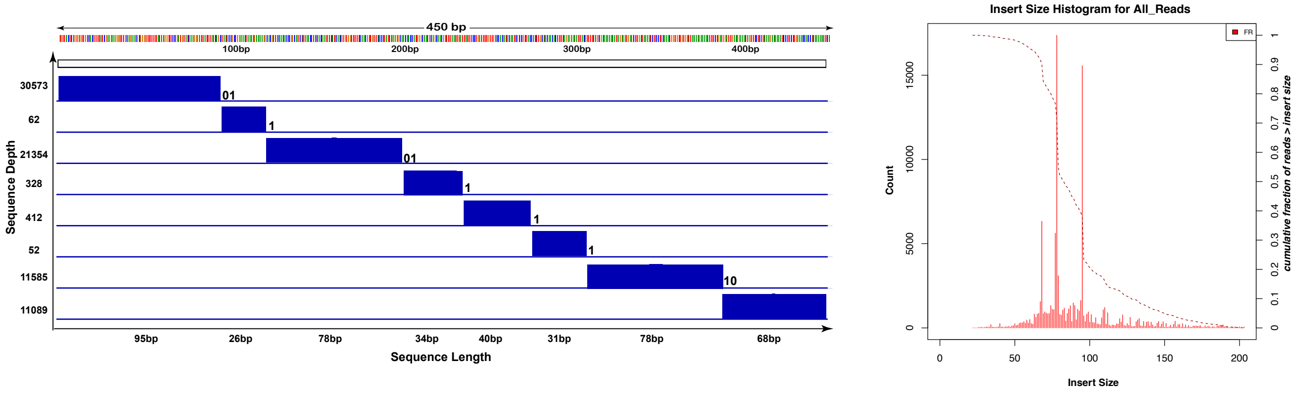


**d**

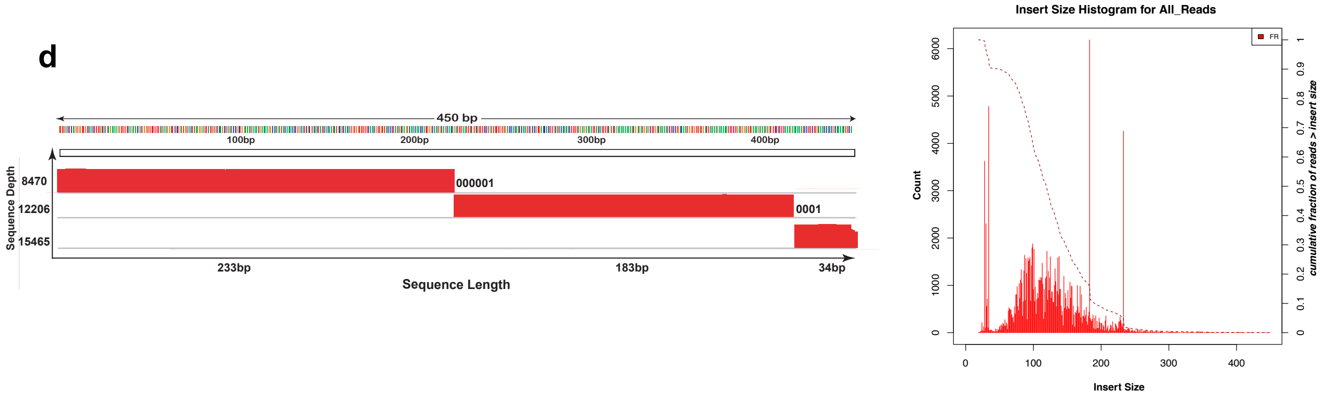


**Figure S5 | LGA readout results.** MiSeq reads analysis of three randomly chosen spatially adjacent registers after recording. **a**) The map and sequence content of the 450 bp register. Designed guides (red) and their positions are shown along with the selected nicking sites (black). Based on the sequenced (retrieved) reads and their lengths, one is able to detect the nicked sites and also the sites which are not nicked and therefore, reconstruct the 10-bit string recorded in each register. The binary strings encoded in the 10-bit registers are as follows: **b**) 0110000100, **c**) 0110111110, **d**) 0000010001.

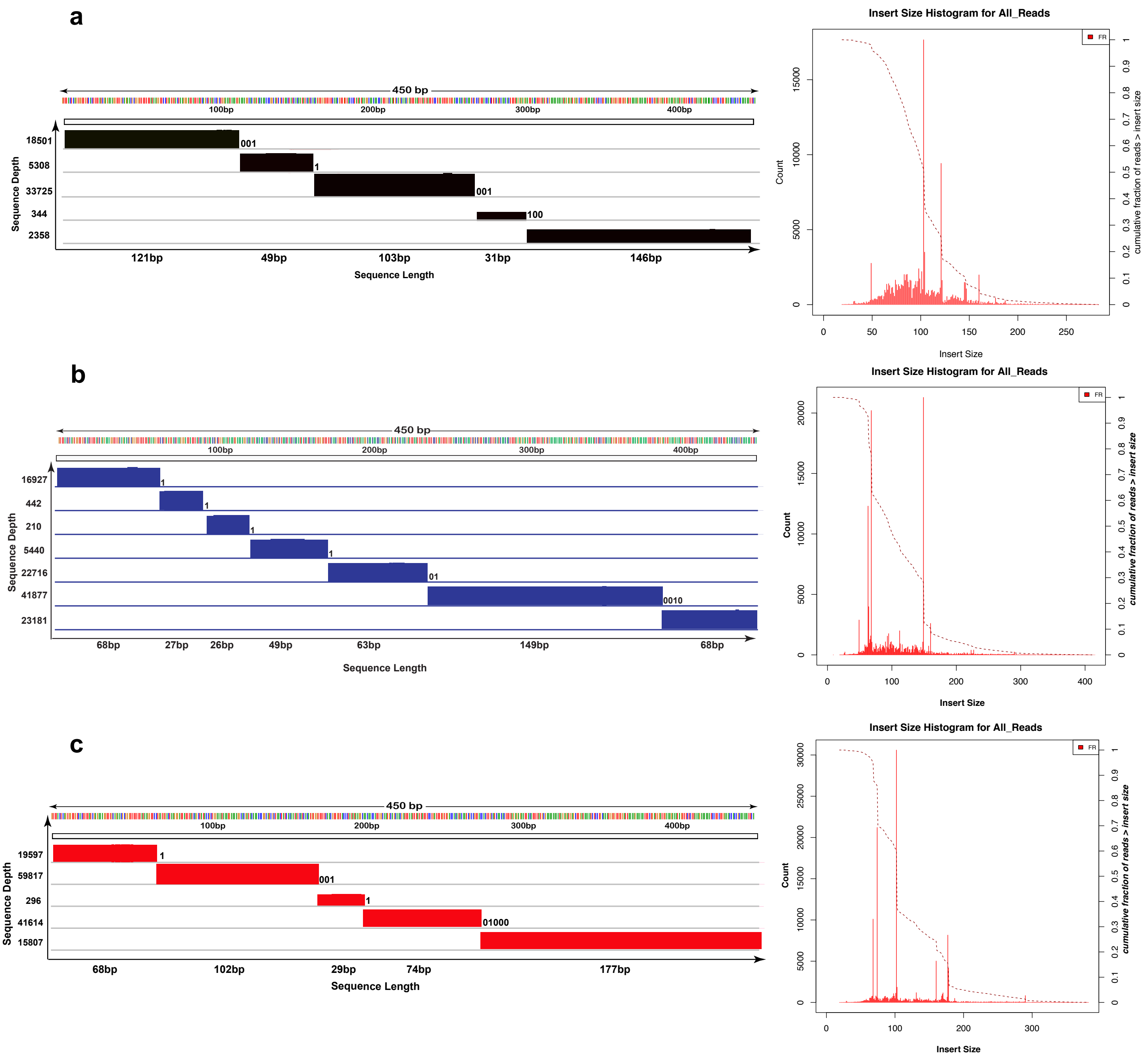
a
450 bp
100bp
200bp
300bp
400bp
Sequence Depth
18501
5308
33725
344
2358
001
1
001
100
121bp
49bp
103bp
31bp
146bp
Sequence Length
Insert Size Histogram for All_Reads
FR
Count
Insert Size
cumulative fraction of reads > insert size
b
450 bp
100bp
200bp
300bp
400bp
Sequence Depth
16927
442
210
5440
22716
41877
23181
1
1
1
1
01
0010
68bp
27bp
26bp
49bp
63bp
149bp
68bp
Sequence Length
Insert Size Histogram for All_Reads
FR
Count
Insert Size
cumulative fraction of reads > insert size
c
450 bp
100bp
200bp
300bp
400bp
Sequence Depth
19597
59817
296
41614
15807
1
001
1
01000
68bp
102bp
29bp
74bp
177bp
Sequence Length
Insert Size Histogram for All_Reads
FR
Count
Insert Size
cumulative fraction of reads > insert size

**d**

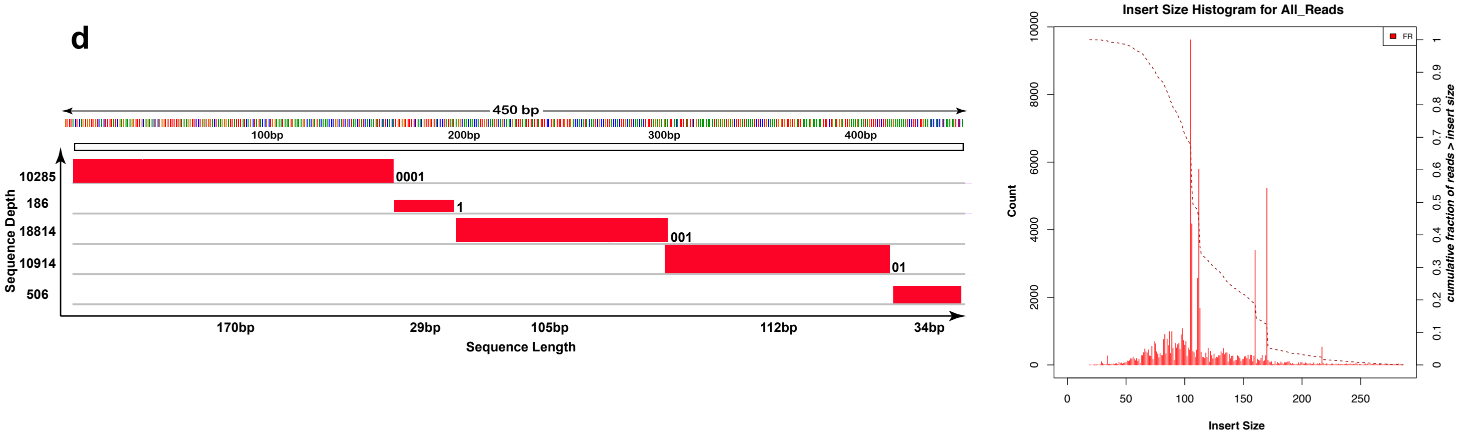


**e**

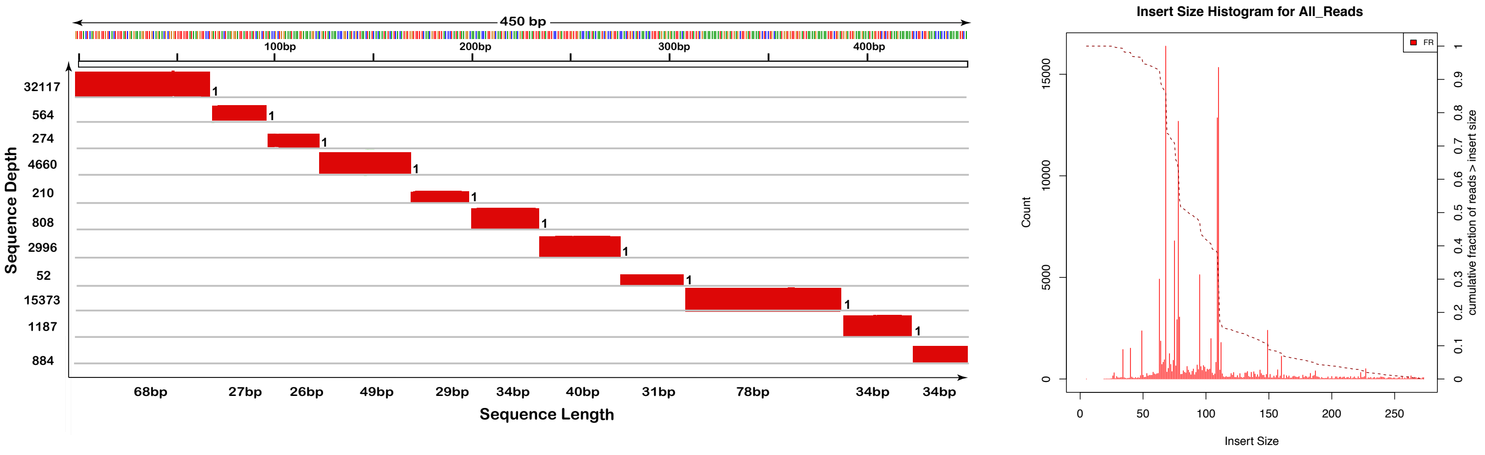


**Figure S6 | LMI readout results**. The MiSeq read analysis of five randomly chosen consecutive registers after recording the LMI binary data file into native DNA. Based on the sequenced (retrieved) reads and their lengths, one can detect the nicked sites as well as the sites that were not nicked and hence, reconstruct the 10-bit string stored in each register. The retrieved register contents equal **a)** 0011001100, **b)** 1111010010, **c**) 1001101000, **d**) 0001100101, **e**) 1111111111

**Table S3|** The files of size 14.4 KB created from the combinatorial library of 1024 nicked native DNA registers. One file is a text file, while the other is an image. The number of different 10-bit strings appearing in the files is shown in the third column.

| Data | File size | Number of different 10-bit strings |
|---|---|---|
| Lincoln's Gettysburg address (Text) | 14 KB | 129 |
| Lincoln's memorial (Image) | 0.4 KB | 1015 |
| Total | 14.4 KB | 1024 |

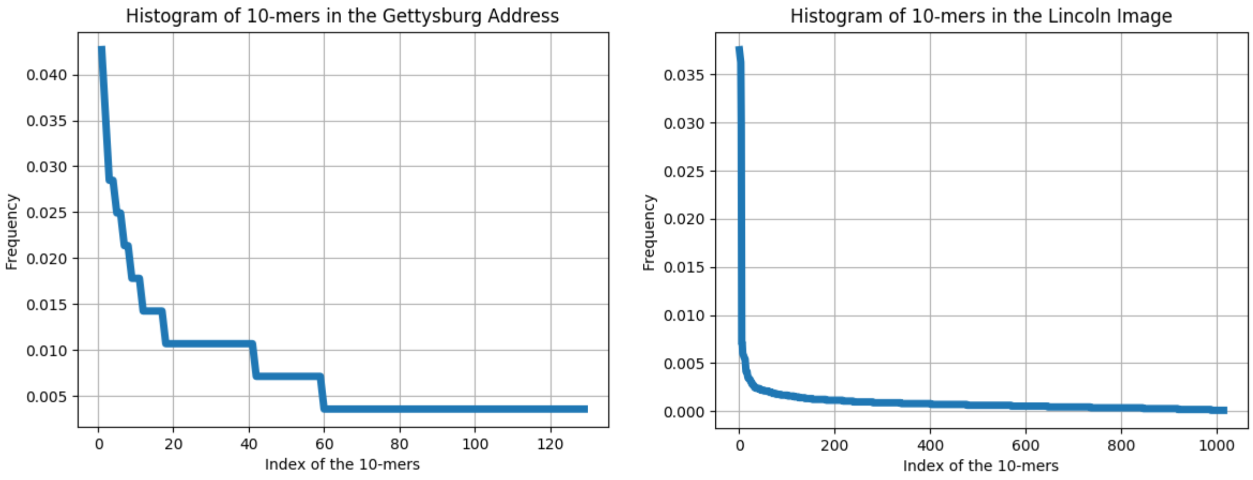


**Figure S7 |** The distribution of all distinct 10-bit strings in the ASCII code of the Gettysburg address (**left**) and the Lincoln memorial image (**right**). In total, 129 and 1015 different 10-mers were present in the text and image file, respectively. Note that for image storage, one may use quantization methods that represent two or more different k-mers by the same recorded nick position. This may lead to subjective changes in the quality of the image but ensure additional savings in cost.

#### 5. Data organization

There are several important conceptual differences between DNA storage and classical recording platforms. In the former case, the storage media (DNA) is simultaneously the recorded data. There is no natural organization of the data as it is stored as a pool of oligos, with addresses that usually require a 10% coding overhead (3). In the latter case, the storage media is a substrate such as tape, disk or array of chargeable cells:

https://en.wikibooks.org/wiki/Introduction_to_Computer_Information_Systems/Storage.

Data is recorded by changing some physical or chemical property of the substrate, such as magnetization or surface structure. Furthermore, data is organized into units of bits that allow for ease of access and/or rewriting (such as blocks, sectors, tracks). Our storage architecture based on native DNA departs from the synthetic DNA storage practice and more closely matches the properties of existing recorders in so far that:

- Native DNA is the storage media, but user content is not hard-wired into the media. Instead, native DNA is used a "punch card" for recording user-defined content. Furthermore, mixtures of shorter register sequences allow for parallel recording on multiple registers, and if the experiments are properly designed, one enzyme/guide pair may be used to record bits in different registers provided that the nicking sites have small sequence edit distance.
- There exists an inherent ordering of the data content dictated by the ordering of the register sequences within the *E. coli* genome. Additional ordering is provided by arranging the DNA registers or mixture of registers into arrays that enable faster retrieval of information and additional structural organization. This array structure resembles the DNA microarray platforms used for synthesizing DNA oligos, except for the fact that DNA does not need to be affixed to the substrate. In addition, although both systems perform data recording within spots/wells, punch card native DNA recording only needs to store the non-zero values and does not have to follow an iterative chain growth protocol as nicks may be created in parallel.
- The volume of native DNA created during a nicking reaction may be orders of magnitude larger than that obtained with classical synthesis. This allows for a large number of readout cycles, and represents a feature shared with modern silicon recorders.

The above described system properties are illustrated in the figure below. When a well contains one register only, all registers contain the same nicking pattern **(a)** but when multiple sequences (registers) are located in a reaction well **(b),** each register type can contain different data. Hence, the second architecture allows for increasing the vial/per-spot storage density.

a b

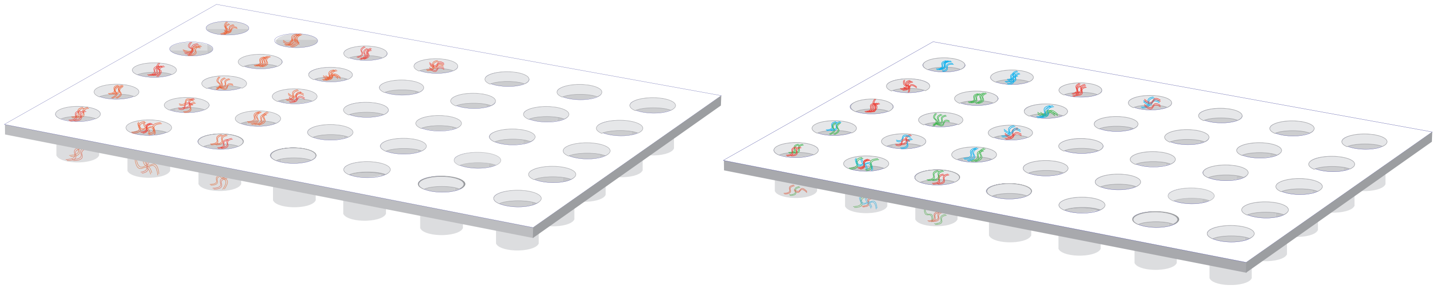

A comparison of a microspot architecture for nick-based storage (left) and the DNA microarray synthesis platforms (right) used by most commercial DNA vendors is illustrated below. Major system design advantages of our storage method include:

- There is no need to fixate primers on the spots. Native DNA used as the nicking substrate may be loaded on the spots without attaching it to the surface.
- There is no need for masking and washing of the glass substrate between two base incorporations. The nicking enzymes with their corresponding guides are added only once at the beginning of the recording process.
- There is no need for sequential incorporation of basis, as all nicking reactions take place simultaneously.

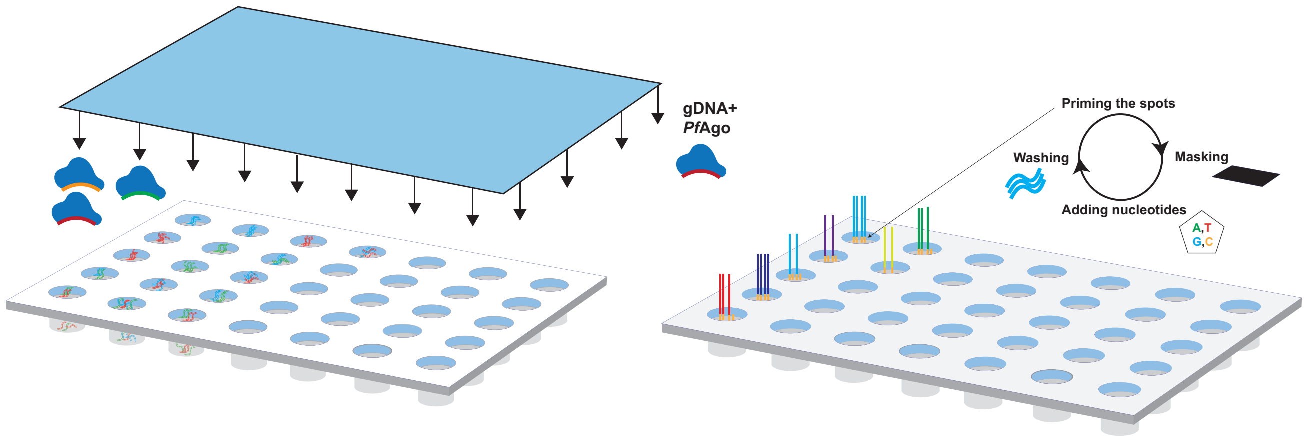

#### 6. Approaches to increase the information density

*Sense and antisense nicking*

A higher information density can be achieved with recording data on both (sense and antisense) strands. In that setting, the binary format is switched to ternary format, i.e., for any predesignated nicking position, a nick on the sense strand denotes 1, a nick on the antisense denotes 2, and no nick denotes 0. Thus, for every nicking site the potential storable data is increased ~1.58 fold. We experimentally showed that the same 450 bp register (LGA) can be nicked on 10 prescribed sites on both strands, six of which are located on the sense strand and four on the antisense strands, corresponding to the 10-bit string of data: 1212121211 (Figure S8). No negative enzyme interactions were noticed, nor did the sense-antisense nick cause issues with DNA instability. The sense-antisense nicking approach may also be used for parallel recording on multiple registers.

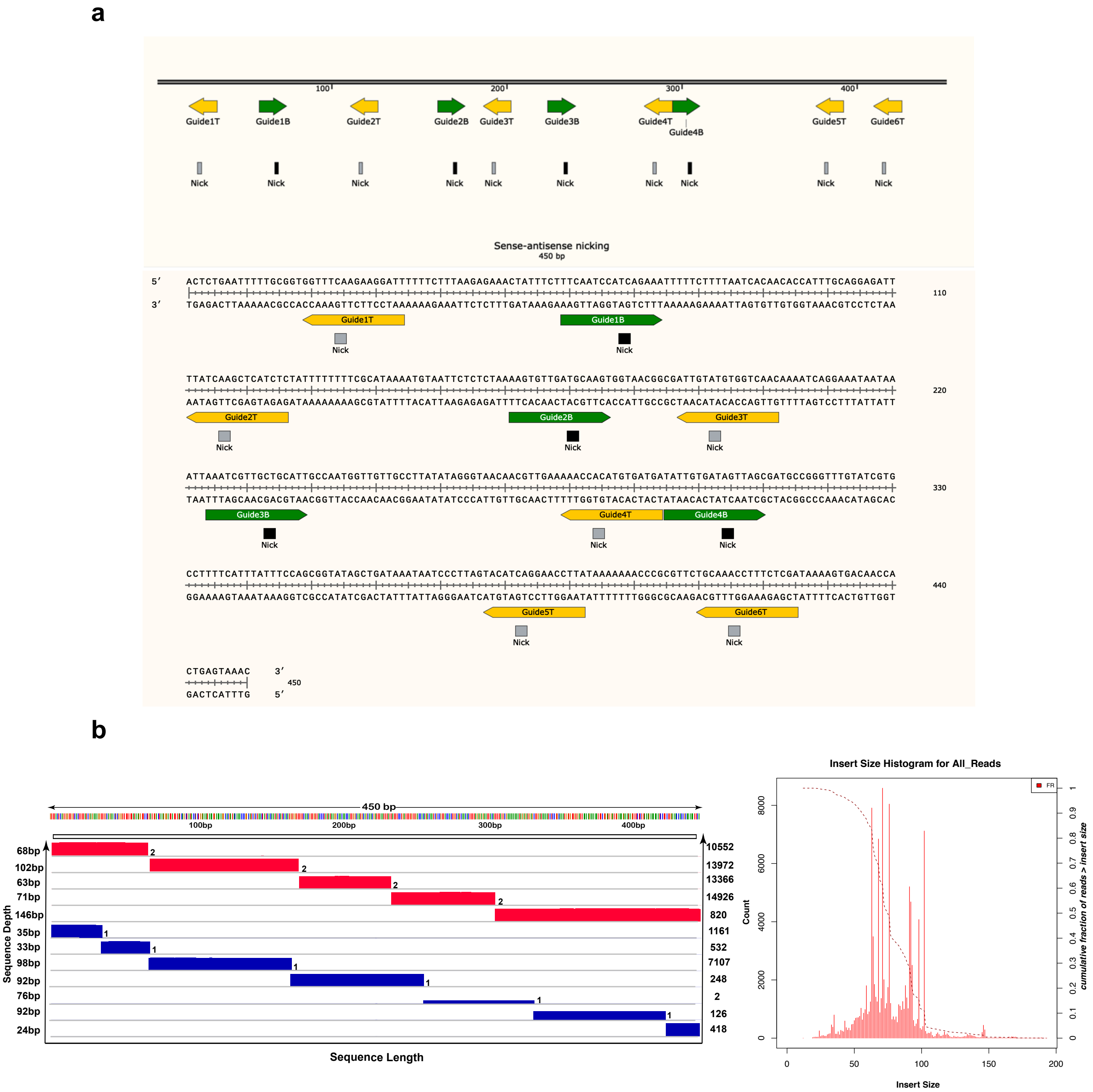


**Figure S8 | Simultaneous nicking of both sense and antisense strands**. **a)** The map and the sequence content of the 450 bp register. The gDNAs are colored in yellow (targeting the sense strand) and green (targeting the antisense strand). Sense strand nicking positions are shown in gray while antisense nicking positions are shown in black. **b)** Insert size distribution and coverage plots of the ssDNAs sequenced via MiSeq. Each ssDNA fragment is separately aligned to the two reference sequences (sense and antisense) and assigned to the position with highest coverage.

*Multi-register recording; maps and sequences*

Another means for increasing information density is to use multiple orthogonal registers. Here, we performed random search to identify four additional register sequences of length 450 bp (orthogonal sequences 1,2,3 and 4; Figure S9a), from the same source (*E. coli* K12 MG1655 genome). All sequences have low pairwise similarities (<50%) with respect to each other. Then, we determined nicking positions and designed gDNAs for all new registers (Figure S9b, c, d and e), as well as all five registers, including LGA (Figure S5a). A total number of 32 nicking positions was used for recording. As a proof of concept, we stored the title "Gettysburg Address" of size 126 bits in our final five registers and successfully recalled it without errors (Figure S10).

To identify the orthogonal register sequences, we used the software Needle (https://www.ebi.ac.uk/Tools/services/web/toolresult.ebi?jobId=emboss_needle-I20181031-164650-0561-47446167-p2m). The alignment results are listed below. Alternative alignment software options include CLUSTAL OMEGA (https://www.ebi.ac.uk/Tools/msa/clustalo/) and MUSCLE (https://www.ebi.ac.uk/Tools/msa/muscle/).

Alignment score, LGA and O1

```
#=======================================
#
# Aligned_sequences: 2
# 1: EMBOSS_001
# 2: EMBOSS_001
# Matrix: EDNAFULL
# Gap_penalty: 10.0
# Extend_penalty: 0.5
#
# Length: 586
# Identity:     244/586 (41.6%)
# Similarity:   244/586 (41.6%)
# Gaps:         272/586 (46.4%)
# Score: 371.0
#
#=======================================
```

Alignment score, LGA and O2

```
#=======================================
#
# Aligned_sequences: 2
# 1: EMBOSS_001
# 2: EMBOSS_001
# Matrix: EDNAFULL
# Gap_penalty: 10.0
# Extend_penalty: 0.5
#
```

### Length: 587
# Identity:     228/587 (38.8%)
# Similarity:   228/587 (38.8%)
# Gaps:         274/587 (46.7%)
### Score: 319.0
#
#=======================================
Alignment score, LGA and O3
#=======================================
#
### Aligned_sequences: 2
### 1: EMBOSS_001
### 2: EMBOSS_001
### Matrix: EDNAFULL
### Gap_penalty: 10.0
### Extend_penalty: 0.5
#
### Length: 610
# Identity:     218/610 (35.7%)
# Similarity:   218/610 (35.7%)
# Gaps:         320/610 (52.5%)
### Score: 301.5
#
#=======================================
Alignment score, LGA and O4
#=======================================
#
### Aligned_sequences: 2
### 1: EMBOSS_001
### 2: EMBOSS_001
### Matrix: EDNAFULL
### Gap_penalty: 10.0
### Extend_penalty: 0.5
#
### Length: 596
# Identity:     235/596 (39.4%)
# Similarity:   235/596 (39.4%)
# Gaps:         292/596 (49.0%)
### Score: 320.5

**a**

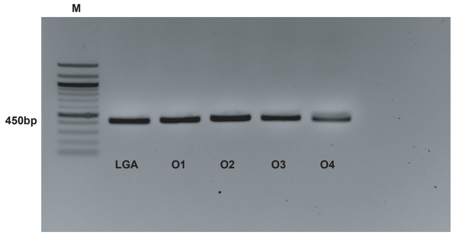
M
450bp
LGA
O1
O2
O3
O4

**b**

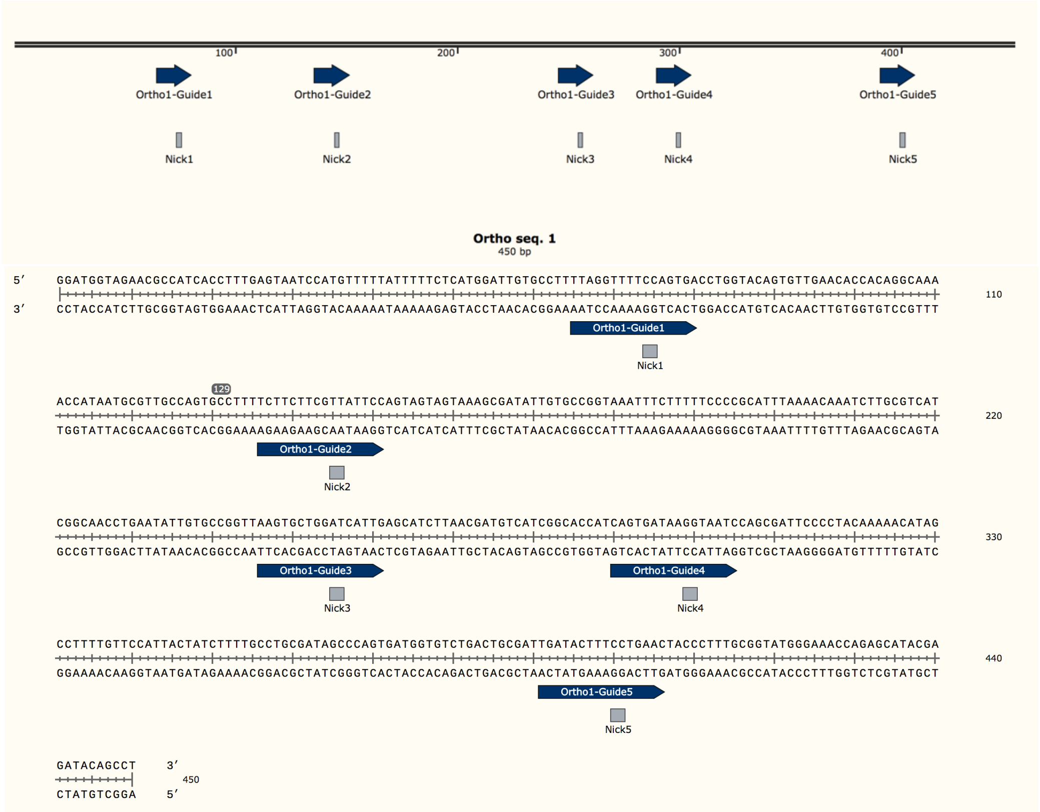
100
200
300
400
Ortho1-Guide1
Ortho1-Guide2
Ortho1-Guide3
Ortho1-Guide4
Ortho1-Guide5
Nick1
Nick2
Nick3
Nick4
Nick5
Ortho seq. 1
450 bp
5′ GGATGGTAGAACGCCATCACCTTTGAGTAATCCATGTTTTTATTTTTCTCATGGATTGTGCCTTTTAGGTTTTCCAGTGACCTGGTACAGTGTTGAACACCACAGGCAAA
110
3′ CCTACCATCTTGCGGTAGTGGAAACTCATTAGGTACAAAAATAAAAAGAGTACCTAACACGGAAAATCCAAAAGGTCACTGGACCATGTCACAACTTGTGGTGTCCGTTT
Ortho1-Guide1
Nick1
129
ACCATAATGCGTTGCCAGTGCCTTTTCTTCTTCGTTATTCCAGTAGTAGTAAAGCGATATTGTGCCGGTAAATTTCTTTTTCCCCGCATTTAAAACAAATCTTGCGTCAT
220
TGGTATTACGCAACGGTCACGGAAAAGAAGAAGCAATAAGGTCATCATCATTTCGCTATAACACGGCCATTTAAAGAAAAAGGGGCGTAAATTTTGTTTAGAACGCAGTA
Ortho1-Guide2
Nick2
CGGCAACCTGAATATTGTGCCGGTTAAGTGCTGGATCATTGAGCATCTTAACGATGTCATCGGCACCATCAGTGATAAGGTAATCCAGCGATTCCCCTACAAAAACATAG
330
GCCGTTGGACTTATAACACGGCCAATTCACGACCTAGTAACTCGTAGAATTGCTACAGTAGCCGTGGTAGTCACTATTCCATTAGGTCGCTAAGGGGATGTTTTTGTATC
Ortho1-Guide3
Ortho1-Guide4
Nick3
Nick4
CCTTTTGTTCCATTACTATCTTTTGCCTGCGATAGCCCAGTGATGGTGTCTGACTGCGATTGATACTTTCCTGAACTACCCTTTGCGGTATGGGAAACCAGAGCATACGA
440
GGAAAACAAGGTAATGATAGAAAACGGACGCTATCGGGTCACTACCACAGACTGACGCTAACTATGAAAGGACTTGATGGGAAACGCCATACCCTTTGGTCTCGTATGCT
Ortho1-Guide5
Nick5
GATACAGCCT 3′
450
CTATGTCGGA 5′

**c**

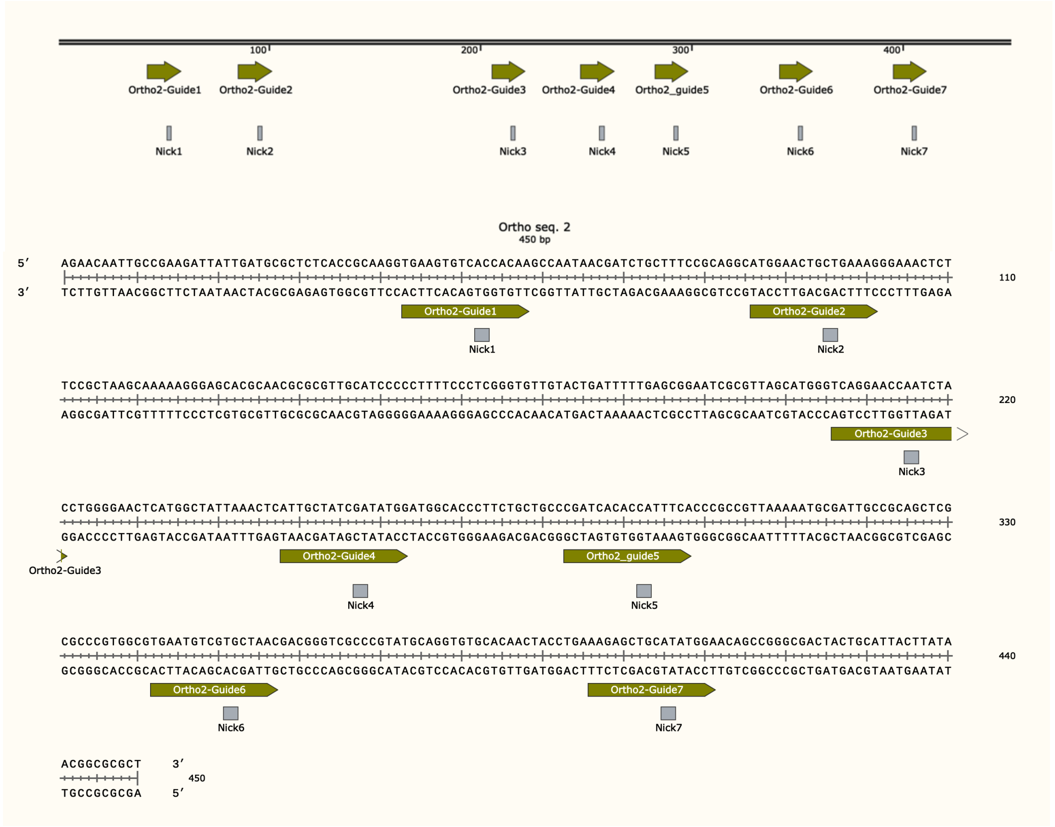

100
200
300
400
Ortho2-Guide1
Ortho2-Guide2
Ortho2-Guide3
Ortho2-Guide4
Ortho2_guide5
Ortho2-Guide6
Ortho2-Guide7
Nick1
Nick2
Nick3
Nick4
Nick5
Nick6
Nick7
Ortho seq. 2
450 bp
5'
3'
110
220
330
440
450


**d**

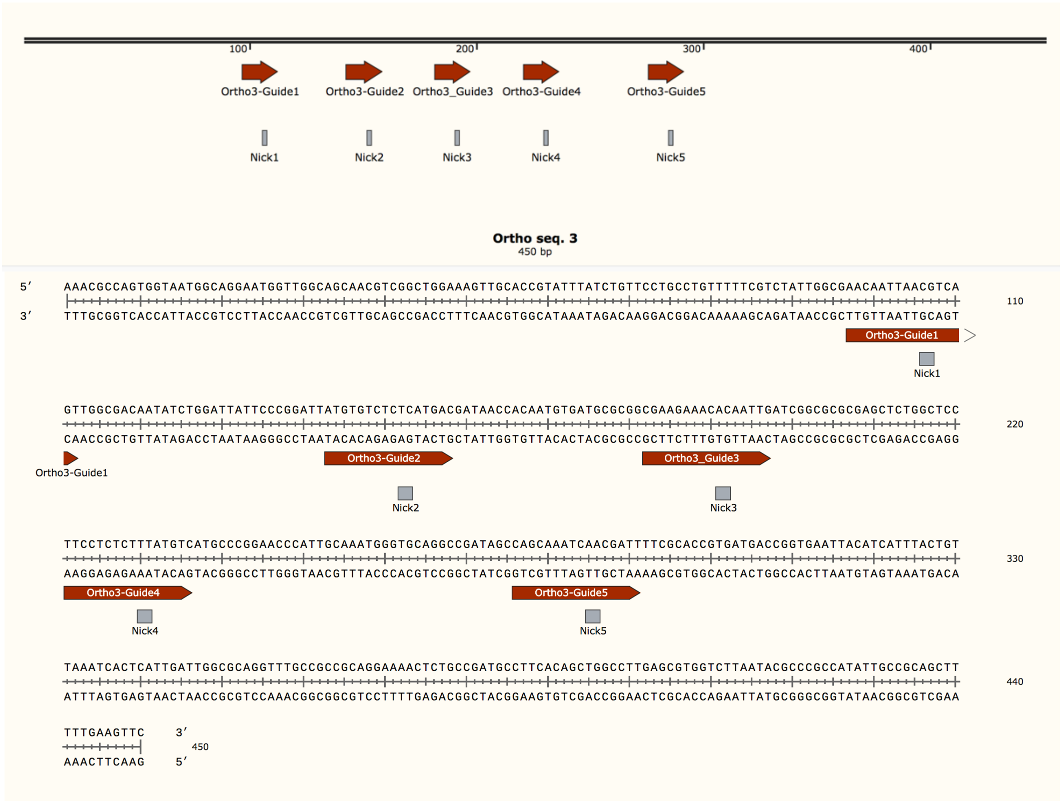

100
200
300
400
Ortho3-Guide1
Ortho3-Guide2
Ortho3_Guide3
Ortho3-Guide4
Ortho3-Guide5
Nick1
Nick2
Nick3
Nick4
Nick5
Ortho seq. 3
450 bp
5'
3'
110
220
330
440
450

**e**

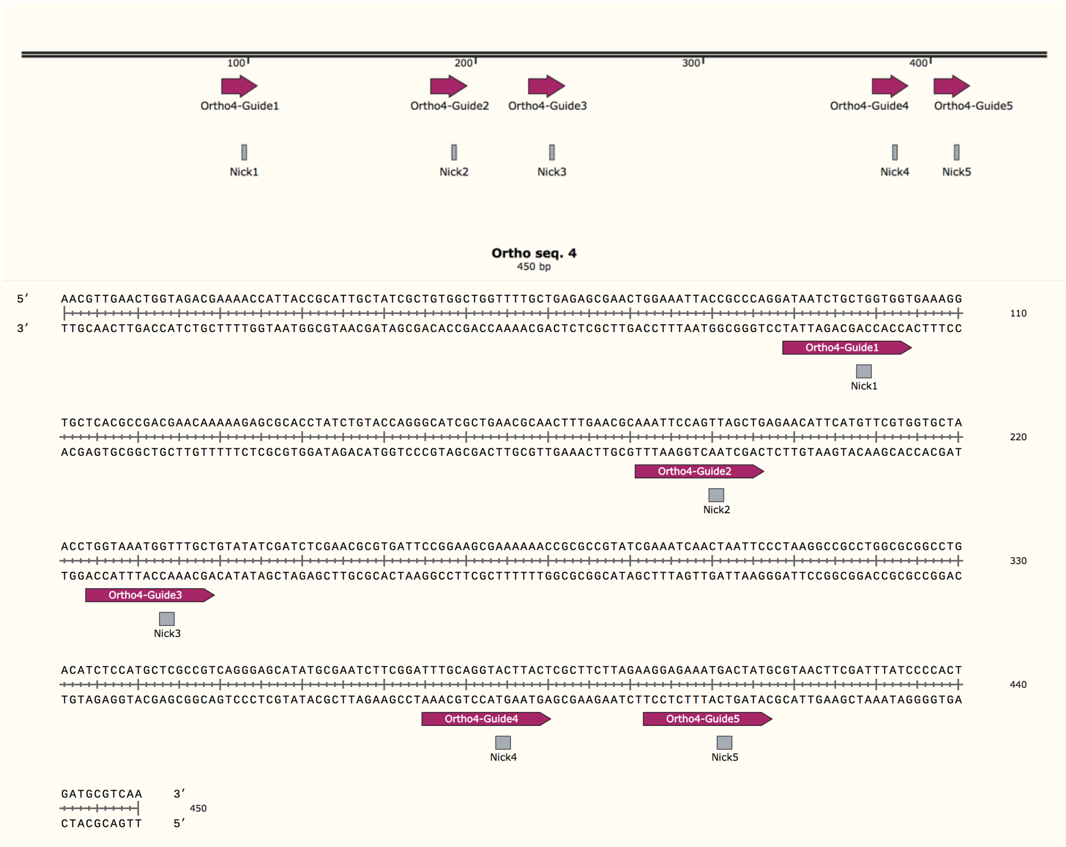


**Figure S9 | Orthogonal registers**. Two registers are termed orthogonal if their edit distance is <50% Such registers do not share substrings with sufficiently similar sequence content that would cause a lack of nicking specificity. **a)** The length of all four added orthogonal registers, along with the previously used LGA register, was verified through gel analysis. All registers were also individually sequenced via Sanger sequencing. **b,c,d and e)** The maps, sequences and selected sequencing results for the four added orthogonal sequences. All gDNAs are indicated by colored arrows, while nicking sites are indicated by gray squares. All nicks were created at the strand shown at the bottom. The registers, LGA, O1, O2, O3 and O4 comprise 10,5,7,5,5 nicking sites (corresponding to the number of bits each is capable of storing), respectively. This amounts to a total of 32 nicking sites, or 32 bits.

**a**

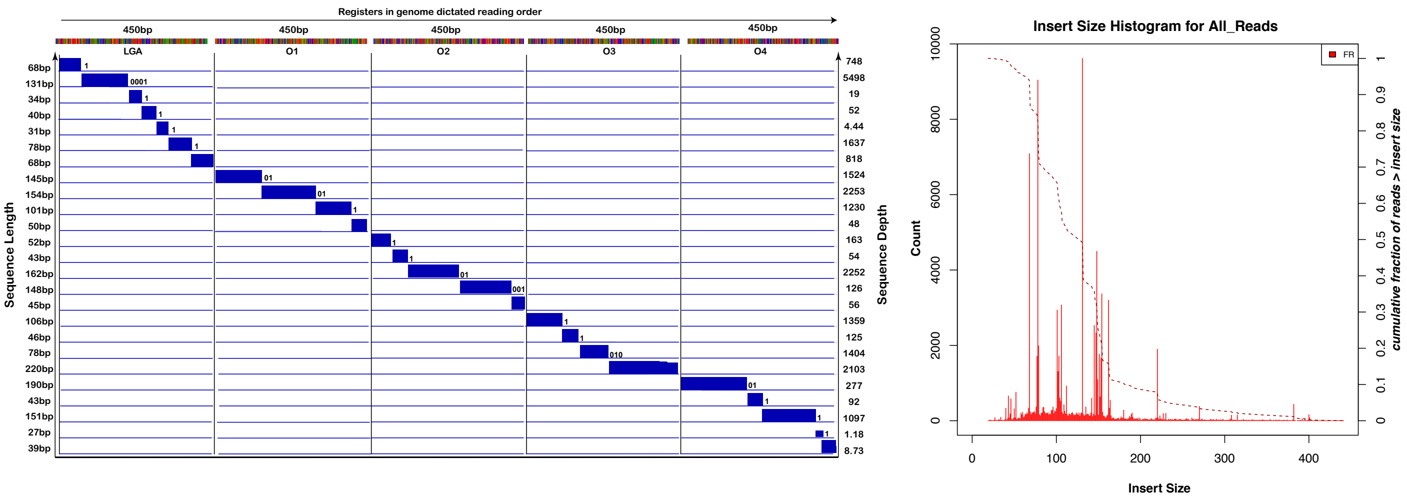
Registers in genome dictated reading order
450bp
LGA
O1
O2
O3
O4
Sequence Length
Sequence Depth
68bp 131bp 34bp 40bp 31bp 78bp 68bp 145bp 154bp 101bp 50bp 52bp 43bp 162bp 148bp 45bp 106bp 46bp 78bp 220bp 190bp 43bp 151bp 27bp 39bp
748 5498 19 52 4.44 1637 818 1524 2253 1230 48 163 54 2252 126 56 1359 125 1404 2103 277 92 1097 1.18 8.73
Insert Size Histogram for All_Reads
FR
Count
Insert Size
cumulative fraction of reads > insert size

**b**

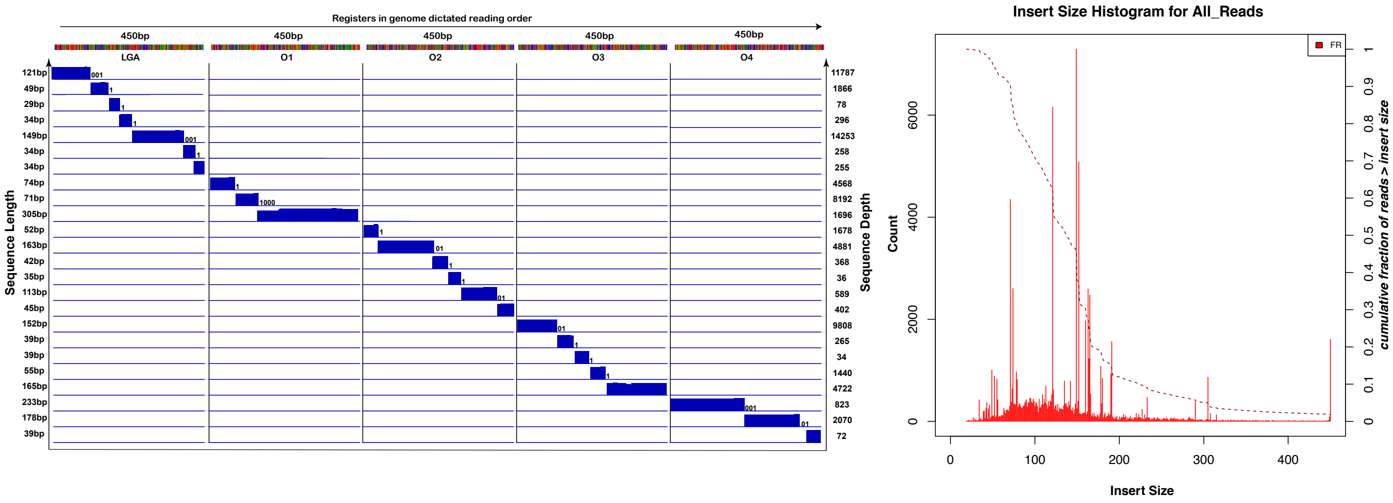
Registers in genome dictated reading order
450bp
LGA
O1
O2
O3
O4
Sequence Length
Sequence Depth
121bp 49bp 29bp 34bp 149bp 34bp 34bp 74bp 71bp 305bp 52bp 163bp 42bp 35bp 113bp 45bp 152bp 39bp 39bp 55bp 165bp 233bp 178bp 39bp
11787 1866 78 296 14253 258 255 4568 8192 1696 1678 4881 368 36 589 402 9808 265 34 1440 4722 823 2070 72
Insert Size Histogram for All_Reads
FR
Count
Insert Size
cumulative fraction of reads > insert size

**c**

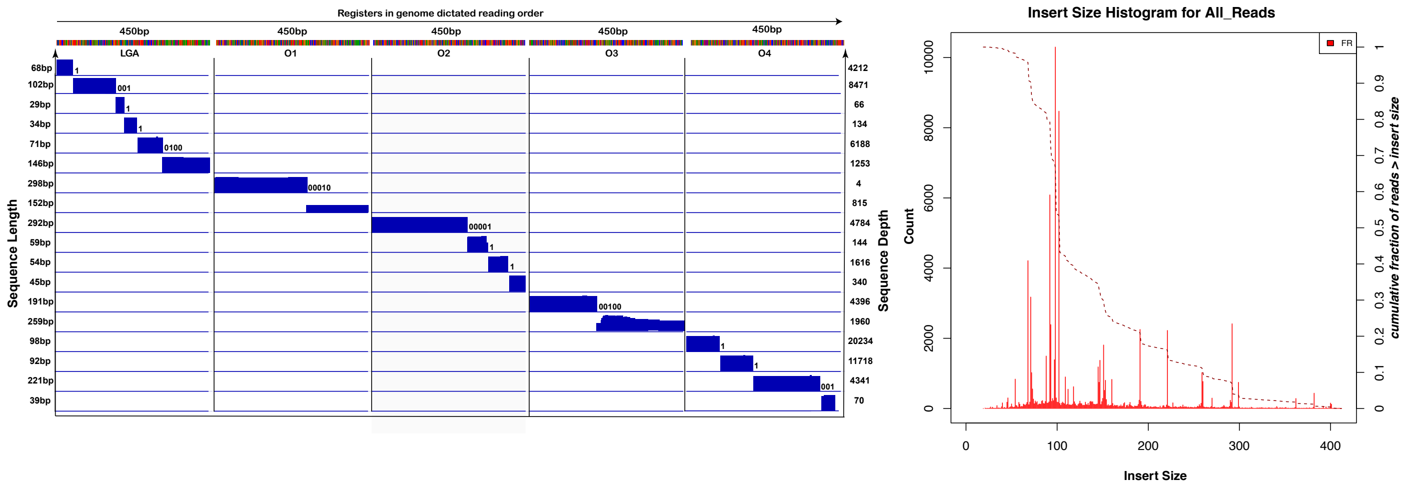


**d**

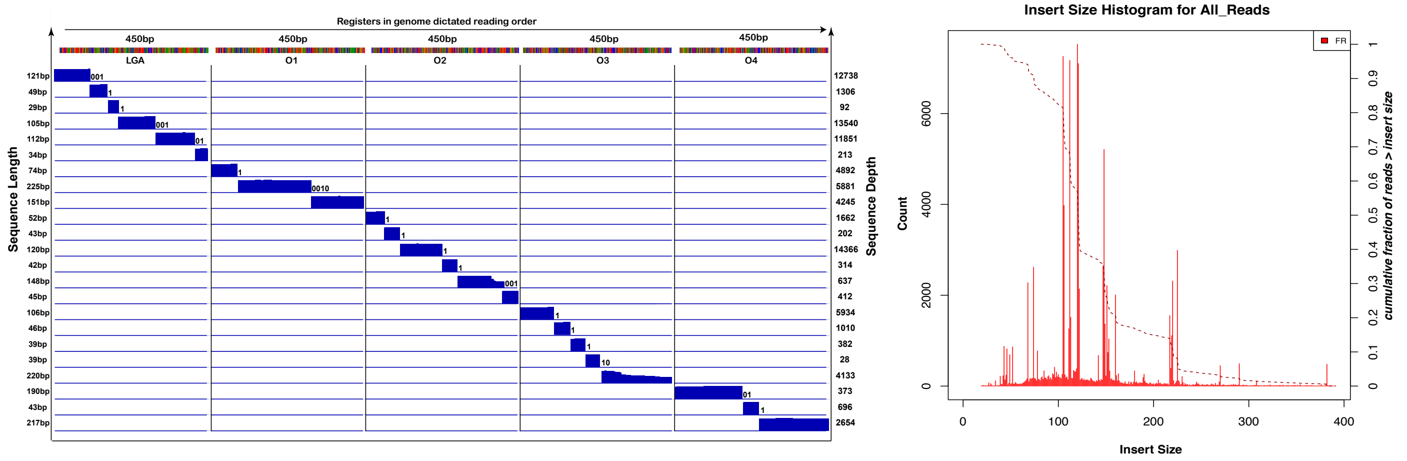


**Figure S10 | Multi-register data recording. a,b,c,** and **d**) A 126-bit string encoded in multiple orthogonal registers via nicking. All registers and corresponding gDNAs for the desired nicking sites were mixed in equimolar ratios. All reactions were performed simultaneously in 4 reaction tubes in parallel. Each vertical section represents one register, while the rows show the read lengths retrieved after sequencing analysis. The read lengths are recorded on the left axis and sequencing depths are indicated on the right axis.

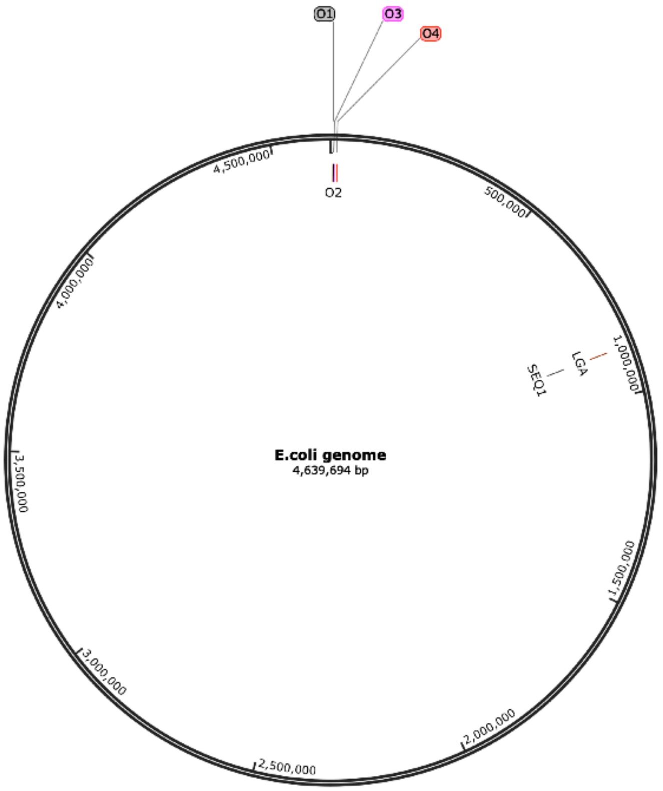


**Figure S11 | *E. coli* genome map, with all used sequences marked.** All of the registers; LGA, O1, O2, O3 and O4 are marked, along with SEQ1, which is used for toehold creation and *Pf*Ago vs. SpCas9 nickase comparative experiments. The map is generated via SnapGene Viewer.

**7. Solid state nanopore simulations**

To demonstrate the potential use of solid-state nanopores for nick detection, we performed all-atom MD simulations, where the nicked DNA (30 nts) is placed just above the $MoS_2$ nanopore membrane and solvated in a water box with $K^+$ and $Cl^-$ ions placed randomly to reach a neutrally charged system with a concentration of 1 M (Figure S12a). Two electrodes are placed in the *cis*- and *trans*-chambers of the ionic solution box, sustained by a constant voltage ($V_{TC}$) of 1 V to drive the nicked DNA through the nanopore. When the nicked DNA is pulled through the pore, it blocks the flow of ions that causes transient reductions in the ionic current measured by an external ammeter. Simultaneously, a lateral voltage ($V_{DS}$) is applied between the source and drain electrodes of the $MoS_2$ membrane to induce and record the transient variations of the transverse electronic sheet currents during the DNA translocation (a detailed description of these calculations may be found in the Methods section). The regions of negative "extremal" correlations between the ionic current and transverse sheet conductance strongly associate with the positions of the nicks.

Figure S12b displays the simulated ionic current (blue) and differential transverse sheet conductance (red) along with the center of mass (CoM) (green) of the nicked site on the DNA strand versus the translocation time of the entire DNA. Although the mean ionic current and transverse sheet conductance do not inherently indicate the presence of a single nick, an increase in the ionic current blockages to a global maximum and a consequent decrease in the sheet conductance to a local minimum during the small time frame, $t_n$, when the nicked site momentarily resides in the pore (represented by the blue box and yellow line from CoM curve at ~ z = 0 Å, where the nanopore membrane is positioned) is observed. The increase in the ionic current is attributed to the interaction between the nicked site with the $MoS_2$ membrane, when the DNA is pulled onto the $MoS_2$ membrane allowing for more ions to translocate through the pore. At the same time, the reduction in the sheet conductance is caused by a drop in the electrostatic potential contour of the nicked site due to the missing atoms on the backbone of the DNA compared to the normal DNA backbone. In conclusion, during the DNA translocation time window ($t_n$), there is only one event where the ionic current increases to a global maximum, and simultaneously the transverse sheet current decreases to a global minimum (within the time frame $t_n$). This event corresponds to the instance when the DNA nick is within the pore. Within other time windows, one may observe ionic current decreases and sheet current increases, but without the extremal magnitude values. For example, in Figure S13, the DNA translocates through the pore during the time interval 0 - ~6.5 ns, and only one nick-translocation event occurs around 2.5-

3 ns when the ionic (sheet) current reaches a global maximum (minimum. In every other time windows, for example during the time frame 3.5-4 ns or 5-5.5 ns, the sheet (ionic) current is above (below) other data points. This finding is due to conformational fluctuations, and not indicative of the presence of DNA-nicks. Similarly, on Figure S14, we can clearly see that the sheet current fluctuates significantly as there are many peaks and dips in the signal. But there is only one event occurring around 2-2.5 ns when the sheet current reaches a minimum and the ionic current increases to a maximum, indicating the presence of a nick.

As further support for this inverse correlation between the ionic current signal and the sheet conductance signal, we calculated the Pearson's correlation coefficient (P) between the two signals for every 0.5 ns frame during the translocation of the DNA molecule. Figure S12c shows the scatter plot of the normalized sheet conductance versus the normalized ionic current for the duration of the translocation time of the DNA strand. It can be clearly seen that there exists a strong inverse correlation (P = -0.8) only during the time interval of length $t_n$, when the nicked site is in the pore. A similar characterizing feature was found for the other two genomic regions used in the writing experiments (Figures S13 and S14), indicating the efficiency and reliability of $MoS_2$ nanopores in detecting single-nicks without denaturing the double-stranded DNA.

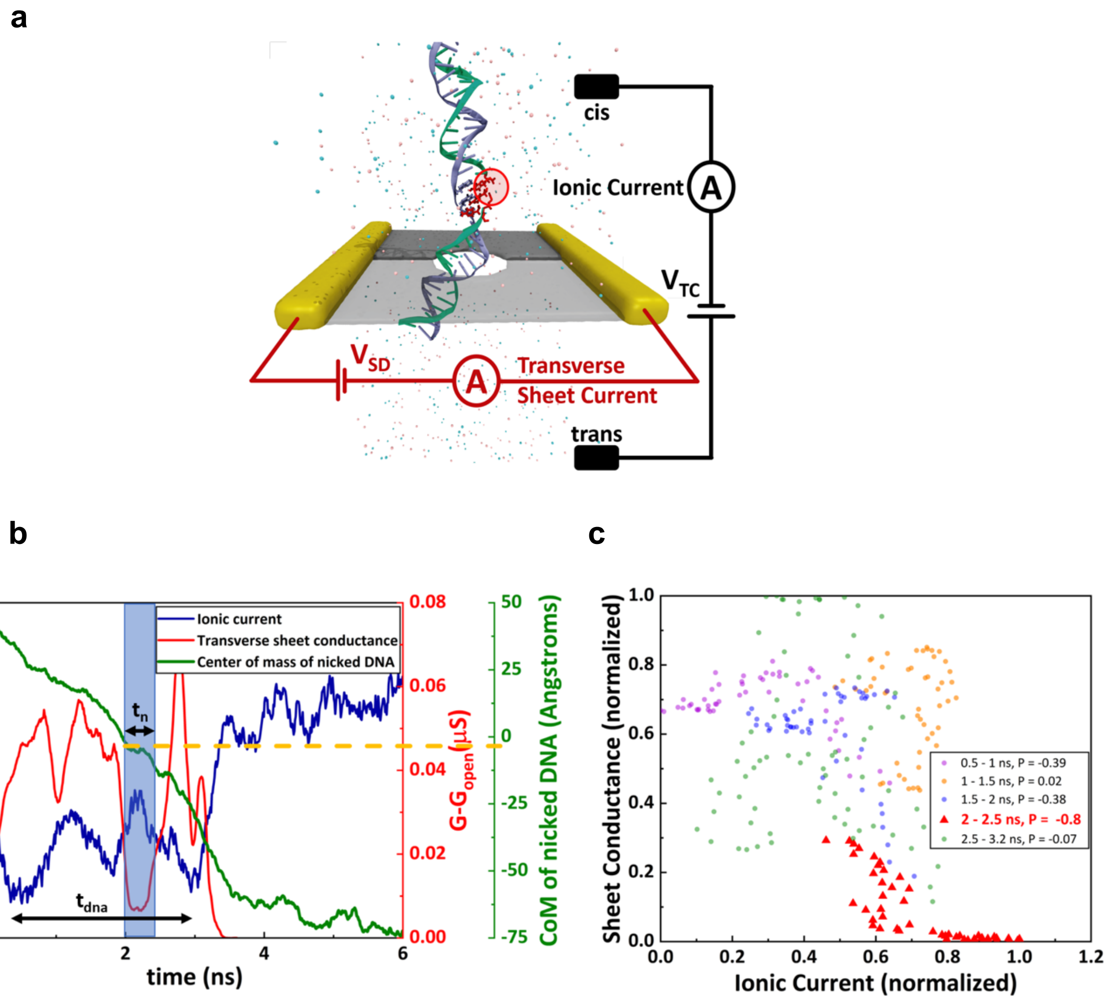


**Figure S12 | Reading with solid-state nanopores**.

(Sequence: 5'- AGACATCTGTTATCACTTAACCCATTACAA) **a**) Schematic illustration of the system setup for detecting the nick created at the center of a 30 nts dsDNA translocating through a MoS2 nanopore with a 2.6 nm diameter. **b**) Plot of the calculated ionic current (blue), differential transverse sheet conductance (red) and center of mass of the nicked site (green) versus time. $t_{dna}$ represents the translocation time of the entire dsDNA strand and $t_n$ represents the time frame during which the nick resides in the pore. **c**) The scatter plot of normalized sheet conductance versus normalized ionic current over time $t_{dna}$. There exist weak inverse correlations between the signals in the time frame 0.5 – 2 ns and weaker correlations within the time frame 2.5 – 3.2 ns (green). A strong inverse correlation with P = -0.8 is observed at 2 – 2.5 ns ($t_n$) (red).

**a** **b**

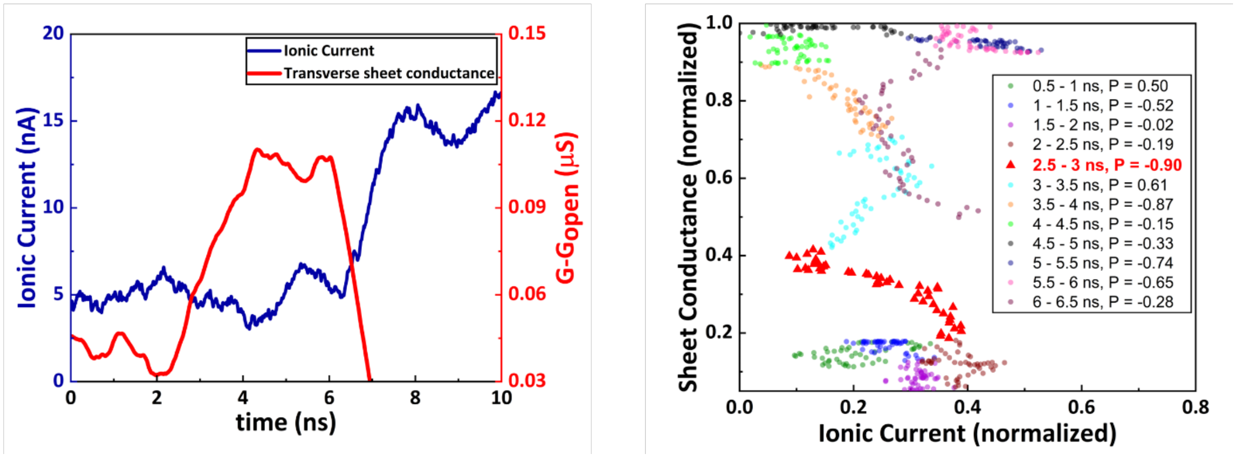


**Figure S13 | Reading with solid-state nanopores**
(Sequence: 5'–CAGATCAATGGAAGTAAATTC CAATTGTTT–3')**.** **a**) Plot of the calculated ionic current (blue) and differential transverse sheet conductance (red) versus time. **b**) Scatter plot of normalized sheet conductance versus normalized ionic current over translocation time. A strong inverse correlation with P = -0.9 is observed only during the time frame 2.5 – 3 ns (red) when the nick (between the bases -TA- in the given example) resides in the pore.

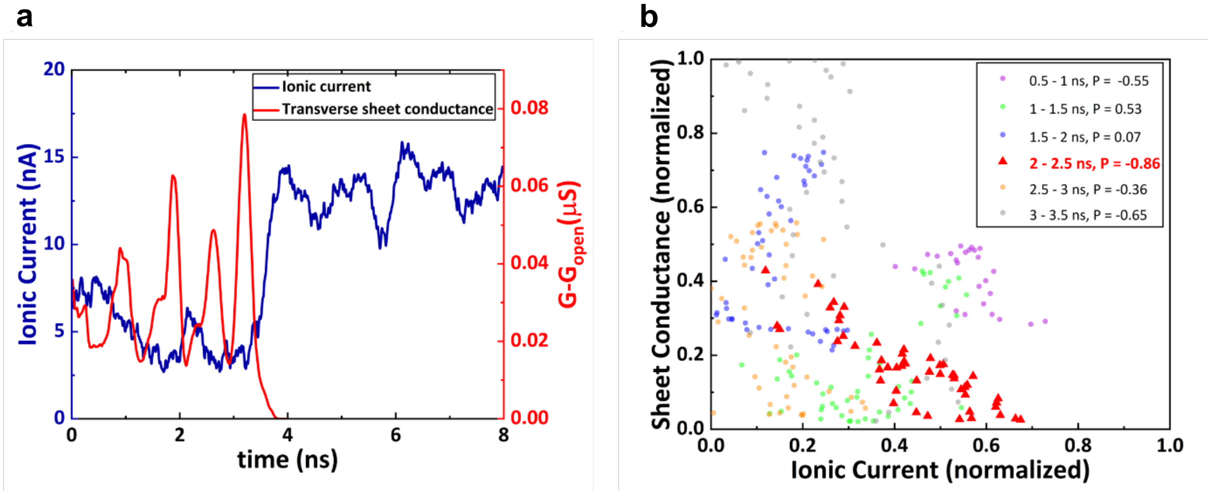


**FigureS14| Reading with solid-state nanopores**
(Sequence 5'-GAACGCCTCTGCAAGGTTCTTTG CTGCCGT-3'). **a**) Plot of the calculated ionic current (blue) and differential transverse sheet conductance (red) versus time. **b**) Scatter plot of the normalized sheet conductance versus normalized ionic current over translocation time. A strong inverse correlation with P = -0.86 is observed only during the time frame 2 – 2.5 ns (red) when the nick (between the bases -GG- in the given example) resides in the pore.

**8. Toehold creation**

As a proof of concept, we created a 14 nt toehold via *Pf*Ago on SEQ1. Both nicks (gray) are created simultaneously, in a single step, on the top strand, via two gDNAs (green). The reaction included: 15 min of heating at 70 °C, 10 min heating at 95 °C, gradual cooling down (0.1 °C/s) to 37 °C, and immediate cooling down to 4 °C. Immediate cooling from 37 °C to 4 °C is used to create the 14 nt ssDNA fragment between the two nicked sites (top strand: AGATCAATGGAAGT), and cause it to remain disassociated from the complementary strand so that the toehold (black) is created. The 14 nt disassociated sequence is then discarded in the process of purification via Zymo-Spin IC, silica-based columns (ZymoResearch®). The results of the MiSeq analysis prove that toehold can be created at the desired site (Figure S15).

**a**

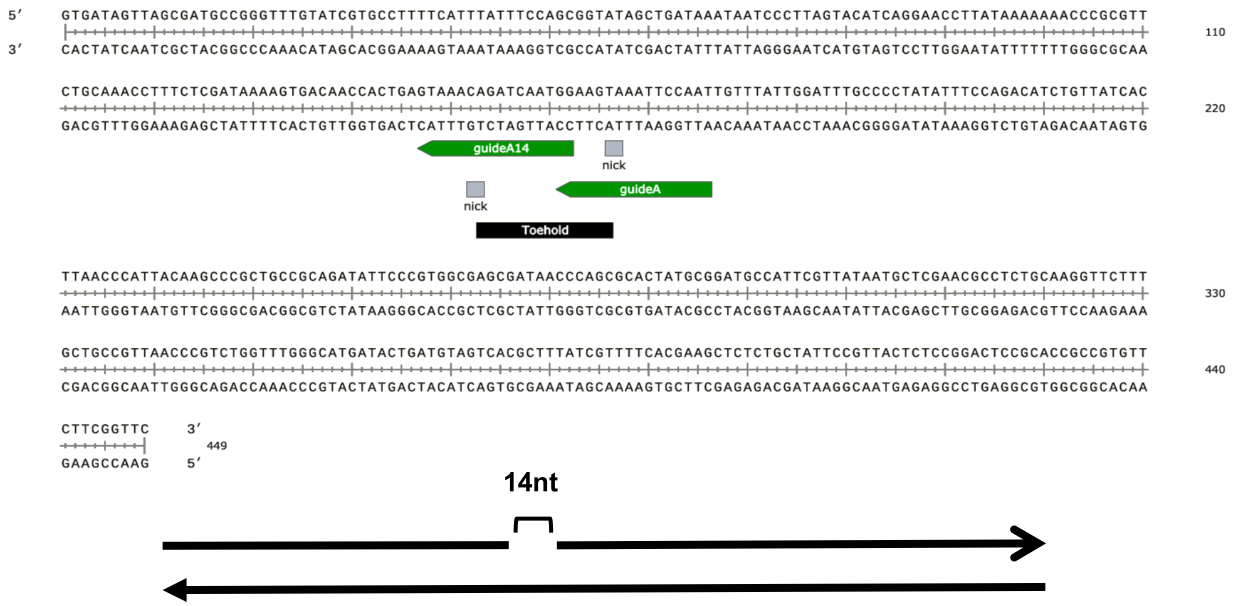


**b**

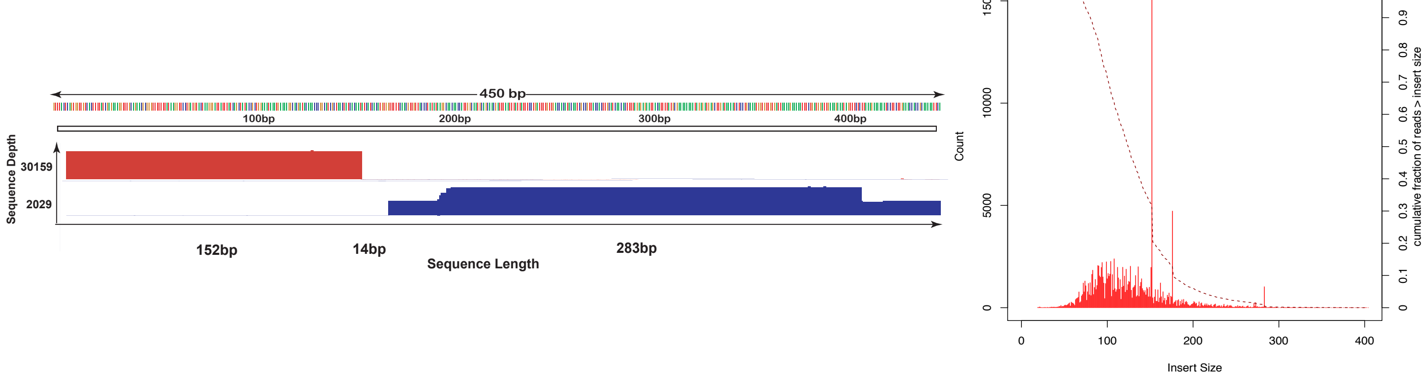


**Figure S15 | Proof of concept for toehold creation. a)** Map (top), sequence (middle) and final schematic view (bottom), describing a 2:1:2 pattern on the 449bp piece of DNA containing the toehold. Nicking sites are depicted in gray, gDNAs are depicted in green, while toeholds are depicted in black. Both nicks appear on the strand depicted on the top. **b)** MiSeq reads analysis shows that after dsDNA denaturation, two 152nt and 253nt ssDNA pieces are detected, confirming that a 14nt toehold was successfully created. The 14 nt gap in the coverage plot also clearly indicates the position of the created toehold.

**9. Non-destructive toehold detection**

We used the LGA register bearing the 10-bit string 0010000000 for the purpose of toehold creation (Figure S16). We used Guide 3, as well as a companion guide to nick a site within 14 nts of the original nick site (i.e., nicking site #3). Both nicks were placed on the sense strand and they lead to disassociation of the intermediary 14 nt long strand (ATTTTATCAAGCTC). It is straightforward to check if the toehold was created by detecting the disassociated strand.

The created toehold region may be used for bitwise random access and for determining the fraction of registers containing a bit at a given position (i.e., determine the concentration of the registers bearing a “1” at a given nicking site). For these experiments, the Reporter sequence for nondestructive reading of toeholds was purchased from IDT, and HPLC purified. The Reporter* was purchased non-purified from IDT. Fluorescence-based kinetics experiments were performed after mixing the native DNA registers and the Reporter. The reaction buffer used was a TAE/$Mg^{2+}$ buffer (0.04 M Tris, 1 mM EDTA, 12.5 mM $Mg^{2+}$, pH balanced to 8.0 by acetate). Outputs were measured on a BioTech Synergy H1 plate reader. The well plates of choice were low volume NBS (non-binding surface) 384 well plates with a clear flat bottom, purchased from Corning corporation (# 3544). The sample volume was chosen to be 18 µL, which resulted in a good signal-to-noise ratio, while minimizing the total DNA mass (i.e., experiment cost). Sealing tapes were attached to the surface of plates to avoid evaporation. The excitation wavelength was 485 nm and the emission wavelength was 520 nm. The excitation bandwidth was 9 nm and the emission bandwidth was 20 nm. Fluorescence was measured from the bottom to reduce the noise caused by the small droplets condensed on the sealing tape. Measurement started 2 min after samples were mixed. The background fluorescence signal (the Reporter by itself) was set to be the baseline and subtracted from all the other measured data.

Native PAGE was run to verify the products of non-destructive reading. 12% native PAGE gels of 0.75 mm thickness were made by mixing 6 mL 40% acrylamide/bis, 2 mL 10× TAE/$Mg^{2+}$ buffer and Milli-Q water to 20 mL, then adding 80 µL 10% APS and 10 µL TEMED to help polymerization. The native DNA register with the toehold and the Reporter were mixed and incubated at 25 °C for 1 hour before being loaded. The Reporter* was added after the register and the Reporter was incubated for 1 hour, while the mixture of these 3 species was incubated at 25 °C for another 2 hours before being loaded. The gels were run at 150 V and 25 °C for 4 hours. The 1×TAE/$Mg^{2+}$ running buffer was refreshed every 2 hours. Subsequently, the gel was directly scanned by a G:

Box Chemi XRQ imager without staining so that the self-fluorescence of each sample could be recorded. The excitation source was the UV illuminator and we used a standard filter.

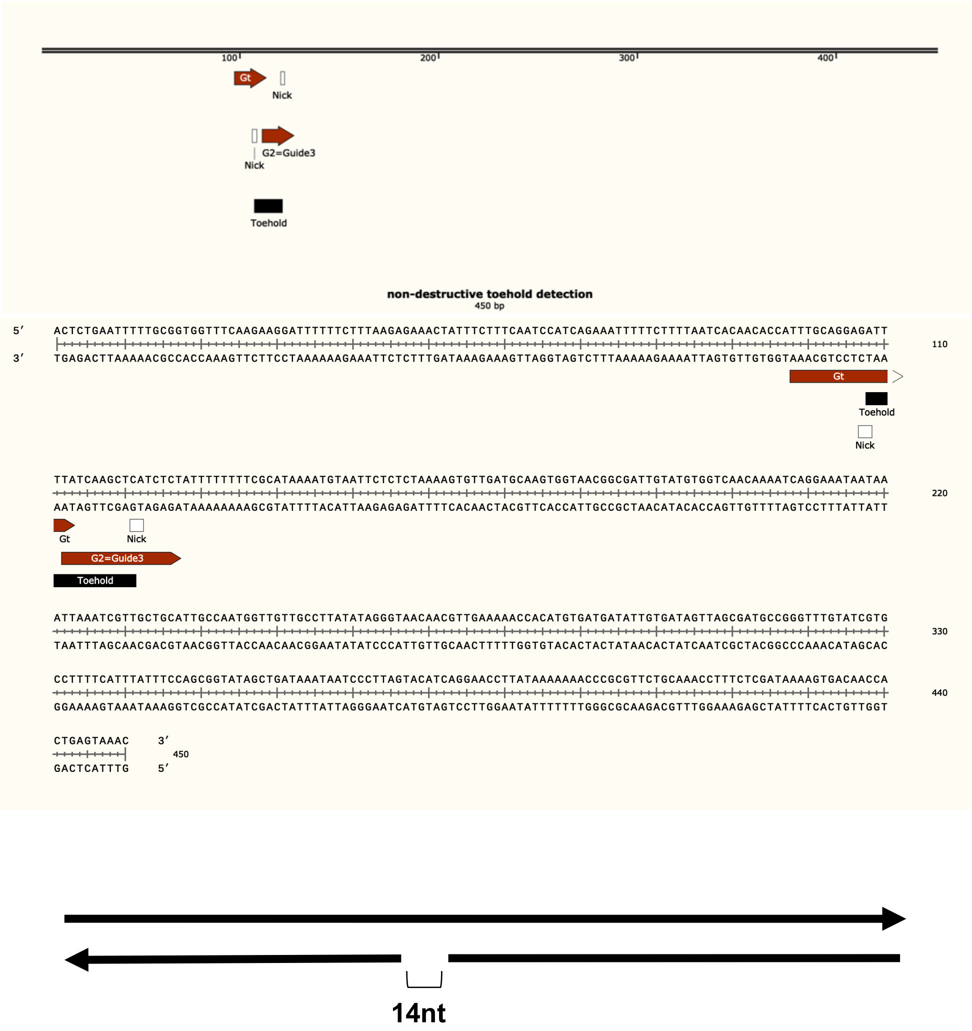


**Figure S16 |** Map (top) and schematic view (bottom) of the 450 bp LGA register sequence (LGA) used for non-destructive toehold detection. Red: The guides used for creating the two nicks. White: Nicks. Black: Toehold region.

#### 10. Extended discussion on Table 1

*Price per stored information bit*

There is no simple direct comparison regarding the costs of our methods and synthesis-based methods due to the fact that the different system components trade available DNA mass, storage density and system delays. Synthesizing a base pair costs ~\$0.001-\$0.01 (data obtained through quotation from Twist Biosciences and IDT) for an average mass of 0.2 fmols per oligo. Assuming that each nucleotide stores up to 1.57 information bits/bp due to coding and addressing redundancy (22), and ignoring the significant cost increase due to missing/low-copy oligos in the libraries (some vendors provide synthetic DNA with 30% of missing oligos, while others often have even higher losses that lead to <10% of usable oligos). Without the coding and missing oligo loss, one "synthetic DNA bit" costs between \$0.0005 to \$0.005. On the other hand, native DNA is virtually available for free; massive amounts of genomic DNA can be extracted and amplified for numerous rounds. The writing tool, *Pf*Ago, is also available at a very low cost: it can be extracted with a high yield that enables several thousands of reactions. The only synthesized components used in our system are gDNAs, which are essential for the writing part. Each 16 nt DNA oligo can be purchased from IDT at a cost of ~\$2.4/oligo, with a mass of 25-30 nmols. For 10 gDNAs, the overall cost is ~\$24. The DNA oligos are 5'-phosphorylated in house via polynucleotide 5'-hydroxyl-kinase (PNK) using the manufacturer's (NEB) protocol. These guides allow for creating a library of all 1024 possible nicking patterns on 10 nicking sites which suffices for storing up to $\frac{30nmol}{7.5pmol} * 1024 * 10\ bits =$ 40,960,000 bits (assuming that we use a 2:1 ratio of guide:enzyme). We hence estimate the cost of storing each bit replica by $\frac{\$24}{40{,}960{,}000\ bits}$ which is approximately $6 \times 10^{-7}$ \$/bits as reported in Table 1. Two remarks are in place. First, the reliability and efficiency of the nicking process is very high (~90%), which is a positive feature of our system as there are no losses associated with missing oligos and synthesis errors. Second, part of our cost reduction may also be attributed to the special organization of the DNA, and hence a more conservative cost reduction estimate is two orders of magnitude.

*Writing/reading latency*

In synthesis-based methods, writing is performed via iterative DNA nucleotide integration which takes hours to complete. In our proposed strategy, the writing step basically involves a set of nicking reaction which can be performed for hundreds of registers simultaneously and do not take more that 40 min (see the Methods section).

On the other hand, the current reading protocol includes library preparation, MiSeq sequencing and reference alignment, which all take time on the scale of hours. This matches the readout times reported for all other synthetic DNA-based storage methods. Nevertheless, as already mentioned, toeholded native DNA may be read very accurately using solid state nanopores (6), while simulation evidence suggests that the same is true of nicked DNA.

*Random access and strand displacement*

The proposed method for creating non-destructive toeholds enables reading the stored information content in a bitwise fashion without ever sequencing the register content. None of the currently known molecular storage platforms has this important feature. In addition, being able to create toeholds allows for performing various toehold-mediated DNA strand displacement reactions, and potential in-memory computing operations. This is another feature of the system not matched by any other existing molecular storage platform.

*Information density*

The ultimate storage density of synthetic DNA-based methods is 2bits/bp, which cannot be achieved by current methods due to the *need to protect against both synthesis and sequencing errors.* For portable DNA-based data storage platforms with nanopore readouts and addressability features, the highest information density so far has been reported to be 1.575 bits/bp (5). In our system, nicking positions are kept at least 25 nt apart, in order to prevent disassociation of the inter-nick strand, which decreases the information density as follows. In our experiments, we recorded $10\log_2 3$ bits on a register of length 450 bps, which results in a density of $\frac{10\log_2 3\ \text{bits}}{450\text{bp}}$ = 0.036 bit/bp. Nevertheless, this information density may be easily increased by raising the number of nicking sites per register (i.e., longer registers or more densely packed nicking sites). For example, by placing nicking sites exactly 25 bps apart, we can increase the number of nicking sites to 18 and the density to 0.0648 bits/bp, while placing nicks exactly 20 bps apart results in a density of 0.08 bits/bp. The recommended approach for increase well density is multi-register writing, as discussed in the main text and supplementary information.

*Coding loss*

Another important feature of punch card DNA recorders is that they exhibit very high recording and readout accuracy and hence do not require large (if any) coding overheads for recording. In synthetic DNA-based storage system, errors occur during the synthesis process either in the form of substitutions, at an average rate <1/200 nt (IDT) or coverage errors, which manifest themselves as missing oligos (Twist, Agilent). In the former case, we observed worst-case error rates of 0.023/nt, or 4 substitution errors per oligo length 170 nts. To correct 4 errors, one needs a minimum distance of at least 9, and for oligos of length 200 bps, the maximum dimension of a 4-error correcting code over a 4-letter alphabet is 176. Hence, the coding redundancy for IDT-based synthesis is roughly 12%. It is significantly more difficult to combat a combination of in-oligo substitution and coverage errors. For the latter case, the interested reader is referred to a previous work (8), proposing the use of Reed-Solomon codes. To combat sequencing errors, one can use large sequencing coverage coupled with additional substitution error redundancy (roughly 10%). In this setting, the coding redundancy amounts to roughly 20% of the sequence length, and a density of 1.6 bits/nt. Further reductions are needed to combat deletion and insertion errors encountered during nanopore sequencing (7).

*Maximum achievable physical density*

It is challenging to provide an accurate and fair comparison of all known DNA-based storage systems, due to the many different performance metrics adopted for such comparisons. A number of prior works used physical density (9,10) to assess the storage capacity of the system, without taking into account the cost and latency of the various components. In (9), an ultimate physical density of 680 PB/g was estimated under idealized assumptions pertaining to error-free synthesis, while in (10) it was shown that under perfect indel correction, an actual density of 215 PB/g and an ultimate physical density of 200 EB/g (considering 1 copy of each information bearing oligo) is achievable. Our scheme **does not rely on any idealized assumptions** regarding synthesis errors as the nicking process is extremely accurate and does not require error-correction redundancy. It nevertheless still offers an achievable density of ~4 EB/g. The density loss is ~50-90 fold compared to (10), which at this level of data compaction is barely relevant, while the savings in cost is roughly four orders of magnitude, and the encoding process highly efficient due to parallel nicking. And, as pointed out in the main text and reiterated in the Supplement, nick-based storage also allows for in-memory strand/nick-displacement computing, bitwise random access, pooling and many other features not matched by any other technology.

*Comparison with alternative native DNA-based storage methods*

Our native DNA-based punch card system allows for cheap and low-latency molecular recording with built-in computational capabilities. Storing a bit replica (one physical copy of the bit) of information costs less than $\$0.0000006$ (Extended Table 1), and the library creation and placement process take less than 1 hour (Table 1). Furthermore, given that *E. coli* and *Pf*Ago enzymes are readily available, the mass of created DNA products can by far exceed that of synthetic DNA, therefore enabling orders of magnitude more readout cycles with NGS devices. Another important feature of nicking-based writing is that one *Pf*Ago molecule with one single guide may be used to make multiple nicks, as it is a multiple turnover enzyme. Hence, a single guide string that carries 32 bits of information can be used to nick more than hundred bits of information. The system also allows for using a ternary message alphabet, based on selecting nicking positions at both the sense and antisense strand (Figure S8). Furthermore, as only positions bearing '1's (or '1's and '2's) need to be stored, the number of nicking steps is effectively reduced by one half. In addition, unlike synthetic DNA, native DNA registers may be of lengths 10,000 bps or longer, and if properly chosen to have little sequence similarity, multiple registers may be nicked in the same vial without any cross-interactions to further speed up the recording process. If multiple registers from the same genome are used, the order in which the bits are read is induced by the natural order of the registers in the bacterial genome. Finally, the nicking reactions can take place simultaneously so that the writing latency is significantly reduced compared to that of synthetic methods.

As a final remark, we observe that further cost savings are possible through the use of registers that share multiple substrings of lengths 16 with high sequence similarity. This is due to the fact that the *Pf*Ago/nicking guide complex may create nicks even when there is a <15-20% mismatch between the guide sequence and the native substring to be nicked. Hence, the same guide may be reused to nick sites at different registers.

#### 11. Combinatorial register mixing

The proposed architecture may be improved in terms of DNA spot density using combinatorial mixing of strands. The simplest means of mixing is to mix the guide sequences corresponding to native DNA substrings directly, placing them in positionally-encoded wells rather than using them for nicking. For the running *E. coli* example one could choose more than 100,000 starting positions for the guides and encode a 100,000-bit long message through the presence or absence of a guide in a well dictated by the binary content (presence - ‘1’, absence – ‘0’). The order of the guides within the reference genome naturally dictates the ordering of the binary symbols within the well. Although conceptually simple, this approach has multiple drawbacks. It requires a pool of hundreds of guides of length 16 to represent a single bit, while one single guide molecule may be used to record hundreds of bits due to the high turnover rate of *Pf*Ago (here, one can think of these hundred bits being replicas of the same bit, leading to a 100-fold increase in the volume of the nicked DNA; alternatively, the same guide/enzyme complex may be used to nick positions in different registers that contain substrings at small edit distance from the reverse-complement of the guide sequence. In this setting, the multiple turnover enables storing hundreds of different bits). Using pools of guides for information storage also limits the number of readout cycles available with the given scheme, and it prevents the use of nanopore sequencers due to the short length of the oligos. Furthermore, with such an implementation, strand-displacement computations are impossible.

On the other hand, nicked registers may be combined in a combinatorial manner, by mixing both different registers and the same registers with different nicking patterns. If nanopores are used for sequencing, individual register information may be retrieved, and the mixture composition determined based on the individual reads. For our running example involving a register with 10 nicking positions, there are $2^{10}$=1024 possible binary nicking patterns. Any mixture of k-out of-1024 patterns, where k=0, …,1024, may be used to encode information. In total, the number of distinguishable mixtures equals $2^{1024}$, resulting in 1024 bits of information. Hence, mixing provides an exponential increase in information content per spot, but requires sequencing of individual registers, which is only possible when using nanopore readouts.

If high-throughput Illumina sequencers are used instead, the identity of the individual registers will be lost, and one would only have access to in-between nicks fragments of the nicked registers.

In this case, one has to resort to using specialized mixing schemes designed for *group testing* and *adder channels*, well-studied entities in information theory. Group testing effectively allows for mixing of k-out-of-N registers, where k<<N. In this case, the number of nicks on the group-testing registers scales as $k \log_2 N$. The optimal mixing scheme for adder channels is the Lindström scheme (11), and for simplicity, we only describe this scheme and the corresponding experiments.

The Lindström scheme is a group testing (coding) method that allows one to identify individual components in a real-valued sum of binary weighted vectors. The scheme is asymptotically optimal in so far that it can discriminate between any of the $2^N$ distinct sums of N binary vectors of length as small as

$$M = \frac{2N}{\log_2 N},$$

as $N \to \infty$. This result has the following interpretation: for any collection of $k \in \{0,1,2,\ldots,N\}$ out of N binary vectors of length M in the Lindström code, $x_{i1}, x_{i2}, \ldots, x_{ik}$, the sum of the vectors

$$x_{i1} + x_{i2} + \cdots + x_{ik} = s$$

uniquely determines the identity of the individual component vectors $x_{i1}, x_{i2}, \ldots, x_{ik}$.
As an illustration, consider the following collection of four vectors of length three:

| | | | |
|---|---|---|---|
| 1 | 0 | 1 | 1 |
| 0 | 1 | 1 | 0 |
| 1 | 1 | 0 | 0 |

Let the collection of the vectors in the sum be encoded by the binary vector $(a,b,c,d)$. For example, $(a,b,c,d) = (1,1,0,0)$ means that the sum involves the first two vectors, $(101)^T$ and $(011)^T$ and hence equals $(112)^T$. For an unknown collection $(a,b,c,d)$ we have the following equations:

$$a + c + d = s_1,$$
$$b + c = s_2,$$

$$a + b = s_3.$$

Clearly, $s_1 + s_2 - s_3 = 2c + d$. Consequently, since $a, b, c, d$ are all binary variables, the value of $2c + d$ uniquely determines both $c$ and $d$. Once these two values are known, the sum

$$s_1 + s_3 = 2a + b + c + d$$

may be used to determine the value of $2a + b$ and hence the individual values of the bits $a, b$. In addition to this simple decoding scheme, one can easily see that all $2^4$=16 different sums of the column vectors are distinct. This distinguishability of sums property will be used to determine the mixture composition of registers without resolving to the use of coverage information, but rather only referring to fragment overlaps and suffix identities, as described on the example to follow. For a general description of the scheme, the reader is referred to (11).

Note that the above scheme was generalized to allow for using non-binary weights in the sum (e.g., in the example above, the elements in $(a, b, c, d)$ may take nonbinary values) in (12). Non-binary values in this setting amount to using different concentrations of the corresponding registers. For example, if four different concentrations are used, $(a, b, c, d) = (2,1,0,3)$ would imply using 1x concentration of the nicked register corresponding to $b$, 2x of the concentration of the nicked register corresponding to $a$ and 3x of the concentration corresponding to $d$. Given the large mass of readily available native and nicked native DNA, this concentration-based code alphabet enlargement is easily implementable. Nevertheless, this approach is out of scope of the proposed experimental investigation.

In the context of nick-based storage, the above results suggest that one can *mix* the same registers with different nicking patterns and with possibly different concentration and still determine, based on the fragments present, what the actual nicking patterns of the individual registers were. For a q-ary concentration alphabet, $\mathrm{N} \log_2 q$ bits can be stored using registers with $\frac{2N}{\log_2 N}$ nicking positions. As on average only half of the positions need to be nicked for recording, $\frac{N}{\log_2 N}$ nicks can be used to store $\mathrm{N} \log_2 q$ bits.

Mixing registers increases the information content per spot but preserves random access to $\frac{2N}{\log_2 N}$ - bit substrings rather than single bits. Computations in memory based on nick-displacement can be easily adapted to this setting as single instructions may be used to update the same register sequences bearing different recorded nick patterns (13,14).

For a practical illustration of the Lindström scheme, we used the following collection of N=17 binary vectors of length M=10, listed column-wise:

| | **1** | **2** | **3** | **4** | **5** | **6** | **7** | **8** | **9** | **10** | **11** | **12** | **13** | **14** | **15** | **16** | **17** |
|---|---|---|---|---|---|---|---|---|---|---|---|---|---|---|---|---|---|
| **1** | 1 | 0 | 0 | 0 | 1 | 1 | 1 | 1 | 1 | 1 | 0 | 0 | 0 | 0 | 1 | 1 | 1 |
| **2** | 0 | 1 | 0 | 0 | 1 | 0 | 0 | 0 | 0 | 0 | 1 | 1 | 1 | 1 | 1 | 1 | 0 |
| **3** | 0 | 0 | 1 | 0 | 0 | 0 | 1 | 0 | 0 | 0 | 1 | 0 | 0 | 0 | 1 | 0 | 0 |
| **4** | 0 | 0 | 0 | 1 | 0 | 0 | 0 | 0 | 1 | 0 | 0 | 0 | 1 | 0 | 0 | 0 | 0 |
| **5** | 1 | 1 | 0 | 0 | 0 | 0 | 1 | 1 | 1 | 1 | 1 | 1 | 1 | 1 | 0 | 0 | 0 |
| **6** | 1 | 0 | 1 | 0 | 1 | 1 | 0 | 0 | 1 | 1 | 1 | 0 | 0 | 0 | 0 | 0 | 0 |
| **7** | 1 | 0 | 0 | 1 | 1 | 1 | 1 | 1 | 0 | 0 | 0 | 0 | 1 | 0 | 1 | 1 | 1 |
| **8** | 0 | 1 | 1 | 0 | 1 | 0 | 1 | 0 | 0 | 0 | 0 | 0 | 1 | 1 | 0 | 0 | 0 |
| **9** | 0 | 1 | 0 | 1 | 1 | 0 | 0 | 0 | 1 | 0 | 1 | 1 | 0 | 0 | 1 | 1 | 0 |
| **10** | 1 | 1 | 1 | 0 | 0 | 0 | 0 | 0 | 1 | 1 | 0 | 0 | 1 | 1 | 1 | 0 | 0 |

To illustrate the protocol for mixing the same register (LGA) with different nicking patterns, we tested three mixtures, including the first three columns (1+2+3), and two subsets of size two of the first three columns (1+2 and 1+3), all in equimolar ratios. We also sequenced the individual columns (1, 2, 3) as controls that allow us to determine if the nicking reactions were successful. The coverage plots and an illustration of the scheme are shown in Figure 17a-c.

Based on the fragments observed in the mixture, since the sequence and the position of each fragment is predetermined, the following observations are immediate:

1- Two fragments that do not overlap: these may be part of the same or different nicking patterns; which of these two cases is true is determined by observing gaps between previously placed fragments;
2- Two fragments that overlap: these have to belong to two different nicking patterns.

Using this rule, one can assign each fragment to its appropriate position and place it in the appropriate nicking context. The procedure resembles simple puzzle solving with rectangular blocks indicating fragments (as illustrated in Figure17b). However, one may observe that in all the above cases, there exist two blocks that end at the same location in the reference genome (Figure17c). In this case, at first it appears that the nicked fragment reconstruction procedure may not produce unique results. But this is not the case, as all combinations of columns generated by the Lindström code have distinct sums and therefore, only one of the alternative placements is allowed by the code.

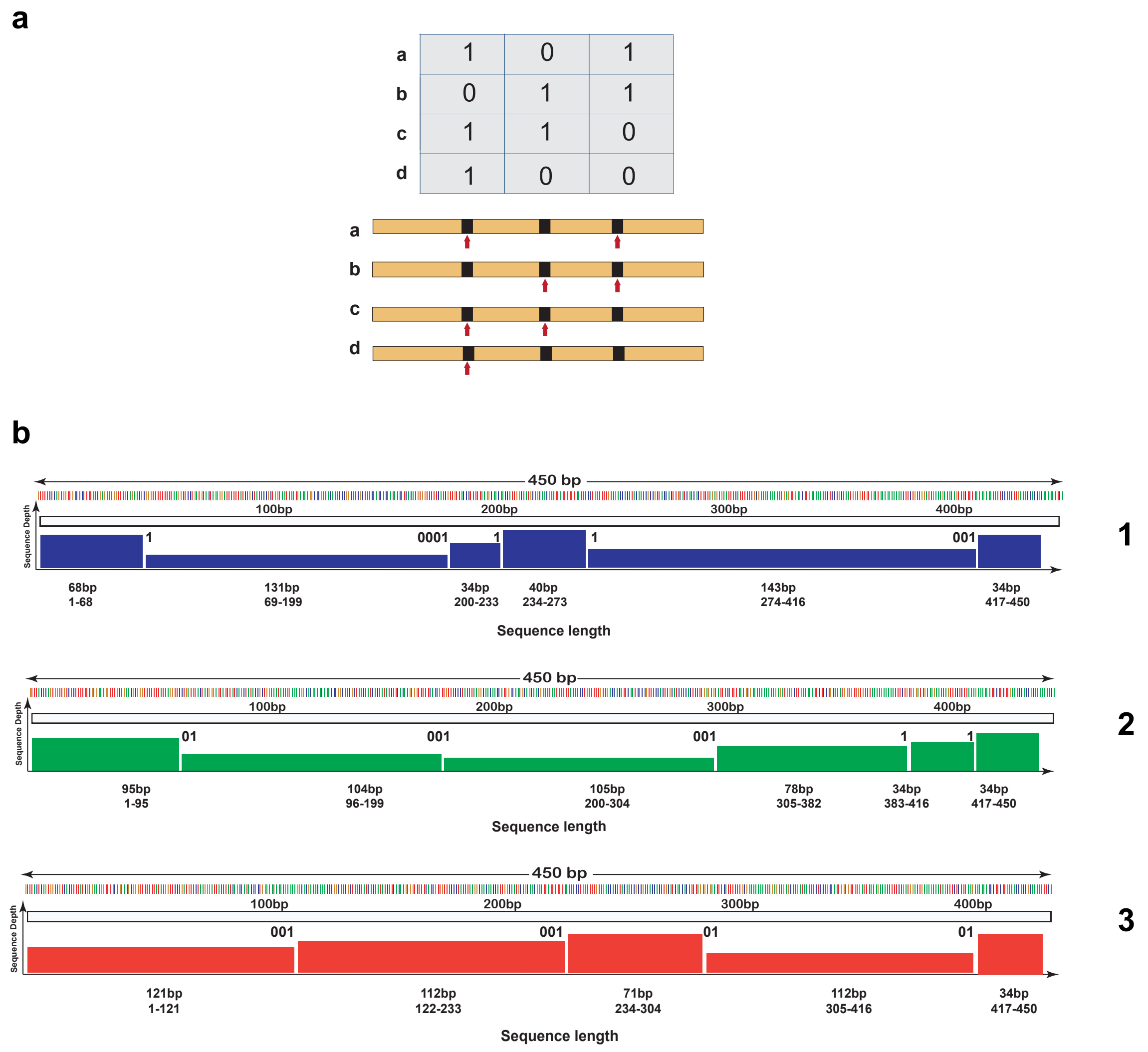

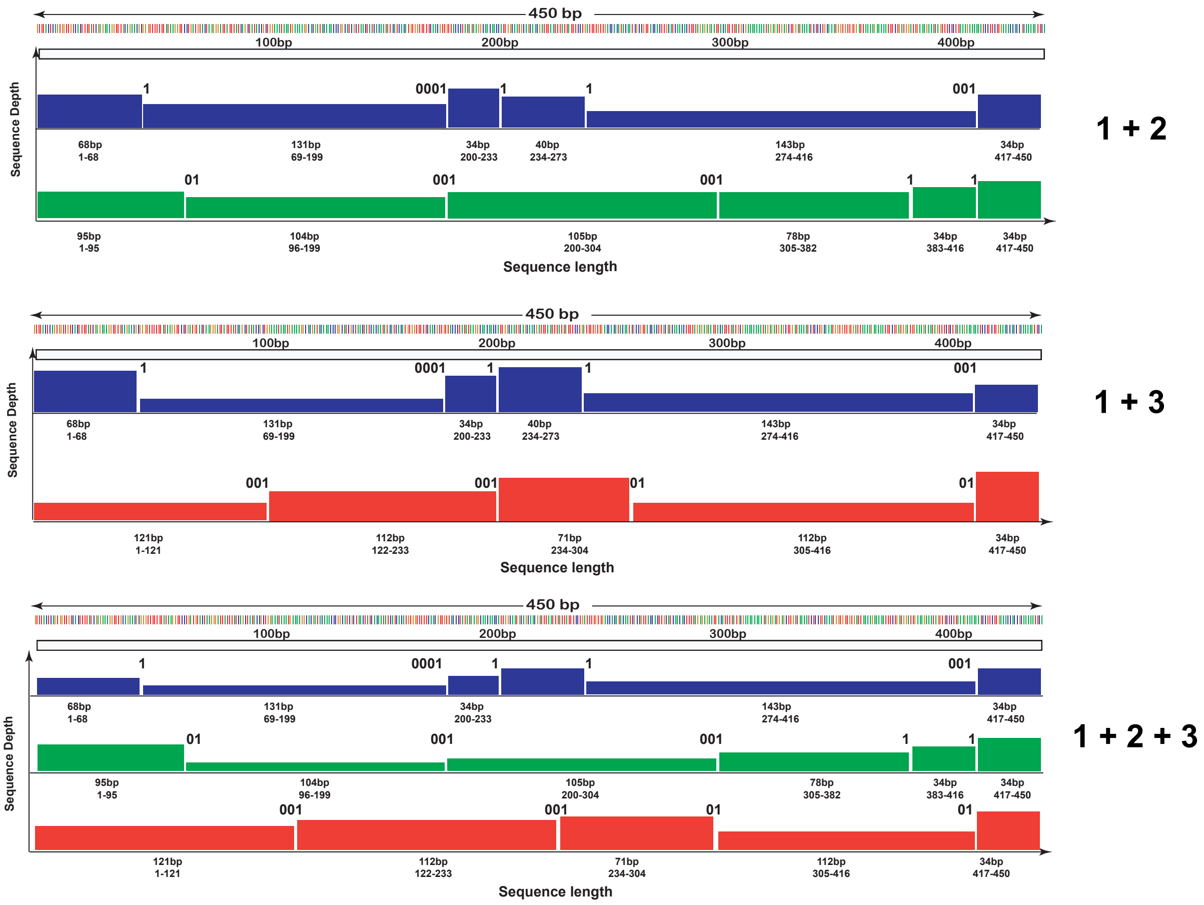

450 bp
100bp
200bp
300bp
400bp
Sequence Depth
Sequence length
68bp 1-68
131bp 69-199
34bp 200-233
40bp 234-273
143bp 274-416
34bp 417-450
95bp 1-95
104bp 96-199
105bp 200-304
78bp 305-382
34bp 383-416
121bp 1-121
112bp 122-233
71bp 234-304
112bp 305-416
1 + 2
1 + 3
1 + 2 + 3


c

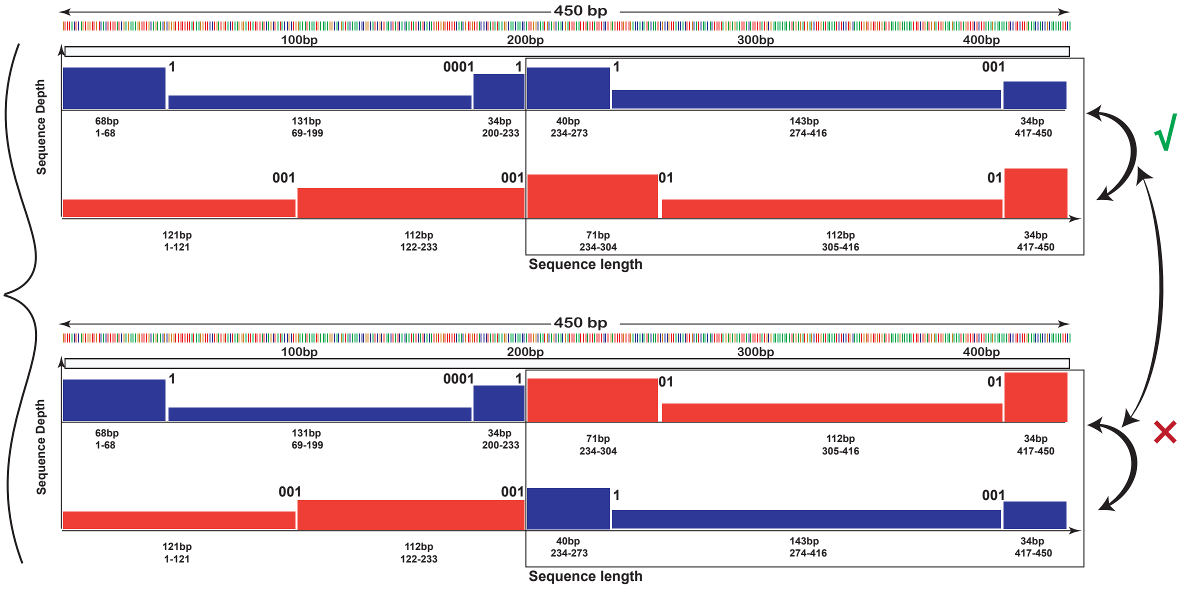

450 bp
100bp
200bp
300bp
400bp
Sequence Depth
Sequence length
68bp 1-68
131bp 69-199
34bp 200-233
40bp 234-273
143bp 274-416
34bp 417-450
121bp 1-121
112bp 122-233
71bp 234-304
112bp 305-416
√
×

**Figure S17 | a)** A collection of four vectors of length three, illustrating how to arrange fragments generated by the Lindström scheme for combinatorial mixing of replicas of the same register bearing different nicking patterns**. b)** The coverage plots of the individual patterns (1, 2 and 3), along with the mixed patterns (1+2, 1+3 and 1+2+3) in their resolved formats. The coverages are shown qualitatively on the y axes, while the sequence lengths are shown on the x axes. **c)** As an example, an alternative pattern (bottom) may be obtained for mixture 1+3 (as well as other mixtures), compared to the correct pattern (top). Consequently, one may read the binary vectors as {1000110101,0010011001} instead of {1000111001, 0010010101}. Nevertheless, a quick inspection shows that the pair of vectors 1000110101,0010011001 is not present in the Lindström code.

#### C. Additional Supplementary Tables

##### Table S4. List of DNA sequences used in this work.

| | |
|---|---|
| **Primary 1kbp sequence for nicking activity verification** | 5'–GCGAACAGCAGCGGTTATGGACAATATTGAACGGTTAATCTCCGCACAGCGGTTGCAGG CTATTTCTGGCGTCGGAGCCTTTAGTTTCAATACGTTATATAAGTTGGTGTGGTTGAAAGAA AATCATCCACAACTGCTGGAACGCGCGCACGCCTGGCTCTTTATTTCGTCGCTGATTAACC ACCGTTTAACCGGCGAATTCACTACTGATATCACGATGGCCGGAACCAGCCAGATGCTGGA TATCCAGCAACGCGATTTCAGTCCGCAAATTTTACAAGCCACCGGTATTCCACGCCGACTCT TCCCTCGTCTGGTGGAAGCGGGTGAACAGATTGGTACGCTACAGAACAGCGCCGCAGCAA TGCTCGGCTTACCCGTTGGCATACCGGTGATTTCCGCAGGTCACGATACCCAGTTCGCCCT TTTTGGCGCTGGTGCTGAACAAAATGAACCCGTGCTCTCTTCCGGTACATGGGAAATTTTAA TGGTTCGCAGCGCCCAGGTTGATACTTCGCTGTTAAGTCAGTACGCCGGTTCCACCTGCGA ACTGGATAGCCAGGCAGGGTTGTATAACCCAGGTATGCAATGGCTGGCATCCGGCGTGCT GGAATGGGTGAGAAAACTGTTCTGGACGGCTGAAACACCCTGGCAAATGTTGATTGAAGAA GCTCGTCTGATCGCGCCTGGCGCGGATGGCGTAAAAATGCAGTGTGATTTATTGTCGTGTC AGAACGCTGGCTGGCAAGGAGTGACGCTTAATACCACGCGGGGGCATTTCTATCGCGCGG CGCTGGAAGGGTTAACTGCGCAATTACAGCGCAATCTACAGATGCTGGAAAAAATCGGGCA CTTTAAGGCCTCTGAATTATTGTTAGTCGGTGGAGGAAGTCGCAACACATTGTGGAATCAGA TTAAAGCCAATATGCTTGATATTCCGGTAAAAGTTCTCGACGACGCCGAAACGACCGTCGC AGGAGCTGCGCTGTTCGGTTGGTATGGCGT–3' |
| **Primary 1.2kbp sequence for nicking activity verification** | 5'–AGCGAATGCAACGGGTGATGAAGTCGTATCATTAGATAAACACATCAACACCAGTGCTA CTGATACAGATCAAATACAAGCATTTATAGTTTCAACATGGATGGCGCCTTTTCAAAATGATA TGTATTCAGAAGATAATCCTATCTCACCTTACTATAAAATTGAGTGGTAGACGATTGACCAAA TAAATCATAACGAGATACTTTTATAATAGTATCTCGTTCCTTTCATTCAATCCTCACACATGAA AAAATATGCGCTCACCAAAAGTTAAATTCTTAACCATCTTCACGTTTTGTATATTTATTACAAA AATGTCTTTCGCCAGCAACTCTTGCTCTAATGAAGCAGGAACAATGTTCAGAATCGAACCAA ATTTAATTAAAGCTATCGCTCTTGTTGAAAGCAATCTAAAAAAGGATAGCATAGGCAAGAAC AGAGATAAGAATAACAATATAAAGAGTTTAGATTATTGGTTAATGCAGATAAACCAGATGCAT ATTCCCCTGTTAAAAAAACGTGGAATAATAAAAGATGAACGAGATTTGTTAGATAACCCTTGT CTGAACATTAAAATAGGAACCGAGATTTTATATAACCATTTTTCACGTTGTGGAGTAACGTG GCAATGCCTGGGGACTTATAATGCGGGTTTTGCAATGGATAACCAAAAAAAGAGACAGCAA TATGCCCCAAAATATATATTGTATATACCAGGCTTAATGAACTAGACAATCGTAAGGCTTCAG CAAAATGAGCCTAAAGCCTCTTTTTTTTATATAGTGCAAACACCCATACGTAAAGCTGTTTTG ATCAACTCGATACCACTGTGAACATCTAATTTCTTCATGATATTGAGTCTATGAGTTTCGACT GTTTTCCTGCTTAACTGCAAGAAATTAGCGATTTCTTTGTTTGTTTTACCATCAGCTATGAGT TGTAATATTTCTGACTCACGATTTGTAAGTTGATTATCAGAGGAATACCTGGAAGATATTTTT TCACAATCCATATGTACACTATCAAAATAAGTGTAACCATTACTAATCGAATCAATAGCACGA TTTAGTTTCGCCGGGTGACTGGTTTTCCATACGCATCCTTTCGCACCTGCTTCAAGCAATTT TATTGAATCAATATACGGCATCTTTGCTACAAATGTAATAATATGAGCATCTATCTTATGCGA TTTAACCCATTTTACGAGTTCGGCCCCCATCATTCCACCTGCA–3' |
| **Primary 1.5kbp sequence for nicking activity verification** | 5'–CGTTGCTGCATTGCCAATGGTTGTTGCCTTATATAGGGTAACAACGTTGAAAAACCACAT GTGATGATATTGTGATAGTTAGCGATGCCGGGTTTGTATCGTGCCTTTTCATTTATTTCCAG CGGTATAGCTGATAAATAATCCCTTAGTACATCAGGAACCTTATAAAAAAACCCGCGTTCTG CAAACCTTTCTCGATAAAAGTGACAACCACTGAGTAAACAGATCAATGGAAGTAAATTCCAA TTGTTTATTGGATTTGCCCCTATATTTCCAGACATCTGTTATCACTTAACCCATTACAAGCCC GCTGCCGCAGATATTCCCGTGGCGAGCGATAACCCAGCGCACTATGCGGATGCCATTCGT TATAATGCTCGAACGCCTCTGCAAGGTTCTTTGCTGCCGTTAACCCGTCTGGTTTGGGCAT GATACTGATGTAGTCACGCTTTATCGTTTTCACGAAGCTCTCTGCTATTCCGTTACTCTCCG GACTCCGCACCGCCGTGTTCTTCGGTTCAAGTCCCAACATCCGGGCGAACTGGCGTGTTTC ATTAGCCCGGTAGCATGAACCATTATCCGTCAGCCACTCCACTGGAGACGACGGAAGATCG TTGCCGAAGCGGCGTTCCACCGCTCCCAGCATGACGTCCTGTACTGTTTCACTGTTGAAGC CGCCGGTAGTGACCGCCCAGTGCAGTGCCTCACGATCACAGCAGTCCAGCGCGAACGTGA CACGCAGTCTCTCTCCGTTATCACAGCAGAACTCGAACCCGTCAGAGCACCATCGCTGATT GCTTTCTTTCACGGCCACTCTGCCTGTATGTGCCCGTTTCGATGGCGGTACAGCAGGTTTT CGCTCAAGCAACAGCGCATTCTGGCGCATGATCCGGTAAACACGTTTGGCATTGATCGCAG GCATACCATCAAGTTCTGCCTGTCTGCGAAGCAGCGCCCATACCCGACGATAACCATACGT TGGCAGCTCTCCGATAACATGGTGTATACGGAGAAGCACATCCGTATCATCAGTGTGACGA CTGCGGCGGCCATCCATCCAGTCATCGGTTCGTCTGAGAATGACGTGCAACTGCGCACGC GACACCCGGAGACAACGGCTGACTAAGCTTACTCCCCATCCCCGGGCAATAAGGGCGCGT |

| | |
|---|---|
| | GCGCTATCCACTTTTTTGCCCGTCCATATTCAACGGCTTCTTTGAGGAGTTCATTTTCCATC<br>GTTTTCTTGCCGAGCAGGCGCTGGAGTTCTTTAATCTGCTTCATGGCGGCAGCAAGTTCAG<br>AGGCAGGAACAACCTGTTCTCCGGCGGCGACAGCAGTAAGACTTCCTTCCTGGTATTGCTT<br>ACGCCAGAGAAATAACTGGCTGGCTGCTACACCATGTTGCCGGGCAACGAGGGAGACCGT<br>CATCCCCGGTTCAAAGCTCTGCTGAACAATTGCGATCTTTTCCTGTGTGGTACGCCGTCTG<br>CGTTTCTCCGGCCCTAAGACATCAATCATCTGTTCTCCAATGACTAGTCTAAAAACTAGTATT<br>AAGACTATCACTTATTTAAGTGATATTGGTTGTCTGGAGATTCAGGGGGCCAGTC–3’ |
| **SEQ1: Used for Multiplexing via *Pf*Ago and SpCas9n, toehold creation with *Pf*Ago** | 5’–GTGATAGTTAGCGATGCCGGGTTTGTATCGTGCCTTTTCATTTATTTCCAGCGGTATAGC<br>TGATAAATAATCCCTTAGTACATCAGGAACCTTATAAAAAAACCCGCGTTCTGCAAACCTTTC<br>TCGATAAAAGTGACAACCACTGAGTAAACAGATCAATGGAAGTAAATTCCAATTGTTTATTG<br>GATTTGCCCCTATATTTCCAGACATCTGTTATCACTTAACCCATTACAAGCCCGCTGCCGCA<br>GATATTCCCGTGGCGAGCGATAACCCAGCGCACTATGCGGATGCCATTCGTTATAATGCTC<br>GAACGCCTCTGCAAGGTTCTTTGCTGCCGTTAACCCGTCTGGTTTGGGCATGATACTGATG<br>TAGTCACGCTTTATCGTTTTCACGAAGCTCTCTGCTATTCCGTTACTCTCCGGACTCCGCAC<br>CGCCGTGTTCTTCGGTTC–3’ |
| **LGA (Register); Used for data LGA and LMI storage, sense-antisense nicking** | 5’–ACTCTGAATTTTTGCGGTGGTTTCAAGAAGGATTTTTTCTTTAAGAGAAACTATTTCTTTC<br>AATCCATCAGAAATTTTTCTTTTAATCACAACACCATTTGCAGGAGATTTTATCAAGCTCATC<br>TCTATTTTTTTTCGCATAAAATGTAATTCTCTCTAAAAGTGTTGATGCAAGTGGTAACGGCGA<br>TTGTATGTGGTCAACAAAATCAGGAAATAATAAATTAAATCGTTGCTGCATTGCCAATGGTTG<br>TTGCCTTATATAGGGTAACAACGTTGAAAAACCACATGTGATGATATTGTGATAGTTAGCGA<br>TGCCGGGTTTGTATCGTGCCTTTTCATTTATTTCCAGCGGTATAGCTGATAAATAATCCCTTA<br>GTACATCAGGAACCTTATAAAAAAACCCGCGTTCTGCAAACCTTTCTCGATAAAAGTGACAA<br>CCACTGAGTAAAC–3’ |
| **Orthogonal Sequence 1** | 5’–GGATGGTAGAACGCCATCACCTTTGAGTAATCCATGTTTTTATTTTTCTCATGGATTGTGC<br>CTTTTAGGTTTTCCAGTGACCTGGTACAGTGTTGAACACCACAGGCAAAACCATAATGCGTT<br>GCCAGTGCCTTTTCTTCTTCGTTATTCCAGTAGTAGTAAAGCGATATTGTGCCGGTAAATTT<br>CTTTTTCCCCGCATTTAAAACAAATCTTGCGTCATCGGCAACCTGAATATTGTGCCGGTTAA<br>GTGCTGGATCATTGAGCATCTTAACGATGTCATCGGCACCATCAGTGATAAGGTAATCCAG<br>CGATTCCCCTACAAAAACATAGCCTTTTGTTCCATTACTATCTTTTGCCTGCGATAGCCCAGT<br>GATGGTGTCTGACTGCGATTGATACTTTCCTGAACTACCCTTTGCGGTATGGGAAACCAGA<br>GCATACGAGATACAGCCT–3’ |
| **Orthogonal Sequence 2:** | 5’–AGAACAATTGCCGAAGATTATTGATGCGCTCTCACCGCAAGGTGAAGTGTCACCACAAG<br>CCAATAACGATCTGCTTTCCGCAGGCATGGAACTGCTGAAAGGGAAACTCTTCCGCTAAGC<br>AAAAAGGGAGCACGCAACGCGCGTTGCATCCCCCTTTTCCCTCGGGTGTTGTACTGATTTT<br>TGAGCGGAATCGCGTTAGCATGGGTCAGGAACCAATCTACCTGGGGAACTCATGGCTATTA<br>AACTCATTGCTATCGATATGGATGGCACCCTTCTGCTGCCCGATCACACCATTTCACCCGC<br>CGTTAAAAATGCGATTGCCGCAGCTCGCGCCCGTGGCGTGAATGTCGTGCTAACGACGGG<br>TCGCCCGTATGCAGGTGTGCACAACTACCTGAAAGAGCTGCATATGGAACAGCCGGGCGA<br>CTACTGCATTACTTATAACGGCGCGCT–3’ |
| **Orthogonal Sequence 3:** | 5’–AAACGCCAGTGGTAATGGCAGGAATGGTTGGCAGCAACGTCGGCTGGAAAGTTGCACC<br>GTATTTATCTGTTCCTGCCTGTTTTTCGTCTATTGGCGAACAATTAACGTCAGTTGGCGACA<br>ATATCTGGATTATTCCCGGATTATGTGTCTCTCATGACGATAACCACAATGTGATGCGCGGC<br>GAAGAAACACAATTGATCGGCGCGCGAGCTCTGGCTCCTTCCTCTCTTTATGTCATGCCCG<br>GAACCCATTGCAAATGGGTGCAGGCCGATAGCCAGCAAATCAACGATTTTCGCACCGTGAT<br>GACCGGTGAATTACATCATTTACTGTTAAATCACTCATTGATTGGCGCAGGTTTGCCGCCGC<br>AGGAAAACTCTGCCGATGCCTTCACAGCTGGCCTTGAGCGTGGTCTTAATACGCCCGCCAT<br>ATTGCCGCAGCTTTTTGAAGTTC–3’ |
| **Orthogonal Sequence 4:** | 5’–AACGTTGAACTGGTAGACGAAAACCATTACCGCATTGCTATCGCTGTGGCTGGTTTTGCT<br>GAGAGCGAACTGGAAATTACCGCCCAGGATAATCTGCTGGTGGTGAAAGGTGCTCACGCC<br>GACGAACAAAAAGAGCGCACCTATCTGTACCAGGGCATCGCTGAACGCAACTTTGAACGCA<br>AATTCCAGTTAGCTGAGAACATTCATGTTCGTGGTGCTAACCTGGTAAATGGTTTGCTGTAT<br>ATCGATCTCGAACGCGTGATTCCGGAAGCGAAAAAACCGCGCCGTATCGAAATCAACTAAT<br>TCCCTAAGGCCGCCTGGCGCGGCCTGACATCTCCATGCTCGCCGTCAGGGAGCATATGCG<br>AATCTTCGGATTTGCAGGTACTTACTCGCTTCTTAGAAGGAGAAATGACTATGCGTAACTTC<br>GATTTATCCCCACTGATGCGTCAA–3’ |

**Table S5. List of oligos used in this work.**

| Primers | |
|---|---|
| **Primary 1kb sequence** | |
| KT_PS1_PF | GCGAACAGCAGCGGTTATG |
| KT_PS1_PR | ACGCCATACCAACCGAACAG |
| **Primary 1.2kb sequence** | |
| KT_PS1.2_PF | AGCGAATGCAACGGGTGATG |
| KT_PS1.2_PR | TGCAGGTGGAATGATGGGGG |
| **Primary 1.5kb sequence** | |
| KT_PS1.5_PF | CGTTGCTGCATTGCCAATGG |
| KT_PS1.5_PR | GACTGGCCCCCTGAATCTC |
| **SEQ1 primers** | |
| KT_TN_PF | GTGATAGTTAGCGATGCCGGG |
| KT_TN_PR | GAACCGAAGAACACGGCGG |
| **LGA Primers** | |
| KT_GA_PF | ACTCTGAATTTTTGCGGTGGTTTCAAG |
| KT_GA_PR | GTTTACTCAGTGGTTGTCACTTTTATCGAGAAAG |
| **Orthogonal sequences primers** | |
| Ortho1-PF | GGATGGTAGAACGCCATCACCT |
| Ortho1-PR | AGGCTGTATCTCGTATGCTCTGG |
| Ortho2-PF | AGAACAATTGCCGAAGATTATTGATGCG |
| Ortho2-PR | AGCGCGCCGTTATAAGTAATGC |
| Ortho3-PF | AAACGCCAGTGGTAATGGCAG |
| Ortho3-PR | GAACTTCAAAAAGCTGCGGCAATATGG |
| Ortho4-PF | AACGTTGAACTGGTAGACGAAAACCA |
| Ortho4-PR | TTGACGCATCAGTGGGGATAAATCG |
| **Guides** | |
| **Primary nicking verification (1kb)** | |
| KT_G1 | /5Phos/CAATAAATCACACTGC |
| **Primary verification (1.5kb)** | |
| KT_G2 | /5Phos/TGCTGTGATCGTGAGG |
| **Primary verification (1.2kb)** | |
| KT_G3 | /5Phos/TGAGTCTATGAGTTTC |
| **Triple Nicking (I) gDNAs** | |
| KT_TN1_G1 | /5Phos/ATCACAACACCATTTG |
| KT_TN1_G2 | /5Phos/TGATGCAAGTGGTAAC |
| KT_TN1_G3 | /5Phos/AAATCGTTGCTGCATT |
| **Triple Nicking (I) crRNAs** | |
| KT_TN1_gCas1 | /AlTR1/rUrUrArArUrCrArCrArArCrArCrCrArUrUrUrGrCrGrUrUrUrArGrArGrCrUrArUrGrCrU/AlTR2/ |

| | |
|---|---|
| KT_TN1_gCas2 | /AlTR1/rArGrUrGrUrUrGrArUrGrCrArArGrUrGrGrUrArArGrUrUrUrArGrArGrCrUrArUrGrCrU/AlTR2/ |
| KT_TN1_gCas3 | /AlTR1/rArArUrCrGrUrUrGrCrUrGrCrArUrUrGrCrCrArArGrUrUrUrArGrArGrCrUrArUrGrCrU/AlTR2/ |
| **Triple Nicking (II) gDNAs** | |
| KT_TN2_G1 | /5Phos/ATTGGAATTTACTTCC |
| KT_TN2_G2 | /5Phos/ATGGGTTAAGTGATAA |
| KT_TN2_G3 | /5Phos/AGCAAAGAACCTTGCA |
| **Triple Nicking (II) crRNAs** | |
| KT_TN2_gCas1 | /AlTR1/rCrCrArUrUrGrArUrCrUrGrUrUrArCrUrCrArGrGrUrUrUrArGrArGrCrUrArUrGrCrU/AlTR2/ |
| KT_TN2_gCas2 | /AlTR1/rGrUrUrArArGrUrGrArUrArArCrArGrArUrGrUrCrGrUrUrUrArGrArGrCrUrArUrGrCrU/AlTR2/ |
| KT_TN2_gCas3 | /AlTR1/rCrGrUrUrCrGrArGrCrArUrUrArUrArArCrGrArArGrUrUrUrArGrArGrCrUrArUrGrCrU/AlTR2/ |
| **Gettysburg Address and Lincoln's memorial** | |
| KT_LGA_G1 | /5Phos/TTCAATCCATCAGAAA |
| KT_LGA_G2 | /5Phos/ATCACAACACCATTTG |
| KT_LGA_G3 | /5Phos/TATCAAGCTCATCTCT |
| KT_LGA_G4 | /5Phos/AAGTGTTGATGCAAGT |
| KT_LGA_G5 | /5Phos/GTATGTGGTCAACAAA |
| KT_LGA_G6 | /5Phos/AAATCGTTGCTGCATT |
| KT_LGA_G7 | /5Phos/GGTAACAACGTTGAAA |
| KT_LGA_G8 | /5Phos/TATTGTGATAGTTAGC |
| KT_LGA_G9 | /5Phos/TTAGTA CATCAGGAAC |
| KT_LGA_G10 | /5Phos/TTCTGCAAACCTTTCT |
| **Orthogonal Sequence 1** | |
| KT_OS1_G1 | /5Phos/TTAGGTTTTCCAGTGA |
| KT_OS1_G2 | /5Phos/TCTTCTTCGTTATTCC |
| KT_OS1_G3 | /5Phos/AAGTGCTGGATCATTG |
| KT_OS1_G4 | /5Phos/CAGTGATAAGGTAATC |
| KT_OS1_G5 | /5Phos/TGATACTTTCCTGAAC |
| **Orthogonal Sequence 2** | |
| KT_OS2_G1 | /5Phos/TGAAGTGTCACCACAA |
| KT_OS2_G2 | /5Phos/ATGGAACTGCTGAAAG |
| KT_OS2_G3 | /5Phos/TCAGGAACCAATCTAC |
| KT_OS2_G4 | /5Phos/ATTGCTATCGATATGG |
| KT_OS2_G5 | /5Phos/TGAATGTCGTGCTAAC |
| KT_OS2_G6 | /5Phos/AAGAGCTGCATATGGA |
| KT_OS2_G7 | /5Phos/CGATCACACCATTTCA |
| **Orthogonal Sequence 3** | |

| | |
|---|---|
| KT_OS3_G1 | /5Phos/AACAATTAACGTCAGT |
| KT_OS3_G2 | /5Phos/ATGTGTCTCTCATGAC |
| KT_OS3_G3 | /5Phos/TTCCTCTCTTTATGTC |
| KT_OS3_G1 | /5Phos/CAGCAAATCAACGATT |
| KT_OS3_G4 | /5Phos/CGAAGAAACACAATTG |
| **Orthogonal Sequence 4** | |
| KT_OS4_G1 | /5Phos/ATAATCTGCTGGTGGT |
| KT_OS4_G2 | /5Phos/AAATTCCAGTTAGCTG |
| KT_OS4_G3 | /5Phos/TGGTAAATGGTTTGCT |
| KT_OS4_G4 | /5Phos/TTTGCAGGTACTTACT |
| KT_OS4_G5 | /5Phos/AGGAGAAATGACTATG |
| **Sense and anti-sense nicking** | |
| KT_SA_G1B | /5Phos/TTCAATCCATCAGAAA |
| KT_SA_G2B | /5Phos/AAGTGTTGATGCAAGT |
| KT_SA_G3B | /5Phos/AAATCGTTGCTGCATT |
| KT_SA_G4B | /5Phos/TATTGTGATAGTTAGC |
| KT_SA_G1T | /5Phos/AATCCTTCTTGAAACC |
| KT_SA_G2T | /5Phos/GAGATGAGCTTGATAA |
| KT_SA_G3T | /5Phos/GTTGACCACATACAAT |
| KT_SA_G4T | /5Phos/TCATCACATGTGGTTT |
| KT_SA_G5T | /5Phos/TAAGGTTCCTGATGTA |
| KT_SA_G6T | /5Phos/TCGAGAAAGGTTTGCA |
| **Non-destructive toehold detection** | |
| Reporter | /5IABkFQ/GAG CTT GAT AAA ATG AGA C/36-FAM/ |
| Reporter* | GTC TCA TTT TAT CAA GCT C |

#### D. References


1. J.H. Conway, N.J.A. Sloane. *Designs, codes and Cryptography*, 4 (1), 31-42 (1994)
2. S.W. McLaughlin et al. *Proceedings of 1995 IEEE International Symposium on Information Theory*, (2002).
3. J. T. Robinson, H. Thorvaldsdóttir, W. Winckler, M. Guttman, E.S. Lander, G. Getz, J. P. Mesirov. *Nat Biotechnol*., 29, 24–26 (2011)
4. Integrated DNA Technologies. Alt-R CRISPR-Cas9 System: In vitro cleavage of target DNA with ribonucleoprotein complex. [Online] Coralville, *Integrated DNA Technologies* (2017)
5. S.H.T. Yazdi, Y. Yuan, J. Ma, H. Zhao, O. Milenkovic. *Sci. Rep*. 5, 14138 (2015).
6. K. Liu, C. Pan, A. Kuhn, A.P. Nievergelt, G. Fantner, O. Milenkovic, A. Radenovic, Detecting topological variations of DNA at single-molecule level, *Nature Communications*, 10, 3 (2019).
7. S.H.T. Yazdi, R. Gabrys, O. Milenkovic. *Scientific reports* 7.1 (2017).
8. R.N. Grass, R. Heckel, M. Puddu, D. Paunescu, W. Stark. *J. Angew. Chem. Int. Ed*. 54, 2552–2555 (2015).
9. G. M.Church, Y.Gao, S. Kosuri. *Science* 337, 1628-1628 (2012).
10. Y. Erlich, D.  Zielinski, D. *Science*, 355, 950-954 (2017).
11. B. Lindström et al., J. N. Srivastava, ed., A survey of Statistical Design and Linear Models, North-Holland Publishing Company, (1975).
12. A. Emad, O. Milenkovic, *IEEE Transactions on Information Theory* 60, 8, 4614-4636, (2014).
13. B. Wang, C. Chalk, D. Soloveichik*, DNA 25 Conference*, Seattle, WA, U.S.A. (2019).
14. T. Chen, M. Riedel, *11th International Workshop on Bio-Design Automation (IWBDA)*, Cambridge, England, U.K. (2019).
